# Trifluoroacetate reduces plasma lipid levels and the development of atherosclerosis in mice

**DOI:** 10.1101/2025.03.06.641713

**Authors:** Wei Tang, Audrey S. Black, Romana Moench, Katayoon Marzban, Juan Antonio Raygoza Garay, James J. Zheng, Louis Conway, Antonio F. M. Pinto, Christopher G. Parker, Alan Saghatelian, Luke J. Leman, M. Reza Ghadiri

## Abstract

Trifluoroacetate (TFA) has been assumed to be an innocuous counterion (to cationic amino acid side chains) present in countless synthetic bioactive peptides and a few FDA-approved therapeutics. We show here that TFA is in fact bioactive and causes dramatic biological effects in multiple strains of mice and cultured human and rat liver cells. In high-fat diet (HFD)-fed low-density lipoprotein receptor-null (LDLr^-/-^) mice, TFA reduces the levels of plasma cholesterol, triglycerides, and the development of atherosclerotic lesions following either oral or intraperitoneal administration. These physiological effects were observed with TFA alone, or with TFA present as a counterion of a variety of short, unrelated synthetic peptide sequences. Mechanistic investigations including RNA-seq, confocal microscopy, western blotting, metabolomics, proteomics, pharmacokinetics, and biochemical assays indicated that TFA induces peroxisome proliferation by activating peroxisome proliferator-activated receptor (PPAR)-alpha. We confirmed that TFA also caused peroxisome proliferation and downstream phenotypic effects in cultured human and rat liver cells, wild-type C57/Bl mice, and apolipoprotein E-null (apoE^-/-^) mice, leading to anti-atherosclerotic effects in the latter strain. Given that TFA is a counterion in many peptides employed in early research and development settings, these findings raise the possibility that TFA may be confounding or contributing to phenotypic changes observed in many studies involving peptides. Although our studies suggest that TFA or its analogues might have therapeutic applications, it should be noted that TFA is also a persistent environmental contaminant that is found at high levels in humans relative to other polyfluoroalkyl substances (PFAS), and is a major metabolite following treatment of patients with common inhaled anesthetics, suggesting that the biological effects reported here could have other implications for human health.

## Introduction

Atherosclerosis remains one of the leading causes of death in the world, despite the widespread use of statins and newer drugs for lowering lipid levels. Following the realization that certain endogenous apolipoproteins (e.g., apoA-I and apoE) are protective against the development of atherosclerosis and other metabolic diseases, intense effort has been invested in developing short peptides that mimic the cardioprotective properties of native apolipoproteins as potential therapeutics [1–3]. Numerous, mostly helical, synthetic apolipoprotein mimetic peptide sequences have been characterized in vitro and in vivo, with at least six distinct peptides (L-4F, D-4F, ETC-642, 5A, CN-105, AEM28) having progressed as far as human clinical trials [3]. The various apolipoprotein mimetic peptides that have been studied exert a range of biological effects, such as cholesterol efflux from cells, binding of pro-inflammatory oxidized lipids, lowering of plasma cholesterol and/or triglyceride levels, and reducing the development of atherosclerotic lesions. An important unresolved challenge in apolipoprotein mimetic peptide research is the discordance sometimes encountered between in vitro properties of experimental peptides and their corresponding activities in vivo (for example, see [4]).

Peptides are typically prepared and employed as trifluoroacetate (TFA) salts in preclinical stages of research. TFA is a reagent during solid-phase peptide synthesis (SPPS), and a component in buffer systems typically used for reversed-phase high-performance liquid chromatography (HPLC) purification of peptides, yielding peptides as TFA salts following purification. Ion pairing between TFA and cationic moieties in peptides (Lys, Arg, His, free N-terminus) is persistent, requiring a dedicated ion exchange step to remove TFA from the purified compounds [5]. A given peptide molecule will contain varying numbers of associated molecules of TFA, depending on the number of cationic moieties present in the sequence. TFA is a counterion in at least two FDA-approved peptide drugs, bivalirudin and corticorelin [5]. Although the presence of TFA counterions is often overlooked, some previous reports have hinted that TFA can affect certain biological pathways. For instance, at relatively high concentrations (millimolar range), TFA can activate the lactate receptor hydroxycarboxylic acid receptor 1 (HCA1) [6], free fatty acid receptor 2 (FFA2) [7], ATP-sensitive potassium channels [8], and TFA can act as an allosteric modulator of the glycine receptor [9]. Also, modestly different antibacterial activities were reported for alkylguanidino urea antibacterial compounds when the counterion was switched from TFA to Cl^-^ [10]. Trifluoroacetate was reported to inhibit proliferation of osteoblasts and chondrocytes in cell culture [11]. Similarly, calcium trifluoroacetate was reported to prevent angiogenesis, although the proposed mechanism was not related to TFA but rather to the alteration of cytosolic calcium levels and the prevention of calcium signaling by VEGF [12].

In the course of our efforts to develop new apolipoprotein mimetic peptides, we were faced with a perplexing observation– several short, unrelated peptide sequences intended as negative controls all provoked a similar phenotypic effect in vivo (reductions in plasma cholesterol levels in a mouse model of atherosclerosis). This was especially surprising given that the peptides were administered orally, and had been expected to undergo rapid digestion in vivo, which would preclude any biological effects. Eventually, we made the remarkable discovery that the observed reductions in plasma cholesterol were not directly related to the peptides themselves, but rather stemmed from TFA that was present with the peptides as counterions. Expanding further, we found that TFA can reduce the development of atherosclerosis in the two most common mouse models of atherosclerosis (LDLr^-/-^ and apoE^-/-^ mice) and can also provoke mechanistically-related phenotypic effects in wild-type C57Bl mice. Based on transcriptomics analysis of tissue from TFA-treated animals, along with both cell culture and in vivo experiments, we established that the mechanism of the observed effects is due (at least in part) to TFA-induced proliferation of peroxisomes. Because most synthetic peptides in preclinical stages of research contain TFA counterions, the discovery that TFA impacts cellular metabolism in vitro and in vivo means that TFA may have played a potentially confounding role in numerous biological studies in which synthetic peptides have been employed. Moreover, the biological effects of TFA reported here may be relevant to human health because TFA is highly persistent and environmental concentrations of TFA are increasing [13–17], resulting in high levels of TFA in humans relative to other polyfluoroalkyl substances (PFAS) [18]. TFA is also a major metabolite of common inhaled anesthetics, leading to high plasma TFA levels in human patients for days following anesthetic administration [19–21].

## Results

### Discovery that TFA reduces plasma cholesterol levels in LDLr^-/-^ mice

Our laboratory previously advanced a series of in vitro and in vivo functional apoA-I mimetic peptide constructs composed of multiple copies of a 23-residue amphiphilic helical peptide attached to a small molecule scaffold [22,23]. The impetus for this work was our hypothesis that multimeric constructs, by virtue of their multiple interacting amphiphilic peptide helices, might better recapitulate the scaffolding and HDL remodeling properties of full-length native apoA-I (which contains 11 helical segments) versus a monomeric helical peptide. Indeed, we observed that a trimeric construct was more efficient in promoting in vitro cholesterol efflux than the corresponding monomer peptide, as well as possessing greater stability to proteolytic degradation in vitro and a longer half-life in plasma following intraperitoneal (i.p.) administration [23].

Whilst these constructs displayed good plasma half-lives, significant HDL remodeling activities, and atheroprotective effects when administered via i.p. injection to high-fat diet (HFD, 1.25% cholesterol, 15.8% fat, and no cholate)-fed LDLr^-/-^ mice, we were surprised that the peptide constructs were also remarkably active when administered orally, despite having undetectable plasma concentrations (lack of oral bioavailability) [22]. Even more puzzling, the 23-residue monomer peptide, included in the 10-week in vivo studies intended as a negative control, exhibited similar oral activity profiles including a marked reduction in total plasma cholesterol levels and atherosclerotic plaques [22]. Considering that linear peptides made up of natural L-amino acids should undergo rapid digestion in the gut, we were left with the possibility that a shorter peptide fragment could be the active species acting on targets in the gut.

To test this idea, a set of four shorter peptide fragments derived from the monomer were synthesized (Figure 1a). The sequences of these peptides were based on results from in vitro pepsin digestions of the parent 23-residue peptide (Figure S2). These fragment peptides ranged from four to eight residues in length and, together, spanned the full length of the parent peptide sequence (Figure 1a). We also synthesized two additional previously reported tetrapeptides unrelated to the monomer peptide, one of which (FREL) was reported to prevent the development of atherosclerosis in vivo, while the other of which (KERS) was reported to be ineffective [24]. Each of these six short peptides was individually administered orally to high-fat fed LDLr^-/-^ mice for two weeks at a 90 mg/kg/day dose (see Figure S1 for dosing schedules). Astoundingly, every one of these peptides caused a statistically significant reduction in plasma cholesterol levels, ranging from 24–55% lower than that of vehicle-treated animals (Figure 1b). The basis for this effect was perplexing. The peptides were too short to fold, were not expected to self-assemble, and lacked any sequence homology. We concluded that the only commonality among the short peptide fragments in question must be the presence of TFA counterions (present following standard HPLC purification as the counterion of cationic groups on the peptides). To test if the TFA counterions were somehow causing the observed phenotypic changes, one of the short peptides (hexamer LSALEE) was converted from the TFA salt to the chloride salt [25]; ^19^F-NMR confirmed the complete removal of TFA from the peptide (Figure S3). In contrast to the TFA salt of the peptide, the LSALEE peptide administered as the chloride salt completely lost its ability to lower plasma cholesterol levels (Figure 1b). This finding implicated TFA, albeit indirectly, as the active species causing reduced plasma cholesterol levels.

**Figure 1.**
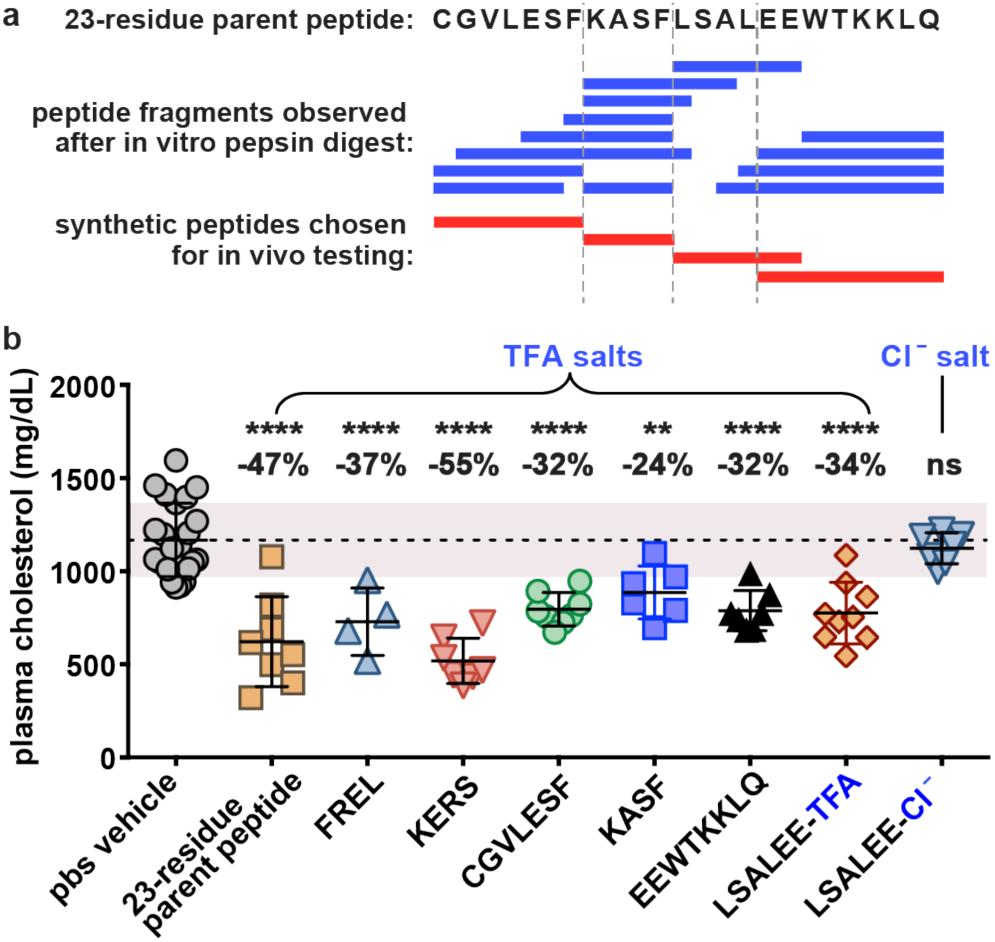
Peptide-TFA salts reduced plasma cholesterol levels in HFD-fed LDLr^-/-^ mice, whereas a corresponding peptide-Cl^-^ salt did not. **a**) In vitro pepsin digestions of the parent 23-residue linear apoA-I mimetic peptide yielded a number of peptide fragments (blue bars). Based on the observed fragments, four short synthetic peptides were chosen for in vivo testing (red bars). **b**) In vivo studies with TFA or Cl^-^-salts of the short linear peptides led us to the discovery that removal of the TFA counterion abolished the observed plasma cholesterol reductions in vivo. Peptides were administered to female HFD-fed LDLr^-/-^ mice *ad libitum* in the drinking water for 2 weeks (n=4-9 per group; 75 mg/kg/day = 28 μmol peptide/kg/day for 23-residue parent peptide; 90 mg/kg/day = ∼130 μmol peptide/kg/day for shorter peptides). All peptides were present as the TFA salt, except LSALEE-Cl^-^. Scatter plots are shown with mean ± SD. The shaded region indicates one standard deviation above and below the mean (dotted line) for the PBS vehicle group (n=23). *p* values were determined by one-way ANOVA, post hoc Tukey-Kramer test; ns, not significantly different; **, *p* < 0.01; ****, *p* < 0.0001.

### TFA mediates dose-dependent cholesterol-lowering effects regardless of the route of administration

A series of dosing studies (see Figure S1 for dosing schedules) unambiguously established that TFA was responsible for the observed reductions in plasma cholesterol. A dose-dependent reduction in cholesterol levels was observed following two-week daily TFA administration in the context of either i.p. injection or oral gavage using female LDLr^-/-^ mice (Figure 2a). Plasma cholesterol levels were not significantly affected by TFA treatment at 22 µmol/kg/day, but began trending lower at doses of 67 µmol/kg/day and higher. Likewise, a two-week daily dose of 200 µmol/kg/day TFA administered *ad libitum* in the drinking water, this time using male LDLr^-/-^ mice, caused a significant reduction in plasma cholesterol. In contrast, an analogous 200 µmol/kg/day dose of acetate included as a negative control did not affect plasma cholesterol levels (Figure 2a). The overall magnitude of plasma cholesterol reduction was similar for the 200 µmol/kg/day dose of TFA (in the range of 25–35% compared to vehicle-treated control animals), regardless of whether the TFA was administered *ad libitum* in the drinking water, by once-daily i.p. injection, or by once-daily oral gavage (Figure 2a). Notably, the 200 µmol/kg/day TFA dose was chosen to be near the middle of the range of TFA doses that were present in the earlier studies as counterion with the fragment peptides (depending on the number of cationic moieties in the particular peptide sequence, the expected amount of TFA administered as peptide counterion in those studies ranged from 130–390 µmol/kg/day). Altogether, these data indicated that TFA treatment produces dose-dependent lipid-lowering effects in male or female HFD-fed LDLr^-/-^ mice regardless of the route of administration, which begin to be observable at doses around 67 µmol/kg/day.

**Figure 2.**
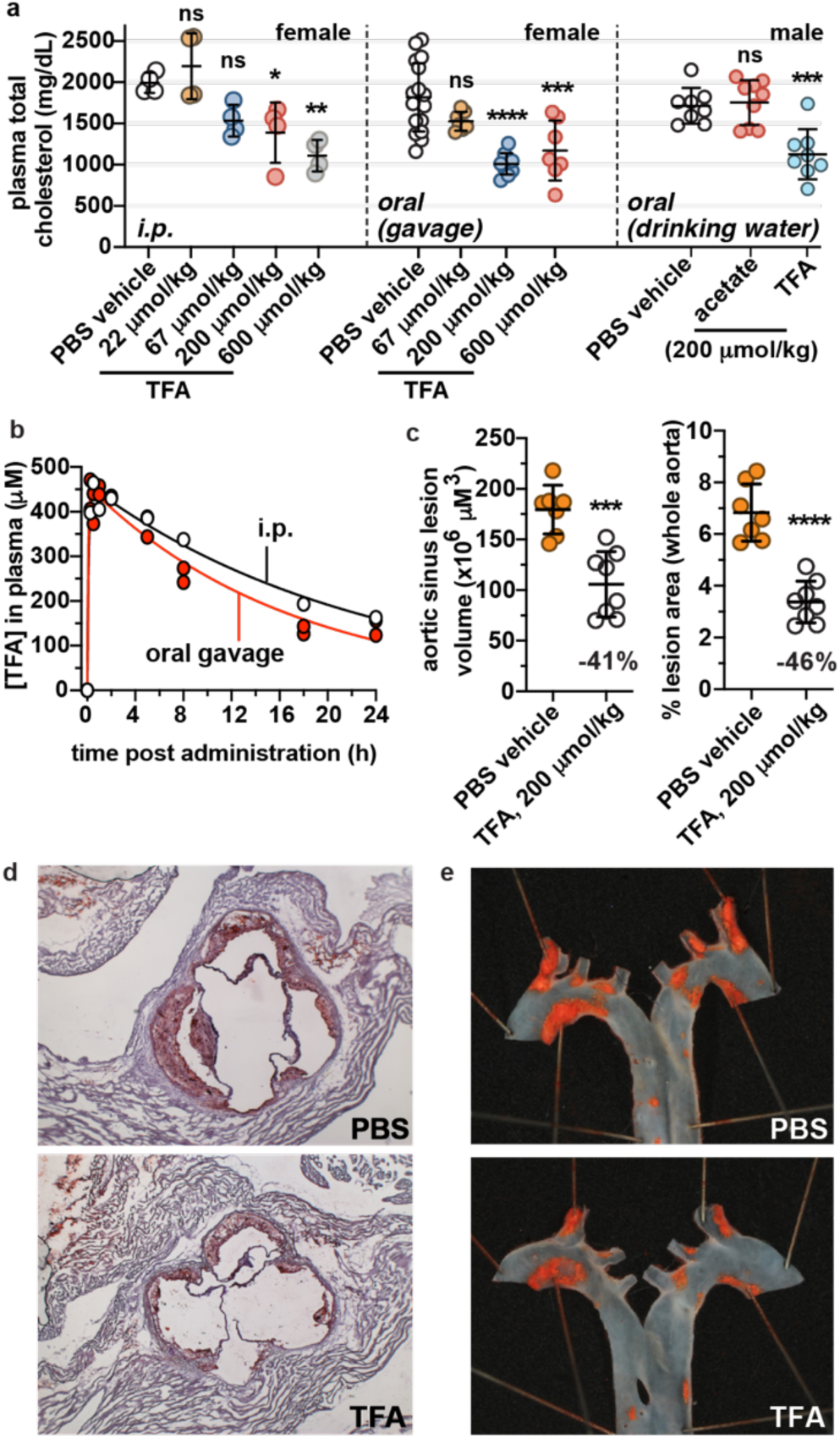
TFA exerts dose-dependent cholesterol-lowering effects, has a long in vivo half-life, and reduces the development of atherosclerosis in HFD-fed LDLr^-/-^ mice. **a**) TFA reduced plasma total cholesterol levels of female HFD-fed LDLr^-/-^ mice in a dose-dependent fashion following two-week administration by daily i.p. injection (n=4/group) or daily oral gavage (n=7-17/group). Cholesterol levels were also reduced in male HFD-fed LDLr^-/-^ mice by TFA (n=8) compared to PBS vehicle (n=8) following two-week administration *ad libitum* in the drinking water (200 μmol/kg/day). Sodium acetate (n=9) served as a negative control. Scatter plots are shown as mean ± SD. *p* values were determined by one-way ANOVA comparing each experimental group to the corresponding PBS vehicle control, post hoc Tukey-Kramer test; ns, not significantly different; *, *p* < 0.05; **, *p* < 0.01; ***, *p* < 0.001; ****, *p* < 0.0001. **b**) Pharmacokinetic traces of TFA in plasma following oral gavage or i.p. administration of a 200 μmol/kg dose to LDLr^-/-^ mice. For each timepoint, samples were taken from n=2 animals. The curves represent a fit of the data to a pharmacokinetic equation for single dose plasma concentration. **c**) TFA (200 μmol/kg/day, n=8) strikingly reduced the development of atherosclerotic lesions in HFD-fed female LDLr^-/-^ mice compared to PBS vehicle (n=8) following ten-week administration *ad libitum* in the drinking water. *p* values were determined by Student’s t-test. Representative images of aortic sinus cross-section (**d**) and aorta (**e**) are shown, with lipid stained red.

### TFA exhibits a relatively long in vivo half-life

To measure the plasma concentration and half-life of TFA following i.p. or oral gavage administration to mice, a ^19^F-NMR-based assay was developed (Figure S4). Groups of mice (n=2/group) were given a 200 µmol/kg dose of TFA, and blood was drawn from the animals at various time points. The plasma was diluted with water and scanned by ^19^F-NMR along with a coaxial tube insert containing trifluoromethoxybenzene as an internal concentration standard. Based on this analysis, we found that plasma concentrations of TFA were very similar for either oral or i.p. routes of administration (similar exposure AUC) (Figure 2b). For both routes of administration, the 200 µmol/kg dose of TFA resulted in a maximal plasma TFA concentration of ∼450 µM around 15 minutes post-administration, which slowly decreased to ∼150 µM at 24 hours post-administration (Figure 2b). The observed long half-life in plasma is presumably due to TFA binding/ion exchange to plasma proteins. Thus, given the long plasma half-life and daily dosing regimen, the plasma concentration of TFA likely would have remained well above 150 µM for the entire duration of our studies involving i.p. or oral gavage administration of TFA.

### TFA reduces the development of atherosclerosis in HFD-fed LDLr^-/-^ mice

Given the key contributing role that high plasma cholesterol levels play in the development of atherosclerosis, the observation of TFA-mediated reductions in plasma cholesterol suggested that TFA might have the potential to reduce the development of atherosclerotic lesions in female LDLr^-/-^ mice. To assess this possibility, we carried out a long-term atheroprotection study. The mice were fed a standard chow diet until ∼10 weeks of age and were then switched to the HFD along with commencement of PBS vehicle (n=8) or TFA in PBS (200 µmol/kg/day, n=8) as the drinking water (see Figure S1 for dosing schedules). As had been observed in the shorter duration studies, plasma cholesterol levels were reduced by two weeks of TFA treatment (29% reduction; p=0.00005, Figure S5), and remained lower in the TFA-treated mice at the 6-week (34% reduction, p=0.0015, Figure S5) and 10-week timepoints (28% reduction, p=0.0010, Figure S5). Following the 10-wk study, we observed a remarkable reduction in the volumes of atherosclerotic lesions in the hearts of the mice (41% reduction) (Figures 2c, S6). Likewise, a greatly reduced plaque area was observed in en face aortic sections (46% reduction) (Figures 2c, S7). Consistent with the reduced development of atherosclerosis, TFA treatment reduced the circulating levels of serum amyloid A (SAA), a marker of systemic inflammation (Figure S8). In a follow-up 8-wk atheroprotection study using a lower dose of TFA (10 µmol/kg/day) with an i.p. route of administration, TFA treatment did not cause changes in plasma cholesterol or triglyceride levels over the course of the study, nor did it reduce the development of aortic lesions (Figure S9).

Overall, including various studies described herein to characterize TFA dose dependency, route of administration, mechanism of action, and TFA analogs (*vide infra*), we have replicated the cholesterol-lowering effect promoted by TFA in LDLr^-/-^ mice in eight independent two-week vehicle-controlled trials using an oral route of administration, and three independent two-week vehicle-controlled trial using i.p. administration. These findings convincingly support that TFA, at concentrations that could be present with experimental peptides as counterion, can reduce plasma lipid levels and the development of atherosclerosis in a leading mouse model of the disease (LDLr^-/-^ mice). TFA treatment did not cause any signs of toxicity or distress in LDLr^-/-^, apoE^-/-^, or wild-type C57Bl animals. There were no differences in weight gain for TFA-treated mice compared to vehicle-treated controls over the course of multi-week studies (Figure S10), nor were there changes in the plasma levels of liver enzymes alanine transaminase (ALT) or aspartate aminotransferase (AST) (Figure S10), which are standard markers of liver damage or disease.

### TFA upregulates genes involved in fatty acid metabolism and acyl-CoA biochemistry

RNA-seq analysis of liver tissue from TFA-treated (i.p. administration) HFD-fed LDLr^-/-^ mice was carried out to explore possible mechanistic underpinnings of the observed effects. The analysis revealed dose-dependent changes in gene expression; principal component analysis of the data indicated that TFA doses of 67 µmol/kg, 200 µmol/kg, and 600 µmol/kg each had distinct gene expression profiles compared to the vehicle control and 22 µmol/kg TFA groups (Figure 3a). At the highest dose of TFA (600 µmol/kg), gene expression was altered for 686 genes with an adjusted p-value <0.05 (Spreadsheet S1). Pathway enrichment analysis of the gene expression data indicated that the pathways most upregulated by TFA treatment were related to the metabolism of fatty acids and peroxisome proliferation (Figure 3a, Spreadsheet S1). Many of the upregulated genes (e.g., enoyl-CoA hydratase and 3-hydroxyacyl CoA dehydrogenase, EHHADH; acyl-CoA oxidase 1, ACOX1) encode enzymes specific to peroxisomal beta-oxidation of fatty acids, while others encode for enzymes outside of the peroxisome (e.g., acyl-CoA thioesterase 1, ACOT1; acyl-CoA thioesterase 2, ACOT2) that are nonetheless target genes of the transcription factor peroxisome proliferator activated receptor (PPAR)-α and are upregulated by peroxisome proliferating agents [26]. Numerous upregulated enzymes were involved in processing of various acyl-coenzyme A (CoA) species, which would be consistent with increased conversion of fatty acids to acyl-CoA upon peroxisome proliferation. However, we also considered that changes in the expression of enzymes involved in CoA biochemistry could reflect the incorporation/participation of TFA as a structural analog of acetate in metabolism. Therefore, based on the biological pathways and enzymes implicated by the transcriptomics analysis, we experimentally tested two main mechanistic possibilities for the action of TFA: (a) TFA treatment induced peroxisome proliferation, leading to shifts in lipid metabolism including enhanced fatty acid oxidation and decreased lipogenesis; and (b) TFA impacted metabolism by generating trifluoroacetylated metabolites and/or proteins, possibly through the intermediacy of trifluoroacetyl-CoA (analog of acetyl-CoA).

**Figure 3.**
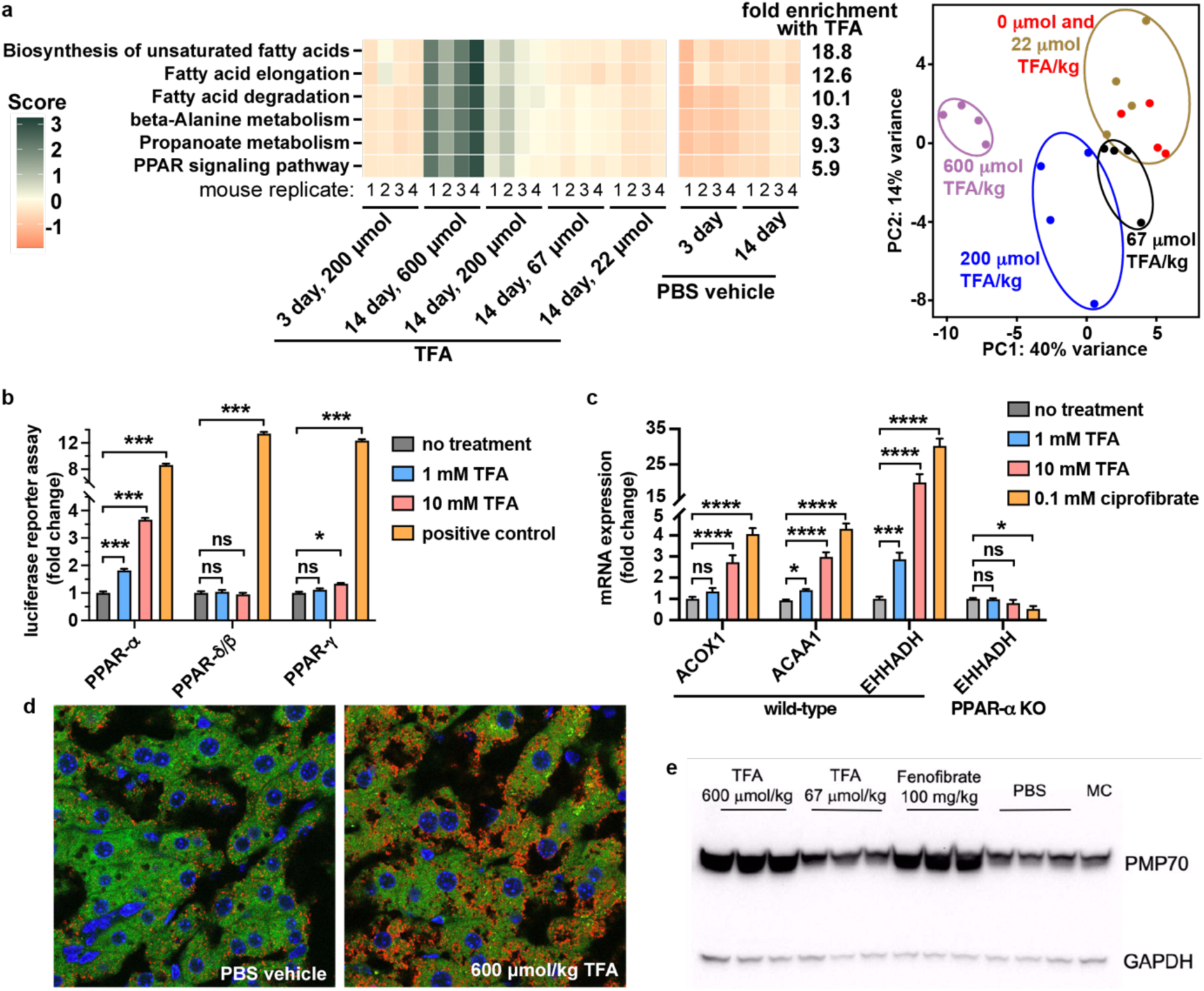
Gene and protein expression changes indicate that TFA stimulates peroxisome proliferation in mice and cultured liver cells. **a**) Left, heatmap showing the top biological pathways from pathway enrichment analysis of gene expression data from liver tissue of LDLr^-/-^ mice following three day or two-week TFA treatment (i.p.). Right, principal component analysis of the gene expression data. **b**) Reporter assay in human HepG2 liver cells transfected with different mouse PPAR isoforms, in which luciferase expression is driven by activation of the PPAR response element. A different positive control was used for each PPAR isoform: 0.1 mM ciprofibrate for PPAR-α, 1 µM GW501516 for PPAR-δ/β, and 10 µM rosiglitazone for PPAR-γ. **c**) Expression of several PPAR target genes in rat FAO liver cells was increased by TFA treatment, as indicated by qPCR. The effect was lost in PPAR-α knockout FAO cells. **d**) Confocal microscopy at 60x magnification of liver cells from the TFA- or vehicle-treated LDLr^-/-^ animals, stained red for PMP70, a marker of peroxisome proliferation. Red, AF568 stain for PMP70; blue, DAPI stain for nuclei; green, autofluorescence. **e**) Western blot for PMP70 in liver tissue from the treated LDLr^-/-^ mice.

### TFA induces peroxisome proliferation in vivo and in cell culture

The transcriptomic analysis suggested that TFA may exert its effects, at least in part, by inducing the proliferation of peroxisomes, which are organelles that carry out lipid and fatty acid metabolism [26]. Further support that TFA might act mechanistically via peroxisome proliferation could be found from prior reports that longer chain PFAS, such as perfluorobutyrate and perfluorooctanoate [27–30], and some halogenated acetates [31], can stimulate peroxisome proliferation. However, these previous studies, as well as toxicological studies of TFA [14,32], indicated that TFA only weakly stimulated peroxisome proliferation, if at all. Notably, the observed reductions in triglycerides and cholesterol caused by TFA are generally consistent with what would be expected if TFA was inducing peroxisome proliferation, in analogy to effects caused by fibrate drugs (e.g., fenofibrate and gemfibrozil). This class of drugs, prescribed as lipid-modifying agents to reduce triglyceride and LDL cholesterol levels and increase HDL levels, bring about their phenotypic effects via activation of PPAR-α, resulting in peroxisome proliferation [33,34].

To experimentally establish if TFA was causing peroxisome proliferation, we used a series of in vitro assays involving cultured human and rodent liver cells. We made use of a reporter assay in human liver HepG2 cells transiently transfected with a PPAR response element-driven luciferase gene and a plasmid containing one of the three PPAR isoforms (PPAR-α, PPAR-β, and PPAR-γ) [27,31]. In the case of human (Figure S11) or mouse (Figure 3b) PPAR-α, TFA concentrations of 1 mM or 10 mM significantly activated luciferase expression, as did the positive control fibrate drug ciprofibrate at 0.1 mM. On the other hand, luciferase was not activated by TFA with human or mouse PPAR-β or PPAR-γ isoforms (Figures 4 and S11), implicating PPAR-α as the isoform responsible for TFA-mediated peroxisome proliferation. Luciferase expression was activated by TFA treatment for 6 h, 24 h, or 48 h timepoints (Figure S11). In cultured rat liver FAO cells, gene expression of several peroxisome proliferation-associated genes (e.g., EHHADH) was increased following treatment with TFA, as confirmed by qPCR (Figure 3c). Western blotting of FAO cell lysates also confirmed that TFA treatment led to higher levels of EHHADH protein in the cells (Figure S12). In contrast, TFA completely lost the ability to induce gene expression of EHHADH in FAO cells in which PPAR-α was knocked out (Figure 3c), supporting that TFA’s actions are mediated by PPAR-α. TFA treatment also increased palmitoyl-CoA oxidation activity (a functional assay for peroxisome proliferation) in the rat FAO cells (Figure S13). Neither gene expression changes by qPCR nor changes in palmitoyl-CoA oxidation activity were observed in TFA-treated human liver HepG2 cells, in line with previous findings that rodent liver cells are more sensitive to induction of peroxisome proliferation than human cells [35,36]. Altogether, these experiments support that TFA activates PPAR-α in liver cells, leading to changes in gene expression and enzyme activity associated with peroxisome proliferation.

**Figure 4.**
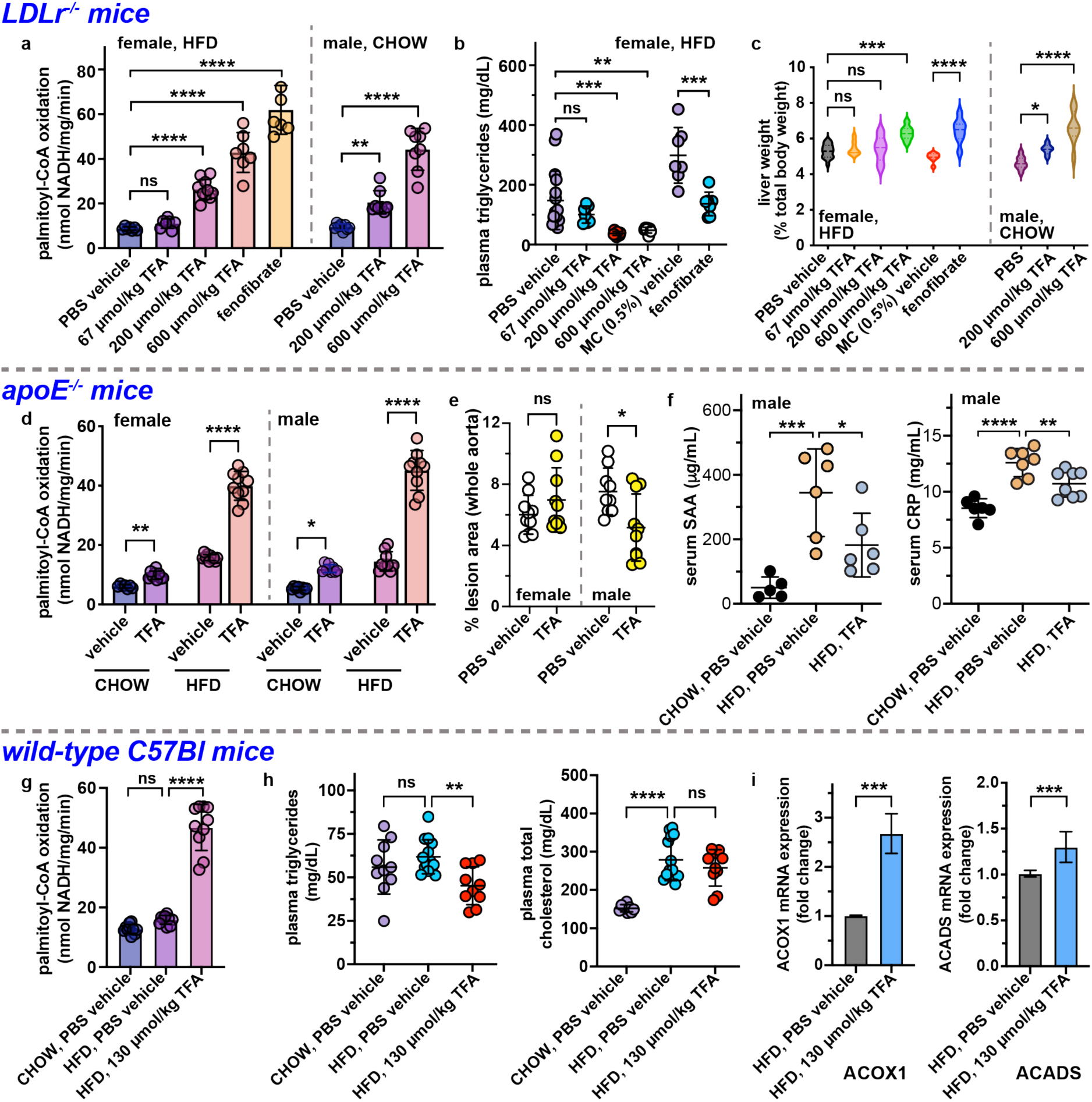
TFA induces various phenotypic effects in LDLr^-/-^ mice, apoE^-/-^ mice, and wild-type C57Bl mice. Daily oral gavage of TFA for two weeks to LDLr^-/-^ mice led to dose-dependent (**a**) increases in liver tissue palmitoyl-CoA oxidase activity, (**b**) decreases in plasma triglyceride levels, and (**c**) increases in liver weight. Fenofibrate in 0.5% methylcellulose (MC) vehicle was used as a positive control; this fibrate drug induces peroxisome proliferation via activation of PPAR-α. (**d**) In both genders of apoE^-/-^ mice, daily administration of TFA (200 µmol/kg) in the drinking water for 10 weeks led to increases in liver tissue palmitoyl-CoA oxidase activity in the context of either a CHOW or HFD diet. In the male apoE^-/-^ mice, the TFA treatment led to (**e**) reductions in atherosclerotic lesions in the aorta and (**f**) reduced levels of inflammation markers SAA and CRP. In wild-type C57Bl male mice, daily oral gavage of 130 µmol/kg TFA for five weeks led to (**g**) increases in liver tissue palmitoyl-CoA oxidase activity, (**h**) reductions in plasma triglyceride and cholesterol levels (although not statistically significant in the case of cholesterol), and (**i**) increased gene expression in the liver for genes associated with peroxisome proliferation, such as ACOX1 and ACADS. In all scatter plots, data are shown as mean ± SD. *p* values were determined by one-way ANOVA comparing the experimental group to its corresponding vehicle group (demarcated by dotted lines); ns, not significantly different; *, *p* < 0.05; **, *p* < 0.01; ***, *p* < 0.001; ****, *p* < 0.0001. The TFA dose was 200 µmol/kg/day unless otherwise noted, and animals were fed a HFD unless a CHOW diet is noted.

We further supported the link between TFA treatment and peroxisome proliferation in vivo. TFA was administered by daily oral gavage to LDLr^-/-^ mice for a 2-wk period, at concentrations ranging from 67 µmol/kg/day to 600 µmol/kg/day. The study included one arm involving female mice fed HFD, as well as another arm involving male mice fed a CHOW diet. As a positive control in the HFD arm, we used the drug fenofibrate, a clinically used fibrate drug that induces peroxisome proliferation by activating PPAR-α. Consistent with the expected effects of peroxisome proliferation, we observed increased levels of peroxisome membrane protein 70 (PMP70), a fatty acid transporter protein that is a marker for peroxisomes, using confocal microscopy (Figure 3d and S14) and Western blotting (Figure 3e) of liver tissue from the treated animals. In both arms of the study, TFA caused dose dependent increases in liver palmitoyl-CoA oxidase activity (Figure 4a), as well as increases in liver weight (Figure 4c), consistent with peroxisome proliferation. TFA treatment also led to reductions in plasma triglyceride levels of the mice fed HFD (Figure 4b), whereas no changes were observed in triglyceride or cholesterol levels of the mice fed a CHOW diet (Figure S15). In general, the effects provoked by TFA were similar to those caused by the positive control peroxisome proliferation inducing agent fenofibrate.

A targeted metabolomics analysis of bile acid levels in the feces and plasma of TFA-treated LDLr-/- mice revealed several notable changes induced by TFA treatment. As a general trend, bile acid levels were lower in the TFA-treated animals, although the changes did not reach the level of statistical significance in most cases. In the plasma, β-muricholic acid (β-MCA), ρο-muricholic acid (ρο-MCA), deoxycholic acid (DCA), and taurochenodeoxycholic acid (TCDCA) were significantly decreased (Figure S16), whereas in the stool levels of ρο-MCA, hyocholic acid (HCA), and lithocholic acid (LCA) were significantly decreased by TFA treatment (Figure S17). The observation of reduced levels of bile acids in TFA-treated mice are consistent with previous reports that peroxisome proliferation causes altered/suppressed bile acid synthesis [37,38].

### TFA treatment induces peroxisome proliferation and exerts anti-atherosclerotic effects in apoE^-/-^ mice

We carried out focused studies in two additional strains of mice, apoE^-/-^ mice and wild-type C57Bl mice, to establish if TFA treatment would cause biological effects in these strains similar to those observed in the LDLr^-/-^ strain. ApoE^-/-^ mice have been widely used for studying the anti-atherogenic effects of drugs, including antihypertensives, LXR agonists, PPAR agonists, statins, etc. [39]. We carried out a 10-week study in which the apoE^-/-^mice were treated with a daily 200 µmol/kg oral dose of TFA in the drinking water (see Figure S1 for dosing schedules). The study included arms (n= 8-10/group) involving male or female mice, fed a standard CHOW diet or a HFD that commenced at the beginning of the TFA treatment. We observed alterations in gene expression indicative of peroxisome proliferation– transcriptomics analysis of liver tissue from both male and female HFD-fed apoE^-/-^ mice, after either three days or two weeks of TFA treatment, revealed expression changes in PPAR-a target genes [26] that were similar to those observed in LDLr^-/-^ mice (Spreadsheet S1), with pathway enrichment analysis indicating upregulation in metabolism of fatty acids and peroxisome proliferation (Spreadsheet S1). We also observed increased palmitoyl-CoA oxidation in liver tissue, in both male and female apoE^-/-^ mice, and in the context of either a standard CHOW diet or HFD (Figure 4d). Unlike the LDLr^-/-^ mice, we did not observe TFA-induced reductions of plasma cholesterol or triglyceride levels in apoE^-/-^ animals (Figure S18). This was not wholly unexpected, because previous studies using various agents (e.g., fibrates, apoA-I mimetic peptides, antibodies) have shown that cholesterol or triglyceride levels are often not reduced in apoE^-/-^ mice, even in cases with beneficial effects on atherosclerotic plaques [24,40–43]. Indeed, analysis of atherosclerotic lesions in the apoE^-/-^ animals indicated that, in male mice, TFA treatment caused a 26% reduction in the of atherosclerotic lesions volumes in the hearts (*p*=0.06, Figure S19, S20) and a 30% reduction in plaque area in en face aortic sections (*p*=0.02, Figure 4e, S21). In the female apoE^-/-^ mice, atherosclerotic lesions were not significantly impacted by TFA treatment (Figures 4e, S19, S22, S23). Consistent with the reduced development of atherosclerosis in the male apoE^-/-^ mice, TFA reduced the circulating levels of SAA and C-reactive protein (CRP), markers of inflammation that correlate with atherosclerosis (Figure 4f).

### TFA induces peroxisome proliferation in wild-type C57Bl mice

Wild type C57Bl mice are used as animal models of certain metabolic diseases, including diet induced obesity and diabetes. To establish whether or not TFA treatment caused observable phenotypic changes in wild type mice modeling metabolic disease, male C57Bl mice were fed a high fat diet for six weeks, at which point TFA treatment by daily oral gavage (130 µmol/kg/day) was commenced; TFA treatment and HFD feeding continued for five weeks, after which animals were sacrificed (see Figure S1 for dosing schedules). Like LDLr^-/-^ and apoE^-/-^ mice, TFA-treated wild-type C57Bl mice exhibited increased palmitoyl-CoA oxidation in liver tissue (Figure 4g). TFA treatment also lowered plasma SAA (Figure S8), triglycerides (Figure 4h), and cholesterol levels in the C57Bl mice (Figure 4h), although the change was not statistically significant in the case of cholesterol. Expression of peroxisome proliferation associated genes (e.g., ACOX1, ACADS) was increased, as determined by qPCR (Figure 4i). We are actively working to more fully characterize the effects of TFA treatment in the context of C57Bl models of diabetes and diet-induced obesity, but this pilot study clearly establishes that TFA impacts fatty acid metabolism in wild type C57Bl mice.

### Trifluoroacetylated metabolites were not observed by metabolomics or biochemical studies

Based on the observed gene expression changes in numerous enzymes involved in CoA biochemistry, and the structural similarity of TFA to acetate, we wondered if TFA might be impacting metabolism by generating trifluoroacetylated metabolites and/or proteins (analogs of acetylated proteins), possibly through the intermediacy of trifluoroacetyl-CoA (analog of acetyl-CoA). As a first step to test this hypothesis, we carried out an untargeted metabolomics analysis to identify metabolites that were significantly changed in concentration following TFA treatment. For the analysis, we compared plasma from female LDLr^-/-^ mice treated by daily oral gavage for two weeks with a 200 µmol/kg TFA daily dose (n=10) or vehicle (n=10). The TFA treatment altered the concentrations of a few hundred plasma metabolites with a fold-change >1.5, p-value < 0.01, and maximum peak intensity >5000 (Spreadsheet S2). We were able to identify about 15 of these metabolites, most of which are phospholipids and fatty acids (Spreadsheet S2). As would be expected, TFA was observed in the plasma of treated mice, but not in the plasma of vehicle control animals. Interestingly, one of the identified metabolites was pantothenic acid, a component of coenzyme A (CoA), which increased about 2-fold in plasma concentration upon TFA treatment. We were unable to identify any metabolites consistent with the presence of a covalently attached trifluoroacetyl moiety, despite searching the mass fragmentation spectra for the presence of trifluoroacetyl fragment ions, and searching for pairs of parent ion masses that differed by the mass of a trifluoroacetyl group (Figure 5). The raw metabolomics mass spectrometry data are accessible at the National Metabolomics Data Repository (Study ID ST003595). Consistent with the absence of trifluoroacetylated metabolites in the plasma, the ^19^F-NMR analysis of plasma samples described above to characterize TFA pharmacokinetics did not reveal any unidentified fluorine signals that might correspond to such metabolites (Figure S4), although the limit of detection in our ^19^F-NMR assay was rather poor (∼10 µM for TFA).

**Figure 5.**
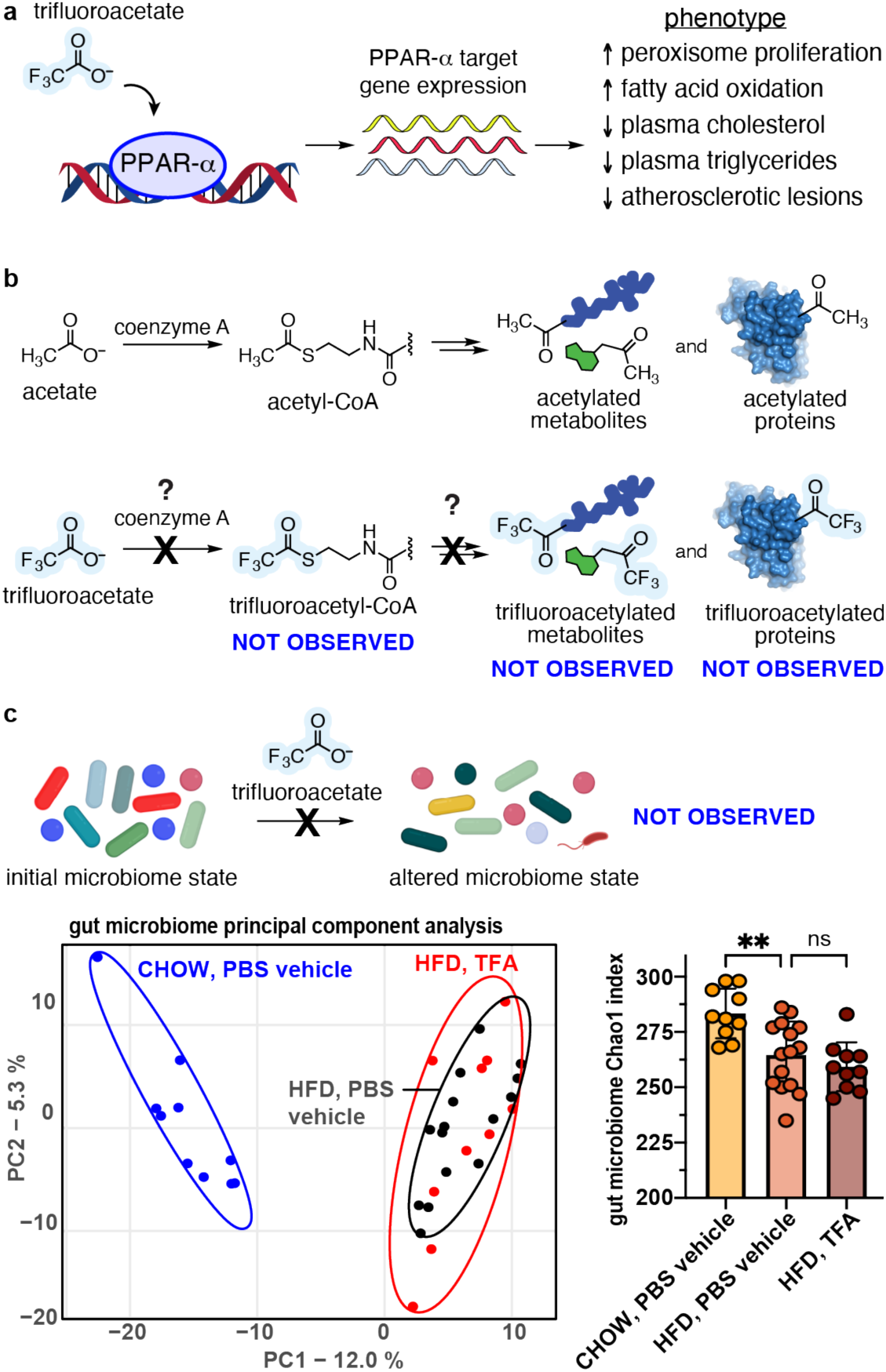
Mechanistically, TFA appears to act by inducing peroxisome proliferation, rather than generating trifluoroacetylated metabolites/proteins or altering the gut microbiome in vivo. **a**) Data including RNA-seq, confocal microscopy, western blotting, metabolomics, and biochemical assays indicated that TFA induces peroxisome proliferation by activating peroxisome proliferator-activated receptor (PPAR)-alpha. **b**) While TFA is structurally similar to acetate, we did not observe trifluoroacetylated metabolites or proteins, nor trifluoroacetyl-CoA, using a combination of metabolomics, proteomics, and biochemical assays. **c**) TFA treatment did not alter the gut microbiome composition, as determined by 16S sequencing of fecal microbiome samples from PBS vehicle- or TFA-treated wild-type C57Bl mice (four-week treatment, 200 µmol/kg/day TFA, n=10-15/group). On the left is shown a Bray-Curtis principal component analysis of beta diversity between samples. On the right, plots of Chao1 index alpha diversity are shown as mean ± SD. *p* values were determined by one-way ANOVA comparing the experimental group to PBS vehicle group; ns, not significantly different; **, *p* < 0.01.

Biochemical assays were used to explore the potential formation of TFA-CoA in vitro. These studies generally followed previously established procedures, in which an organic acid of interest is incubated with rat liver microsomes or liver cell lysate in the presence of ATP and CoA to determine if the acid is converted into the corresponding acyl-CoA derivative [45,46]. Whereas we clearly observed the formation of the acyl-CoA derivatives of acetate and ciprofibrate by LCMS in this assay, we did not observe any evidence of TFA-CoA formation (Figure S26). Likewise, incubation of purified acetyl CoA synthase protein from yeast with acetate and ATP led to the formation of acetyl-CoA, whereas no TFA-CoA was observed when acetate was replaced with TFA. Finally, coincubation of TFA along with acetate or ciprofibrate in the above assays did not affect the formation of either acetyl-CoA or ciprofibrate-CoA (Figure S26), suggesting that the presence of TFA did not inhibit or compete with the other acids as a substrate. Thus, there is no experimental support for the formation of TFA-CoA (Figure 5), although we cannot strictly rule out that TFA-CoA was generated and rapidly hydrolyzed.

### Proteomics analyses did not reveal trifluoroacetylated proteins

Protein acylation, especially with short chain acids such as acetate, can alter cellular phenotypes by modifying the epigenetic and cell signalling landscapes. To investigate if TFA treatment led to the formation of trifluoroacetylated proteins *in vivo*, analogous to protein acetylation, we carried out a quantitative proteomics study using liver tissue from mice (n=3 per group) that had been treated with TFA (200 µmol/kg/day) or vehicle for two weeks. The liver samples were homogenized, labelled using isobaric tandem mass tags (TMT) [47], and analysed by LCMS to identify any potential differences between the treatment conditions. We carried out both an open search of the data using MSFragger [48] to identify unspecified mass differences between predicted and experimentally observed peptide spectra, and a closed search of the data using Proteome Discoverer with trifluoroacetylation of lysine as a potential modification. We observed no evidence of trifluoroacetylated proteins using either of these approaches (Figure 5), though other expected protein modifications were identified (Spreadsheet S3). The raw proteomics mass spectrometry files are accessible at the PRIDE repository (dataset identifier PXD057035).

### TFA treatment did not directly affect the gut microbiome in vitro or in vivo

A number of studies indicated that TFA was not operating through a mechanism involving direct effects on the gut microbiota. Using in vitro gut microbiota cultures of cecal material taken from HFD-fed LDLr^-/-^ mice, we established that overnight treatment with 1 mM TFA did not cause changes to the microbiome composition (Figure S24). In contrast, two positive control gut microbiome remodelling cyclic peptides [44] shifted the in vitro microbiota community into distinct compositions as expected, regardless of whether the peptides were administered as the TFA or the Cl^-^ salt (Figure S24). In a separate in vitro study, the gut microbiota meta-transcriptomics profile of fresh gut microbiota cultures derived from HFD-fed LDLr^-/-^ mice was unchanged following a six-hour in vitro treatment with 1 mM TFA, both for transcription activity at the organism or transcript function level (Figure S24). To test the potential effects of TFA on gut microbiota in vivo, fecal samples from TFA- or vehicle-treated LDLr^-/-^ mice (two-week, 200 µmol/kg/day, n=4/group) or wild-type C57Bl mice (four-week, 200 µmol/kg/day, n=10-15/group) were analysed using 16S sequencing. As expected, the mouse diet (CHOW vs HFD) led to clear differences in gut microbiome composition (Figures 5c and S25). On the other hand, TFA did not cause observable changes in alpha diversity (Chao1 or Shannon index) or beta diversity (Bray-Curtis dissimilarity) in either strain of mouse compared to the vehicle control group (Figures 5c and S25). We conclude that TFA does not directly impact gut microbiota. This conclusion is consistent with the observation that oral and i.p. routes of TFA administration both provoked similar phenotypic outcomes, even though the initial concentrations of TFA in the gut would likely be considerably higher following oral administration.

### Structural analogs of TFA reduce plasma lipid levels in LDLr^-/-^ mice

Intrigued by the observed lipid lowering effects of TFA in vivo, we carried out a small exploration of the structure-activity space by investigating a number of TFA analogs in HFD-fed LDLr^-/-^ mice (Figure 6). The molecules were each administered orally at a 200 µmol/kg daily dose for two weeks, and included the small organic acids acetate, trichloroacetate (TCA), difluoroacetate, 2,2-difluoropropionate, and 2,2-difluorobutyrate. We also examined the 2,2,2-trifluoroacetamide and *N*-ethylacetamide derivatives of TFA, trifluoromethane-sulfonamide, and the carboxylic acid isosteres [49] 5-trifluoromethytetrazole and 3-(trifluoromethyl)-1,2,4-oxadiazol-5-ol. The sulfonamide caused acute toxicity in the mice and was discontinued before completion of the study; the remaining molecules were all well-tolerated. Neither TCA nor acetate affected plasma cholesterol levels in the mice. Likewise, neither of the carboxylic acid isosteres 5-trifluoromethytetrazole or 3-(trifluoromethyl)-1,2,4-oxadiazol-5-ol reduced plasma cholesterol levels. On the other hand, difluoroacetate and 2,2-difluorobutyrate brought about statistically significant reductions in cholesterol levels (Figure 6). Both of the TFA acetamides reduced plasma cholesterol and triglyceride levels; however, follow-up studies revealed that these molecules were relatively quickly converted to TFA in vivo (half-life < 1 h) upon oral administration (Figures S27 and S28). ^19^F-NMR analysis of plasma drawn one hour following oral gavage of 2,2,2-trifluoroacetamide showed a 1.8:1 ratio of TFA to trifluoroacetamide present in plasma, due to conversion of the administered trifluoroacetamide to TFA in vivo (Figure S27). Similarly, at the one-hour timepoint following oral gavage of *N*-ethyl-trifluoroacetamide, we observed a 3.8:1 ratio of TFA to 2,2,2-trifluoroacetamide present in plasma (none of the administered *N*-ethyl-trifluoroacetamide was observed), indicating that *N*-ethyl-trifluoroacetamide is also converted to TFA, possibly through the intermediacy of 2,2,2-trifluoroacetamide (Figure S28). Thus, the reductions in plasma lipid levels observed for the TFA acetamides are likely due to the effects of TFA itself. In contrast, ^19^F-NMR analyses of plasma from mice treated with difluoroacetate or 2,2-difluorobutyrate indicated that each of these analogs was unchanged and observable in plasma at the one-hour timepoint following oral gavage (Figure S29), suggesting that difluoroacetate and 2,2-difluorobutyrate directly cause cholesterol reductions in the mice.

**Figure 6.**
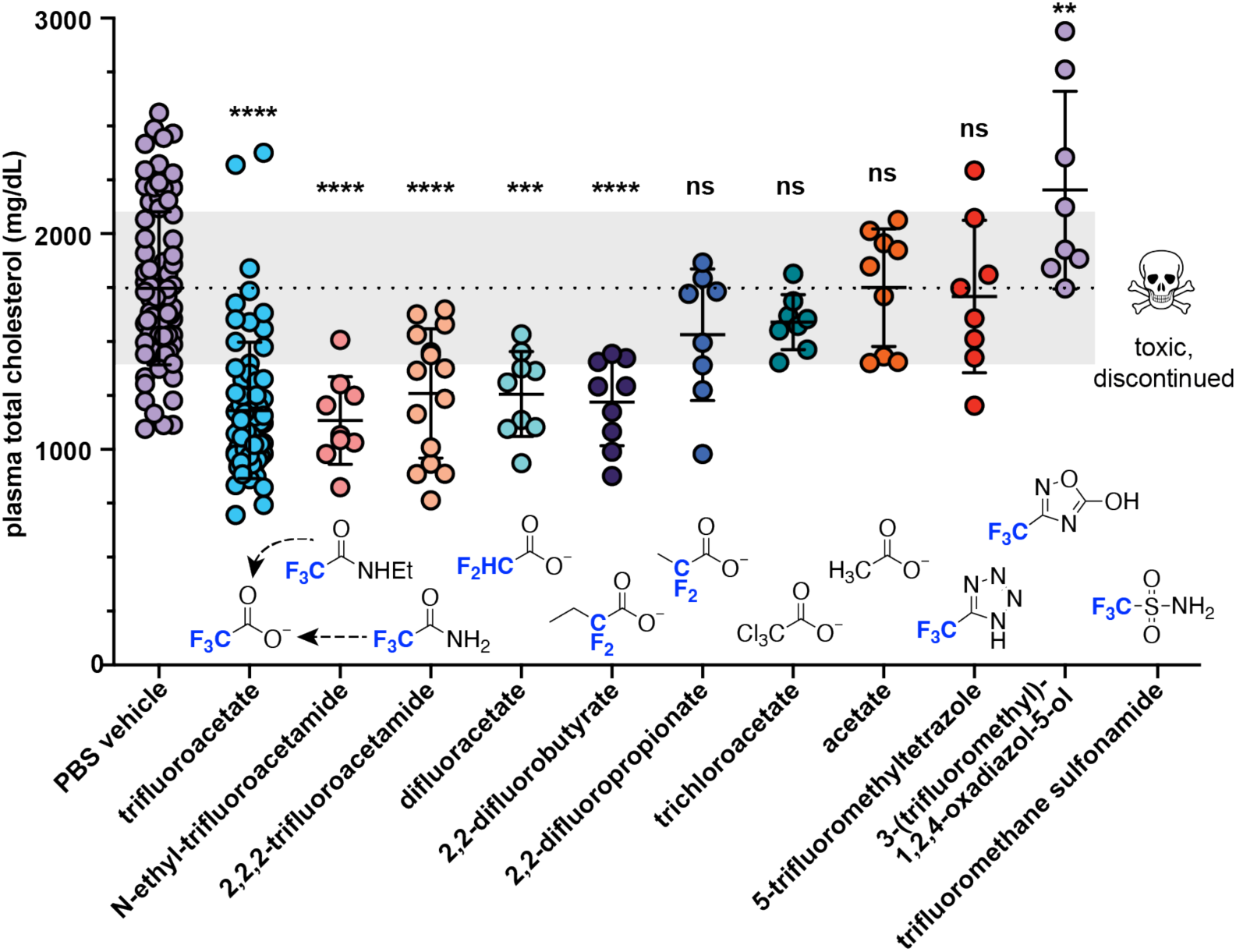
Several structural analogs of TFA reduce plasma cholesterol levels in HFD-fed LDLr^-/-^ mice. Each analog was administered orally for 2 weeks at 200 μmol/kg/day. The reductions in plasma cholesterol levels observed for N-ethyl-trifluoroacetamide and 2,2,2-trifluoroacetamide are likely due to the effects of TFA itself, based on the observation of rapid conversion of these agents into TFA in vivo following oral administration (Figures S22 and S23). N-ethyl-trifluoroacetamide, 2,2,2-trifluoroacetamide, 5-trifluoromethyltetrazole, 3-(trifluoromethyl)-1,2,4-oxadiazol-5-ol, and trifluoromethane sulfonamide were administered by oral gavage, whereas the remainder of the TFA analogs were administered in the drinking water. The data for PBS vehicle and trifluoroacetate include groups treated by oral gavage and groups treated via drinking water. The data are shown as mean ± SD. *p* values were determined by one-way ANOVA comparing the experimental group to PBS vehicle group; ns, not significantly different; **, *p* < 0.01; ***, *p* < 0.001; ****, *p* < 0.0001. The dotted line and gray shaded area correspond to the mean and ± one SD for PBS vehicle, respectively.

## Discussion

Our findings demonstrate that TFA impacts fatty acid and lipid metabolism in numerous biological contexts (LDLr^-/-^, apoE^-/-^, and wild-type C57Bl mice and cultured human HepG2 and rat FAO liver cells), at least in part by promoting peroxisome proliferation. We first observed that TFA treatment reduced the plasma levels of cholesterol and triglycerides in high fat-fed LDLr^-/-^ mice, but subsequently confirmed that TFA also mediated phenotypic changes in apoE^-/-^ and wild-type C57Bl mice, as well as the cultured liver cells. The observed effects of TFA differed somewhat among the three strains of mice. In all three strains, upregulation of PPAR-α target genes and increased palmitoyl-CoA oxidase activity were observed, supporting that TFA treatment had induced peroxisome proliferation. In HFD-fed LDLr^-/-^ mice, significantly reduced plasma cholesterol and triglyceride levels were observed, whereas these plasma lipid levels were unchanged in apoE^-/-^ mice, and triglyceride levels (but not cholesterol) were significantly reduced in wild-type C57Bl mice. On the other hand, TFA treatment reduced the development of atherosclerotic lesions in both LDLr^-/-^ and apoE^-/-^ strains of mice, albeit to a lesser degree in apoE^-/-^ mice and only in male animals. Plasma SAA levels were reduced by TFA treatment in all three strains of mice. The lipid lowering effects of TFA were observed in both genders of LDLr^-/-^ mice. Altogether, these efforts suggest the observed TFA-mediated phenotypic changes are rather general.

Peptides are often employed as TFA salts in early research and development settings, so our findings raise the possibility that TFA may be confounding or contributing to phenotypic changes observed in various in vitro and in vivo studies. Since TFA is present in peptide samples as a counterion of basic residues (Lys, Arg, His, or free N-terminus), the amount of TFA with a given peptide will depend on the number of basic sites in the peptide sequence. Most commonly, helical apolipoprotein mimetic peptides range from 18-37 residues in length, contain multiple basic residues, and are administered in the range of 20–60 mg/kg, but sometimes up to 100 mg/kg [2,50–52]. A 40 mg/kg peptide dose would be expected to contain ∼70 µmol/kg TFA for **4F** or **mR18L** (four basic sites each), 85 µmol/kg TFA for **5A** (nine basic sites), and 105 µmol/kg TFA for **ELK-2A** (11 basic sites) [53–55]. Under the conditions of our studies, TFA concentrations of 200 µmol/kg reproducibly reduced plasma cholesterol levels compared to vehicle in HFD-fed LDLr^-/-^ mice, while concentrations of 67 µmol/kg caused reductions in lipid levels that did not reach the degree of statistical significance. Further studies are warranted to determine if the presence of TFA may be unexpectedly contributing to the outcomes of biological studies involving apolipoprotein mimetic peptides, or peptides in other contexts. It is notable that the phenotypic effects brought about by TFA (e.g., reduced plasma lipid levels, decreased development of atherosclerosis, lowered biomarkers of inflammation like SAA and CRP) are similar to those that are typically sought for apolipoprotein mimetic peptides, potentially increasing the possibility that TFA would play a confounding role in this area of research.

Given the prevalence of TFA as a counterion for synthetic peptides, it is surprising that the lipid lowering effects of TFA have not been reported before now. To our knowledge, the only previous indication that TFA can reduce lipid levels in vivo is to be found in a 2014 patent [56], which claimed that orally administered calcium trifluoroacetate lowered plasma cholesterol levels in a male Gottingen minipig (n=1, daily administration for four weeks), as did intravenously administered calcium trifluoroacetate in male or female beagle dogs (n=2 or 4 animals/sex/group, daily administration for 2-4 weeks). The TFA doses reported in the patent ranged from 380 µmol/kg/day up to 7,500 µmol/kg/day, which are higher than those used in the studies described here (typically, 200 µmol/kg/day). The patent does not describe the diets of the treated animals, nor does it give any discussion of the relevant mechanism(s) of action for calcium trifluoroacetate. The data reported in the patent do not appear to have been published in a scientific journal, nor has the patent been previously cited.

Earlier studies of peroxisome proliferation mediated by perfluorinated acids (PFAS) or halogenated acetates, generally carried out in cell culture, have reported that TFA only weakly stimulates peroxisome proliferation, if at all [14,28–32]. In contrast, multiple lines of evidence from our studies (including gene expression changes, confocal microscopy, western blotting, metabolomics, and biochemical assays) support that TFA induces robust peroxisome proliferation by activating the PPAR-α pathway. This discrepancy could be due to a number of factors, such as differences in TFA dose, duration of treatment, or different biological systems under study. Also, the animals under study here typically consumed a high fat diet, which may make the effects of TFA more pronounced because of the higher plasma lipid levels present. Follow-up experiments are needed to establish whether TFA directly binds to PPAR-α as an exogenous ligand, or activates PPAR-α through a ligand-independent pathway.

While our findings implicate a mechanism involving peroxisome proliferation for TFA-induced lipid perturbations, it should be noted that other biological pathways may also play a role. For example, GPR81 and GPR109a are G-protein coupled receptors for lactate and butyrate, respectively, but they also bind other small organic salts, including TFA [57–59]. TFA binds to GPR81 with similar affinity as lactate (∼5 mM EC_50_), albeit with only ∼60% of the maximum response stimulated by lactate [58]; TFA can also bind to GPR109a at high (>10 mM) concentrations [58]. Activation of these receptors inhibits the breakdown of fatty acids in adipose tissue, resulting in reduced transport of fatty acids to the liver [60]. The beneficial effects of the drug niacin, including increased HDL levels, lowered LDL, and lowered triglycerides are due to activation of GPR109a [59]. Additionally, GPR81 is involved in regulating immune tolerance and suppressing inflammation [61]. Thus, the known phenotypic effects associated with activation of GPR81 and/or GPR109a would be consistent with some of those observed for TFA-treated mice, although the observed plasma concentrations of TFA following our typical 200 µmol/kg/day dose (∼450 µM maximum concentration) were considerably lower than the reported TFA EC_50_ values for either receptor [58].

We also considered the possibility that the effects of TFA were due to the formation of a transient trifluoroacetyl-CoA intermediate that perturbed metabolic pathways in vivo. This hypothesis was based on the structural similarity of TFA to acetate, the previous observations that some exogenous organic acids (including fibrate drugs) are converted to the corresponding acyl-CoA derivatives in rat microsomes [45,46], and the changes we observed in gene expression for numerous enzymes involved in CoA processing (Spreadsheet S1). In the case of TFA, our experiments do not support the formation of TFA-CoA species or other covalent trifluoroacetylated metabolites or proteins. We were not able to detect evidence of small molecule metabolites containing trifluoroacetyl groups using untargeted metabolomics, nor did we detect any signs of trifluoroacetylated proteins using a proteomics approach, nor did we detect the formation of TFA-CoA with liver microsomes or purified acetyl-CoA synthetase. Thus, while we cannot rule out the possibility that TFA-CoA species are generated in vivo and hydrolyzed/metabolized too rapidly to detect, it appears more likely that the altered gene expression observed for numerous enzymes involved in CoA biochemistry is a reflection of their involvement in processing acyl-CoA intermediates in fatty acid metabolism during peroxisome proliferation (Figure 5) [29,62]. Trifluoroacetylated proteins are observed in patients following treatment with certain inhaled anaesthetics such as halothane, although these proteins are believed to arise not from TFA but rather trifluoroacetyl chloride produced via oxidative metabolism of the anaesthetic [19,63].

In addition to TFA, we found that difluoracetate and 2,2-difluorobutyrate also significantly reduced plasma cholesterol levels in LDLr^-/-^ mice, while several other TFA analogs did not. It is somewhat surprising that trichloroacetate (TCA) did not affect lipid levels in the mice, because TCA is reported to more effectively induce peroxisome proliferation than TFA in cell culture [31]. The lack of in vivo efficacy for TCA could be due to poor adsorption or distribution compared to TFA, or its metabolism in vivo. We observed that 2,2,2-trifluoroacetamide and N-ethyltrifluoroacetamide underwent relatively rapid conversion to TFA (half-life < 1 h) following oral administration to the mice. N-ethyltrifluoroacetamide gave rise to both TFA and trifluoroacetamide. N-deethylation of N-ethyl acetamides to the corresponding acetamide (CONHEt → CONH_2_) has been observed previously in the microsomal metabolism of N,N-diethyl-toluamide (DEET) [64]. In other molecular contexts, trifluoracetamide groups are more stable against metabolism, as exemplified by Paxlovid (nirmatrelvir) in which only ∼4% of the dose administered to humans is eventually converted to the des-trifluoroacetylated metabolite [65].

Although TFA or its analogues might have therapeutic applications, the findings reported here may have other implications for human health. TFA is found at high levels in humans relative to other PFAS [18]. Based on the limited studies available, TFA contamination seems to be ubiquitous in the environment [66,67], and environmental TFA levels are predicted to increase in coming decades due to newer PFAS alternatives [13,16]. Another potential source of human exposure to TFA is treatment with the inhalation anesthetics such as halothane and isoflurane [19], which are among the most commonly used agents for induction and maintenance of general anesthesia for surgery. These anesthetics are metabolized in the liver to TFA as a major metabolite [19–21]. In one study, peak plasma concentrations of TFA in patients following treatment with halothane were reported to be 580 µM at 24 h following the start of anesthesia, and remained higher than 200 µM for at least 96 h [20]. These plasma TFA levels are higher than those found here to cause phenotypic changes in mice (∼450 µM maximum TFA plasma concentration following a 200 µmol/kg/day dose), although treatment of the mice involved daily dosing with TFA for a two-week duration. Whereas newer anesthetics such as isoflurane (∼2% metabolized to TFA) and desflurane (∼0.2% metabolized to TFA) have become the standard of care in developed nations, halothane is still widely used in the developing world (∼50% metabolized to TFA) [19,63]. We observed that TFA treatment activated a human PPAR-α reporter assay in human liver HepG2 cells, although neither gene expression changes nor changes in palmitoyl-CoA oxidation activity were observed in the cultured human liver cells, which is consistent with previous findings that rodent liver cells are more sensitive to induction of peroxisome proliferation than human cells [35,36]. In light of these findings, further studies are warranted to better characterize the response of human tissues to TFA exposures.

Supplementary Information is available free of charge at xxxx:

Supplemental figures and methods (PDF)

Spreadsheet S1 providing analysis of RNAseq data (XLSX)

Spreadsheet S2 providing analysis of untargeted metabolomics data (XLSX) Spreadsheet S3 providing analysis of proteomics data (XLSX)

Mass spectrometry proteomics data have been deposited to the ProteomeXchange Consortium via the PRIDE partner repository with the dataset identifier PXD057035. The untargeted metabolomics data for this study is available for download from the National Metabolomics Data Repository with the Study ID ST003595 at http://dx.doi.org/10.21228/M88R7M. The RNA-seq data for this study is available for download from Bioproject at accession # PRJNA917696.

## Acknowledgements

We gratefully acknowledge funding from the National Institutes of Health (NHLBI grant R01HL118114 to M.R.G. and NIGMS grant R01GM102491 to A.S.), the Skaggs Institute of Chemical Biology (M.R.G.), and the Ferring Foundation (A.S.). We thank the Scripps Research Genomics, Nuclear Magnetic Resonance, and Metabolomics and Mass Spectrometry cores for their expert advice and technical assistance.

## References

1. Wolkowicz, P., White, C. R., Anantharamaiah, G. M. Apolipoprotein mimetic peptides: An emerging therapy against diabetic inflammation and dyslipidemia. Biomolecules 2021, 11, 627.

2. Leman, L. J., Maryanoff, B. E., Ghadiri, M. R. Molecules that mimic apolipoprotein A-I: Potential agents for treating atherosclerosis. J. Med. Chem. 2014, 57, 2169–2196.

3. Wolska, A., Reimund, M., Sviridov, D. O., Amar, M. J., Remaley, A. T. Apolipoprotein mimetic peptides: Potential new therapies for cardiovascular diseases. Cells 2021, 10, 597.

4. Ditiatkovski, M., Palsson, J., Chin-Dusting, J., Remaley, A. T., Sviridov, D. Apolipoprotein A-I mimetic peptides. Discordance between in vitro and in vivo properties—brief report. Arterioscler. Thromb. Vasc. Biol. 2017, 37, 1301–1306.

5. Sikora, K., Jaśkiewicz, M., Neubauer, D., Migoń, D., Kamysz, W. The role of counter-ions in peptides—an overview. Pharmaceuticals 2020, 13, 442.

6. Liu, C., Wu, J., Zhu, J., Kuei, C., Yu, J., Shelton, J., Sutton, S. W., Li, X., Yun, S. J., Mirzadegan, T., Mazur, C., Kamme, F., et al. Lactate inhibits lipolysis in fat cells through activation of an orphan G-protein-coupled receptor, GPR81. J. Biol. Chem. 2009, 284, 2811–2822.

7. Schmidt, J., Smith, N. J., Christiansen, E., Tikhonova, I. G., Grundmann, M., Hudson, B. D., Ward, R. J., Drewke, C., Milligan, G., Kostenis, E., Ulven, T. Selective orthosteric free fatty acid receptor 2 (FFA2) agonists: Identification of the structural and chemical requirements for selective activation of FFA2 versus FFA3. J. Biol. Chem. 2011, 286, 10628–10640.

8. Han, J., Kim, N., Kim, E. Trifluoroacetic acid activates ATP-sensitive K+ channels in rabbit ventricular myocytes. Biochem. Biophys. Res. Comm. 2001, 285, 1136–1142.

9. Tipps, M. E., Iyer, S. V., John Mihic, S. Trifluoroacetate is an allosteric modulator with selective actions at the glycine receptor. Neuropharmacol. 2012, 63, 368–373.

10. Ardino, C., Sannio, F., Pasero, C., Botta, L., Dreassi, E., Docquier, J.-D., D’Agostino, I. The impact of counterions in biological activity: Case study of antibacterial alkylguanidino ureas. Mol. Div. 2023, 27, 1489–1499.

11. Cornish, J., Callon, K. E., Lin, C. Q., Xiao, C. L., Mulvey, T. B., Cooper, G. J., Reid, I. R. Trifluoroacetate, a contaminant in purified proteins, inhibits proliferation of osteoblasts and chondrocytes. Am. J. Physiol. 1999, 277, E779–783.

12. Bussolati, B., Ribatti, D., Munaron, L., Bartorelli, A., Bussolati, G. Anti-angiogenic properties of calcium trifluoroacetate. Microvasc. Res. 2009, 78, 272–277.

13. Arp, H. P. H., Gredelj, A., Glüge, J., Scheringer, M., Cousins, I. T. The global threat from the irreversible accumulation of trifluoroacetic acid (TFA). Environ. Sci. Technol. 2024, 58, 19925–19935.

14. Dekant, W., Dekant, R. Mammalian toxicity of trifluoroacetate and assessment of human health risks due to environmental exposures. Arch. Toxicol. 2023, 97, 1069–1077.

15. Scheurer, M., Nödler, K., Freeling, F., Janda, J., Happel, O., Riegel, M., Müller, U., Storck, F. R., Fleig, M., Lange, F. T., Brunsch, A., Brauch, H.-J. Small, mobile, persistent: Trifluoroacetate in the water cycle – overlooked sources, pathways, and consequences for drinking water supply. Water Res. 2017, 126, 460–471.

16. Pickard, H. M., Criscitiello, A. S., Persaud, D., Spencer, C., Muir, D. C. G., Lehnherr, I., Sharp, M. J., De Silva, A. O., Young, C. J. Ice core record of persistent short-chain fluorinated alkyl acids: Evidence of the impact from global environmental regulations. Geophys. Res. Lett. 2020, 47, e2020GL087535.

17. Freeling, F., Björnsdotter, M. K. Assessing the environmental occurrence of the anthropogenic contaminant trifluoroacetic acid (TFA). Curr. Op. Green Sus. Chem. 2023, 41, 100807.

18. Duan, Y., Sun, H., Yao, Y., Meng, Y., Li, Y. Distribution of novel and legacy per-/polyfluoroalkyl substances in serum and its associations with two glycemic biomarkers among chinese adult men and women with normal blood glucose levels. Environ. Int. 2020, 134, 105295.

19. Kharasch, E. Adverse drug reactions with halogenated anesthetics. Clin. Pharmacol. Ther. 2008, 84, 158–162.

20. Kharasch, E. D., Hankins, D., Mautz, D., Thummel, K. E. Identification of the enzyme responsible for oxidative halothane metabolism: Implications for prevention of halothane hepatitis. Lancet 1996, 347, 1367–1371.

21. Hitt, Ben A., Mazze, Richard I., Cousins, Michael J., Edmunds, Henry N., Barr, Gary A., Trudell, James R. Metabolism of isoflurane in Fischer 344 rats and man. Anesthesiology 1974, 40, 62–67.

22. Zhao, Y., Black, A. S., Bonnet, D. J., Maryanoff, B. E., Curtiss, L. K., Leman, L. J., Ghadiri, M. R. In vivo efficacy of HDL-like nanolipid particles containing multivalent peptide mimetics of apolipoprotein A-I. J. Lipid Res. 2014, 55, 2053–2063.

23. Zhao, Y., Imura, T., Leman, L. J., Curtiss, L. K., Maryanoff, B. E., Ghadiri, M. R. Mimicry of high-density lipoprotein: Functional peptide-lipid nanoparticles based on multivalent peptide constructs. J. Am. Chem. Soc. 2013, 135, 13414–13424.

24. Navab, M., Anantharamaiah, G. M., Reddy, S. T., Hama, S., Hough, G., Frank, J. S., Grijalva, V. R., Ganesh, V. K., Mishra, V. K., Palgunachari, M. N., Fogelman, A. M. Oral small peptides render HDL antiinflammatory in mice and monkeys and reduce atherosclerosis in ApoE null mice. Circ. Res. 2005, 97, 524–532.

25. Andrushchenko, V. V., Vogel, H. J., Prenner, E. J. Optimization of the hydrochloric acid concentration used for trifluoroacetate removal from synthetic peptides. J. Pept. Sci. 2007, 13, 37–43.

26. Rakhshandehroo, M., Knoch, B., Müller, M., Kersten, S. Peroxisome proliferator-activated receptor alpha target genes. PPAR Research 2010, 2010, 612089.

27. Takacs, M. L., Abbott, B. D. Activation of mouse and human peroxisome proliferator– activated receptors (α, β/δ, γ) by perfluorooctanoic acid and perfluorooctane sulfonate. Toxicol. Sci. 2006, 95, 108–117.

28. Permadi, H., Lundgren, B., Andersson, K., Sundberg, C., Depierre, J. W. Effects of perfluoro fatty acids on peroxisome proliferation and mitochondrial size in mouse liver: Dose and time factors and effect of chain length. Xenobiotica 1993, 23, 761–770.

29. Just, W. W., Gorgas, K., Hartl, F. U., Heinemann, P., Salzer, M., Schimassek, H. Biochemical effects and zonal heterogeneity of peroxisome proliferation induced by perfluorocarboxylic acids in rat liver. Hepatology 1989, 9, 570–581.

30. Bjork, J. A., Wallace, K. B. Structure-activity relationships and human relevance for perfluoroalkyl acid–induced transcriptional activation of peroxisome proliferation in liver cell cultures. Toxicol. Sci. 2009, 111, 89–99.

31. Walgren, J. L., Jollow, D. J., McMillan, J. M. Induction of peroxisome proliferation in cultured hepatocytes by a series of halogenated acetates. Toxicology 2004, 197, 189–197.

32. Summary of toxicological and metabolism studies for flurtamone. https://www.bayer.com/sites/default/files/m-482307-01-5.pdf. Bayer CropScience, 2014.

33. Kim, N. H., Kim, S. G. Fibrates revisited: Potential role in cardiovascular risk reduction. Diabetes Metab. J. 2020, 44, 213–221.

34. Hong, F., Xu, P., Zhai, Y. The opportunities and challenges of peroxisome proliferator-activated receptors ligands in clinical drug discovery and development. Int. J. Mol. Sci. 2018, 19, 2189.

35. Gonzalez, F. J., Shah, Y. M. Pparα: Mechanism of species differences and hepatocarcinogenesis of peroxisome proliferators. Toxicology 2008, 246, 2–8.

36. Lawrence, J. W., Li, Y., Chen, S., DeLuca, J. G., Berger, J. P., Umbenhauer, D. R., Moller, D. E., Zhou, G. Differential gene regulation in human versus rodent hepatocytes by peroxisome proliferator-activated receptor (PPAR) α: Pparα fails to induce peroxisome proliferation-associated genes in human cells independently of the level of receptor expression. J. Biol. Chem. 2001, 276, 31521–31527.

37. Post, S. M., Duez, H., Gervois, P. P., Staels, B., Kuipers, F., Princen, H. M. G. Fibrates suppress bile acid synthesis via peroxisome proliferator–activated receptor-α–mediated downregulation of cholesterol 7α-hydroxylase and sterol 27-hydroxylase expression. Arterioscler. Thromb. Vasc. Biol. 2001, 21, 1840–1845.

38. Hunt, M. C., Yang, Y.-Z., Eggertsen, G., Carneheim, C. M., Gåfvels, M., Einarsson, C., Alexson, S. E. H. The peroxisome proliferator-activated receptor α (PPARα) regulates bile acid biosynthesis. J. Biol. Chem. 2000, 275, 28947–28953.

39. Jawien, J. The role of an experimental model of atherosclerosis: apoE-knockout mice in developing new drugs against atherogenesis. Curr. Pharm. Biotechnol. 2012, 13, 2435–2439.

40. Morgantini, C., Imaizumi, S., Grijalva, V., Navab, M., Fogelman, A. M., Reddy, S. T. Apolipoprotein A-I mimetic peptides prevent atherosclerosis development and reduce plaque inflammation in a murine model of diabetes. Diabetes 2010, 59, 3223–3228.

41. Dunér, P., Mattisson, I. Y., Fogelstrand, P., Glise, L., Ruiz, S., Farina, C., Borén, J., Nilsson, J., Bengtsson, E. Antibodies against apoB100 peptide 210 inhibit atherosclerosis in apoE-/- mice. Sci. Rep. 2021, 11, 9022.

42. Wouters, K., Shiri-Sverdlov, R., Gorp, P. J. v., Bilsen, M. v., Hofker, M. H. Understanding hyperlipidemia and atherosclerosis: Lessons from genetically modified apoE and LDLr mice. Clin. Chem. Lab. Med. 2005, 43, 470–479.

43. DeClercq, V., Yeganeh, B., Moshtaghi-Kashanian, G.-R., Khademi, H., Bahadori, B., Moghadasian, M. H. Paradoxical effects of fenofibrate and nicotinic acid in apo E-deficient mice. J. Cardiovasc. Pharmacol. 2005, 46, 18–24.

44. Chen, P. B., Black, A. S., Sobel, A. L., Zhao, Y., Mukherjee, P., Molparia, B., Moore, N. E., Aleman Muench, G. R., Wu, J., Chen, W., Pinto, A. F. M., Maryanoff, B. E., et al. Directed remodeling of the mouse gut microbiome inhibits the development of atherosclerosis. Nat. Biotechnol. 2020, 38, 1288–1297.

45. Bronfman, M., Amigo, L., Morales, M. N. Activation of hypolipidaemic drugs to acyl-coenzyme A thioesters. Biochem. J. 1986, 239, 781–784.

46. Aarsland, A., Berge, R. K. Peroxisome proliferating sulphur- and oxy-substituted fatty acid analogues are activated to acyl coenzyme A thioesters. Biochem. Pharmacol. 1991, 41, 53–61.

47. McAlister, G. C., Huttlin, E. L., Haas, W., Ting, L., Jedrychowski, M. P., Rogers, J. C., Kuhn, K., Pike, I., Grothe, R. A., Blethrow, J. D., Gygi, S. P. Increasing the multiplexing capacity of TMTs using reporter ion isotopologues with isobaric masses. Anal. Chem. 2012, 84, 7469–7478.

48. Yu, F., Teo, G. C., Kong, A. T., Haynes, S. E., Avtonomov, D. M., Geiszler, D. J., Nesvizhskii, A. I. Identification of modified peptides using localization-aware open search. Nat. Comm. 2020, 11, 4065.

49. Lassalas, P., Gay, B., Lasfargeas, C., James, M. J., Tran, V., Vijayendran, K. G., Brunden, K. R., Kozlowski, M. C., Thomas, C. J., Smith, A. B., III, Huryn, D. M., Ballatore, C. Structure property relationships of carboxylic acid isosteres. J. Med. Chem. 2016, 59, 3183–3203.

50. Reddy, S. T., Navab, M., Anantharamaiah, G. M., Fogelman, A. M. Apolipoprotein A-I mimetics. Curr. Opin. Lipidol. 2014, 25, 304–308.

51. Chattopadhyay, A., Navab, M., Hough, G., Gao, F., Meriwether, D., Grijalva, V., Springstead, J. R., Palgnachari, M. N., Namiri-Kalantari, R., Su, F., Van Lenten, B. J., Wagner, A. C., et al. A novel approach to oral apoA-I mimetic therapy. J. Lipid Res. 2013, 54, 995–1010.

52. Su, F., Kozak, K. R., Imaizumi, S., Gao, F., Amneus, M. W., Grijalva, V., Ng, C., Wagner, A., Hough, G., Farias-Eisner, G., Anantharamaiah, G. M., Van Lenten, B. J., et al. Apolipoprotein A-I (apoA-I) and apoA-I mimetic peptides inhibit tumor development in a mouse model of ovarian cancer. Proc. Natl. Acad. Sci. U.S.A. 2010, 107, 19997–20002.

53. Datta, G., Chaddha, M., Hama, S., Navab, M., Fogelman, A., Garber, D. W., Mishra, V. K., Epand, R. M., Epand, R. F., Lund-Katz, S., Phillips, M. C., Segrest, J. P., et al. Effects of increasing hydrophobicity on the physical-chemical and biological properties of a class A amphipathic helical peptide. J. Lipid Res. 2001, 42, 1096–1004.

54. Handattu, S. P., Datta, G., Epand, R. M., Epand, R. F., Palgunachari, M. N., Mishra, V. K., Monroe, C. E., Keenum, T. D., Chaddha, M., Anantharamaiah, G. M., Garber, D. W. Oral administration of l-mr18l, a single domain cationic amphipathic helical peptide, inhibits lesion formation in ApoE null mice. J. Lipid Res. 2010, 51, 3491–3499.

55. D’Souza, W., Stonik, J. A., Murphy, A., Demosky, S. J., Sethi, A. A., Moore, X. L., Chin-Dusting, J., Remaley, A. T., Sviridov, D. Structure/function relationships of apolipoprotein A-I mimetic peptides: Implications for antiatherogenic activities of high-density lipoprotein. Circ. Res. 2010, 107, 217–227.

56. Bulgheroni, A., Legora, M., Ceriani, D., Mailland, F. (Inventors), Polichem S.A. (Assignee). Use of trifluoroacetic acid and salts thereof to treat hypercholesterolemia. Patent EP2796135, 2014.

57. Chai, J. T., Digby, J. E., Choudhury, R. P. GPR109A and vascular inflammation. Curr. Atheroscler. Rep. 2013, 15, 325.

58. Liu, C., Wu, J., Zhu, J., Kuei, C., Yu, J., Shelton, J., Sutton, S. W., Li, X., Yun, S. J., Mirzadegan, T., Mazur, C., Kamme, F., et al. Lactate inhibits lipolysis in fat cells through activation of an orphan G-protein-coupled receptor, GPR81. J. Biol. Chem. 2009, 284, 2811–2822.

59. Wanders, D., Judd, R. L. Future of GPR109A agonists in the treatment of dyslipidaemia. Diabet. Obesity Metab. 2011, 13, 685–691.

60. Dvorak, C. A., Liu, C., Shelton, J., Kuei, C., Sutton, S. W., Lovenberg, T. W., Carruthers, N. I. Identification of hydroxybenzoic acids as selective lactate receptor (GPR81) agonists with antilipolytic effects. ACS Med. Chem. Lett. 2012, 3, 637–639.

61. Ranganathan, P., Shanmugam, A., Swafford, D., Suryawanshi, A., Bhattacharjee, P., Hussein, M. S., Koni, P. A., Prasad, P. D., Kurago, Z. B., Thangaraju, M., Ganapathy, V., Manicassamy, S. GPR81, a cell-surface receptor for lactate, regulates intestinal homeostasis and protects mice from experimental colitis. J. Immunol. 2018, 200, 1781.

62. Ball, M. R., Gumaa, K. A., McLean, P. Effect of clofibrate on the CoA thioester profile in rat liver. Biochem. Biophys. Res. Commun. 1979, 87, 489–496.

63. Njoku, D., Laster, M. J., Gong, D. H., Eger, E. I., Reed, G. F., Martin, J. L. Biotransformation of halothane, enflurane, isoflurane, and desflurane to trifluoroacetylated liver proteins: Association between protein acylation and hepatic injury. Anesth. Analges. 1997, 84, 173–178.

64. Constantino, L., Iley, J. I. M. Microsomal metabolism of N,N-diethyl-m-toluamide (DEET, DET): The extended network of metabolites. Xenobiotica 1999, 29, 409–416.

65. Singh, R. S. P., Walker, G. S., Kadar, E. P., Cox, L. M., Eng, H., Sharma, R., Bergman, A. J., Van Eyck, L., Hackman, F., Toussi, S. S., Kalgutkar, Amit S., Obach, R. S. Metabolism and excretion of nirmatrelvir in humans using quantitative fluorine nuclear magnetic resonance spectroscopy: A novel approach for accelerating drug development. Clin. Pharmacol. Ther. 2022, 112, 1201–1206.

66. Freeling, F., Scheurer, M., Koschorreck, J., Hoffmann, G., Ternes, T. A., Nödler, K. Levels and temporal trends of trifluoroacetate (TFA) in archived plants: Evidence for increasing emissions of gaseous TFA precursors over the last decades. *Environ. Sci*, Tech. Lett. 2022, 9, 400–405.

67. Björnsdotter, M. K., Yeung, L. W. Y., Kärrman, A., Jogsten, I. E. Ultra-short-chain perfluoroalkyl acids including trifluoromethane sulfonic acid in water connected to known and suspected point sources in sweden. *Environ*. Sci. Tech. 2019, 53, 11093–11101.

68. Mullick, A. E., Tobias, P. S., Curtiss, L. K. Modulation of atherosclerosis in mice by toll-like receptor 2. J. Clin. Invest. 2005, 115, 3149–3156.

69. Patel, H., Ewels, P., Manning, J., Garcia, M. U., Peltzer, A., Hammarén, R., Botvinnik, O., Talbot, A., Sturm, G., bot, n.-c., Zepper, M., Moreno, D., et al. Nf-core/rnaseq: Nf-core/rnaseq v3.15.1 - Augmented Aluminium Axolotl 2024.

70. Ulgen, E., Ozisik, O., Sezerman, O. U. Pathfindr: An R package for comprehensive identification of enriched pathways in omics data through active subnetworks. *Front*. Genetics 2019, 10,

71. Concordet, J.-P., Haeussler, M. Crispor: Intuitive guide selection for CRISPR/Cas9 genome editing experiments and screens. Nuc. Acids Res. 2018, 46, W242–W245.

72. Ran, F. A., Hsu, P. D., Wright, J., Agarwala, V., Scott, D. A., Zhang, F. Genome engineering using the CRISPR-Cas9 system. Nat. Prot. 2013, 8, 2281–2308.

73. Wegner, K., Just, S., Gau, L., Mueller, H., Gerard, P., Lepage, P., Clavel, T., Rohn, S. Rapid analysis of bile acids in different biological matrices using LC-ESI-MS/MS for the investigation of bile acid transformation by mammalian gut bacteria. Anal. Bioanal. Chem. 2017, 409, 1231–1245.

74. Conway, L. P., Jadhav, A. M., Homan, R. A., Li, W., Rubiano, J. S., Hawkins, R., Lawrence, R. M., Parker, C. G. Evaluation of fully-functionalized diazirine tags for chemical proteomic applications. Chem. Sci. 2021, 12, 7839–7847.

75. Perez-Riverol, Y., Bai, J., Bandla, C., García-Seisdedos, D., Hewapathirana, S., Kamatchinathan, S., Kundu, Deepti J., Prakash, A., Frericks-Zipper, A., Eisenacher, M., Walzer, M., Wang, S., et al. The PRIDE database resources in 2022: A hub for mass spectrometry-based proteomics evidences. Nuc. Acids Res. 2021, 50, D543–D552.

76. Kong, A. T., Leprevost, F. V., Avtonomov, D. M., Mellacheruvu, D., Nesvizhskii, A. I. MSFragger: Ultrafast and comprehensive peptide identification in mass spectrometry–based proteomics. Nat. Meth. 2017, 14, 513–520.

77. Chang, H.-Y., Kong, A. T., da Veiga Leprevost, F., Avtonomov, D. M., Haynes, S. E., Nesvizhskii, A. I. Crystal-C: A computational tool for refinement of open search results. J. Proteome Res. 2020, 19, 2511–2515.

78. Teo, G. C., Polasky, D. A., Yu, F., Nesvizhskii, A. I. Fast deisotoping algorithm and its implementation in the MSFragger search engine. J. Proteome Res. 2021, 20, 498–505.

79. Geiszler, D. J., Kong, A. T., Avtonomov, D. M., Yu, F., Leprevost, F. d. V., Nesvizhskii, A. I. PTM-Shepherd: Analysis and summarization of post-translational and chemical modifications from open search results. Mol. Cell. Proteomics 2021, 20, 100018.

