## Supplemental Figures and Methods for "Trifluoroacetate reduces plasma lipid levels and the development of atherosclerosis in mice"

### Supplementary Figures

#### ***LDLr$^{-/-}$ mice***

**2-wk duration treatment** Mice: LDLr$^{-/-}$, female or male, 4–9 mice per group
Diets: HFD = TD.94059 Atherogenic Diet (37% fat, 1.25% cholesterol)
CHOW = LabDiet 5053 (13% fat)
Treatment: variable TFA dose and route of administration

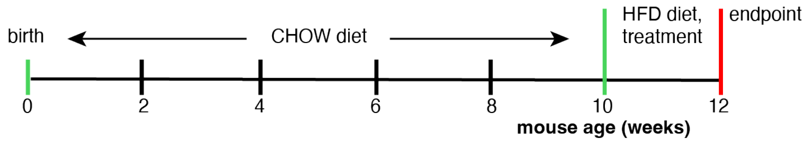

**10-wk duration treatment** Mice: LDLr$^{-/-}$, female, 8 mice per group
Diets: HFD = TD.94059 Atherogenic Diet (37% fat, 1.25% cholesterol)
CHOW = LabDiet 5053 (13% fat)
Treatment: Daily 200 μmol/kg TFA or PBS vehicle, oral in drinking water

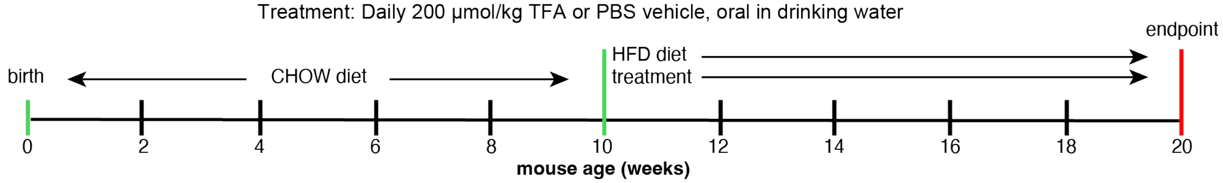

#### ***apoE$^{-/-}$ mice***

**10-wk duration treatment** Mice: apoE$^{-/-}$, female or male, 10 mice per group
Diets: HFD = TD.88137 Adjusted Calories Diet (42% fat, 0.2% cholesterol)
CHOW = LabDiet 5053 (13% fat)
Treatment: Daily 200 μmol/kg TFA or PBS vehicle, oral in drinking water

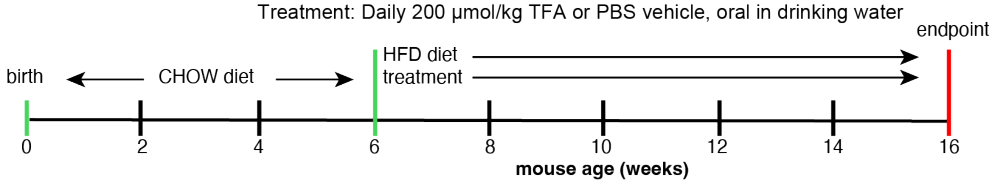

#### ***wild-type C57Bl mice***

**5-wk duration treatment** Mice: male C57BL/6J DIO (stock # 380050) or C57BL/6J DIO control mice (stock # 380056), 10 mice/group
Diets: HFD = Research Diets D12492 (60% fat)
CHOW = Research Diets D12450J (10% fat)
Treatment: Daily 130 μmol/kg TFA or PBS vehicle, oral gavage

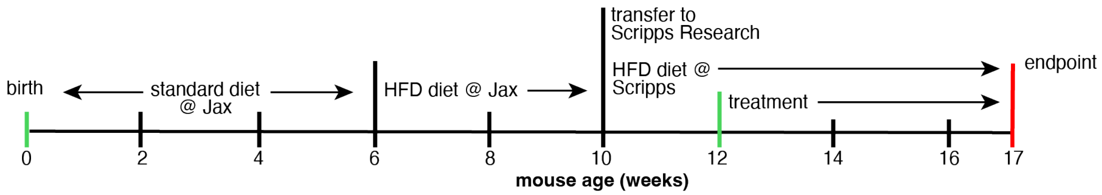

**Figure S1.** Schedules of feeding and dosing for the described studies.

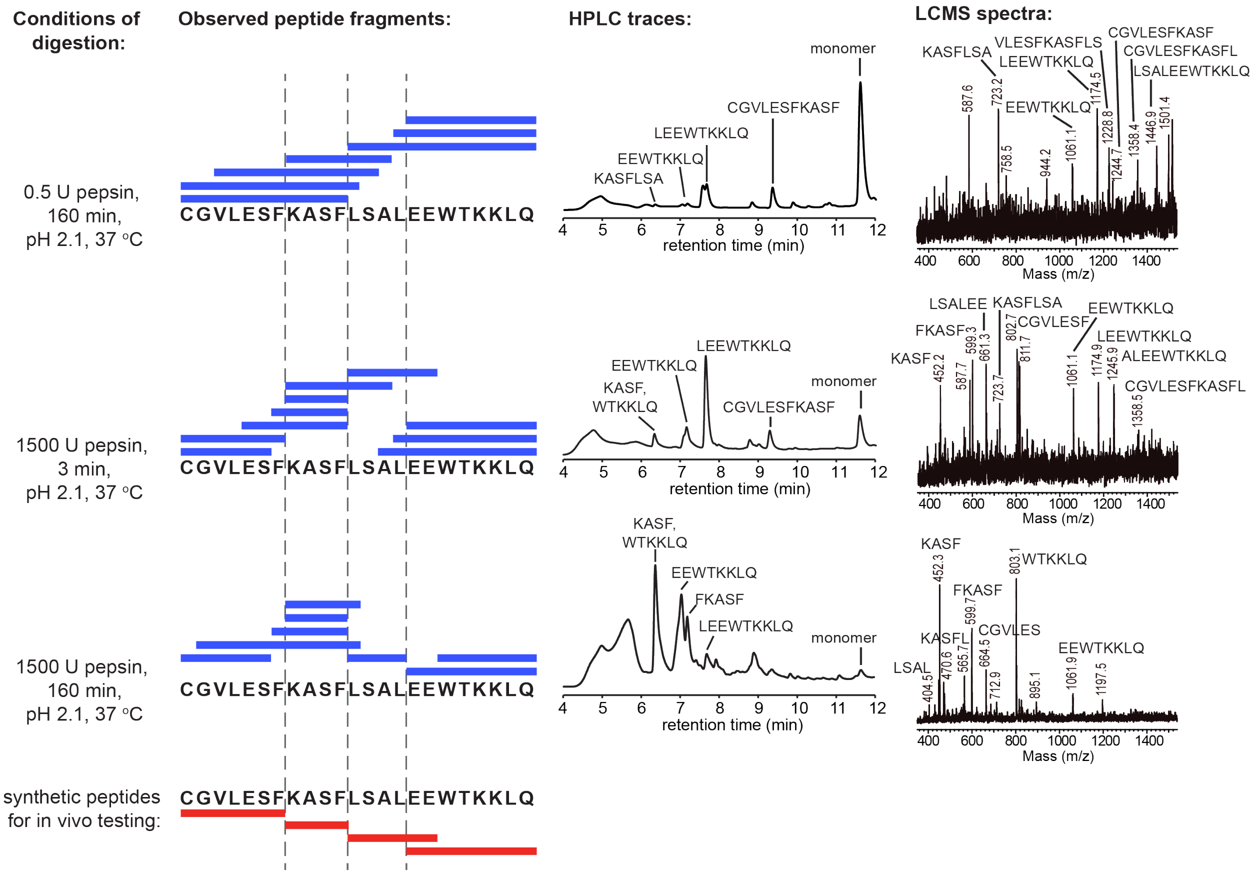

**Figure S2.** To determine which peptide fragments might be produced following oral administration of the 23-residue parent apoA-I mimetic peptide, in vitro digestions with pepsin were carried out at pH 2, 37° C. The three digestions involved increasing enzyme activity to approximate early, middle, and late peptide fragments that might be formed in vivo. The digestion reactions were quenched using pepstatin, and the digests were analyzed by LCMS analysis. Based on our analysis of the observed peptide fragments produced in the digests (blue bars), we chose four peptide fragments to synthesize for follow-up in vivo testing (red bars). The Cys residue in the parent peptide was alkylated with iodoacetamide, resulting in a residue mass of 161.04 Da. It should be noted that fragments containing a Trp residue will have a higher extinction coefficient than those without Trp, resulting in higher UV absorbance in HPLC traces.

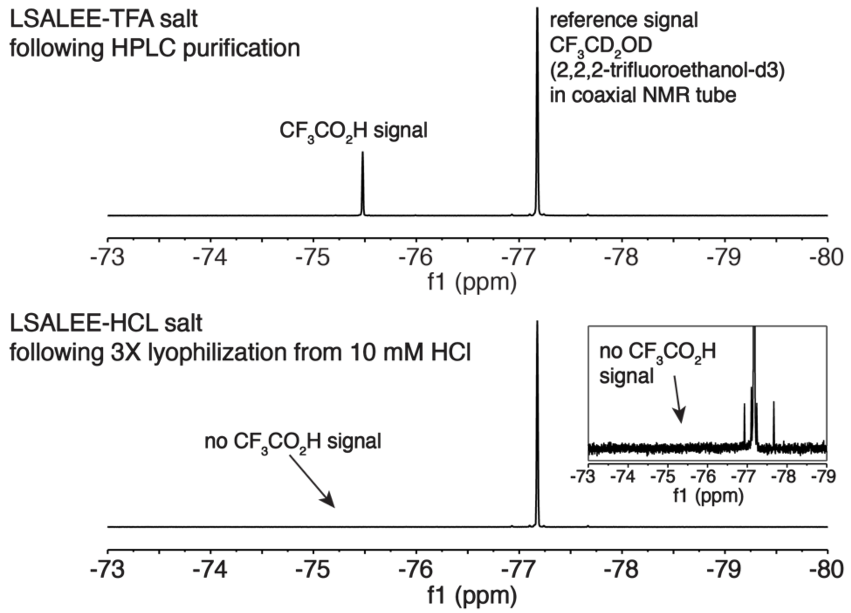

**Figure S3.** $^{19}$F-NMR analysis confirming complete removal of TFA in hexapeptide LSALEE.

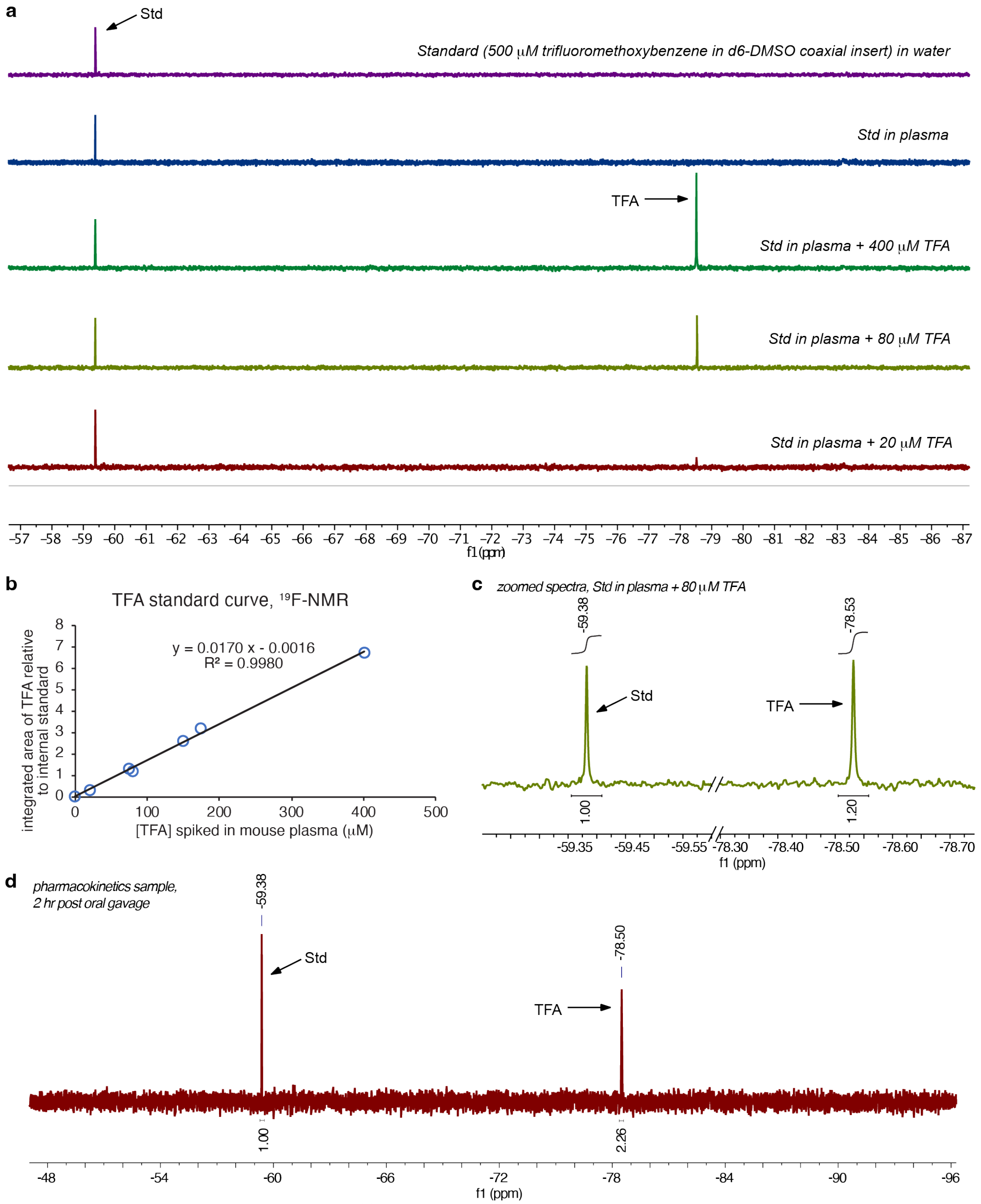

**Figure S4.** Development of the $^{19}$F-NMR assay to measure TFA concentration in plasma of treated mice. Plasma samples were diluted to 500 µL total volume and placed in an NMR tube along with a coaxial tube insert containing 500 µM trifluoromethoxybenzene in d6-DMSO as an internal concentration standard **a**) $^{19}$F-NMR spectra of plasma from untreated mice spiked with known concentrations of TFA, used to generate a standard curve. **b**) The standard curve for measuring TFA in plasma samples from TFA-treated mice. **c**) Zoomed $^{19}$F-NMR spectra of the peaks for Std and TFA in the plasma sample spiked with 80 µM TFA. **d**) $^{19}$F-NMR spectra of plasma drawn from a mouse two hours after oral gavage with a 200 µmol/kg dose of TFA.

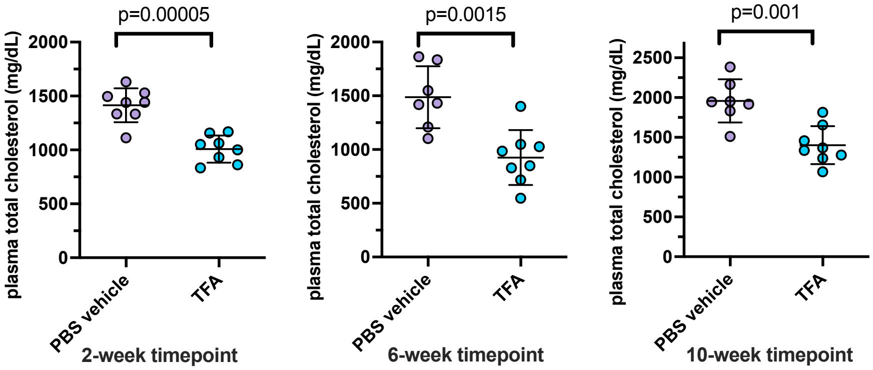

**Figure S5.** Plasma cholesterol levels in female LDLr$^{-/-}$ mice throughout a 10-week anti-atherosclerosis study. TFA was administered treatment *ad libitum* in the drinking water (200 μmol/kg/day). The data are shown as mean ± SD. *p* values were determined by one-way ANOVA comparing the experimental group to PBS vehicle group.

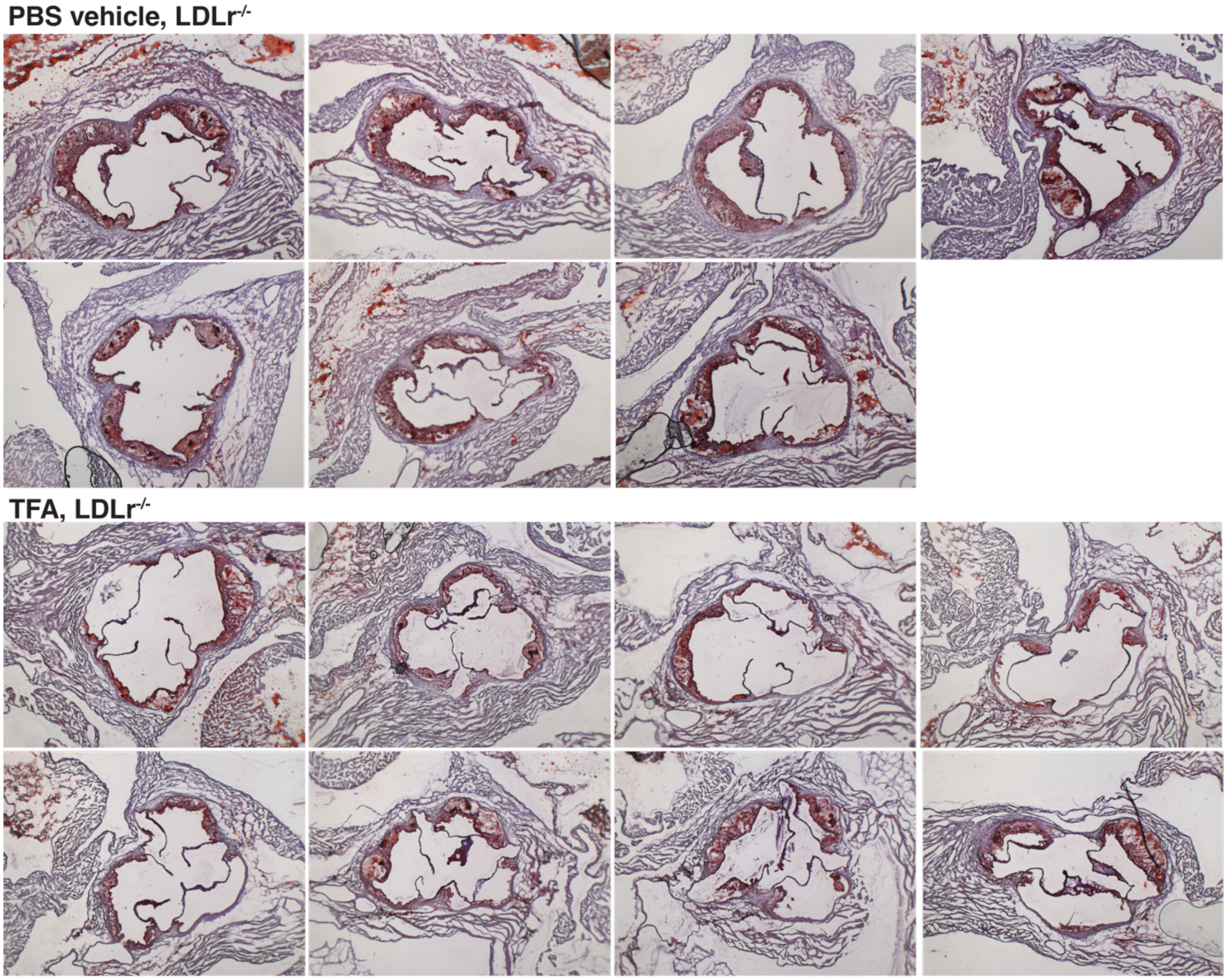

**Figure S6.** Aortic sinus cross-sections from TFA- and PBS vehicle-treated HFD-fed LDLr$^{-/-}$ mice (female) following ten-week treatment *ad libitum* in the drinking water (200 μmol/kg/day). The lipid is stained red.

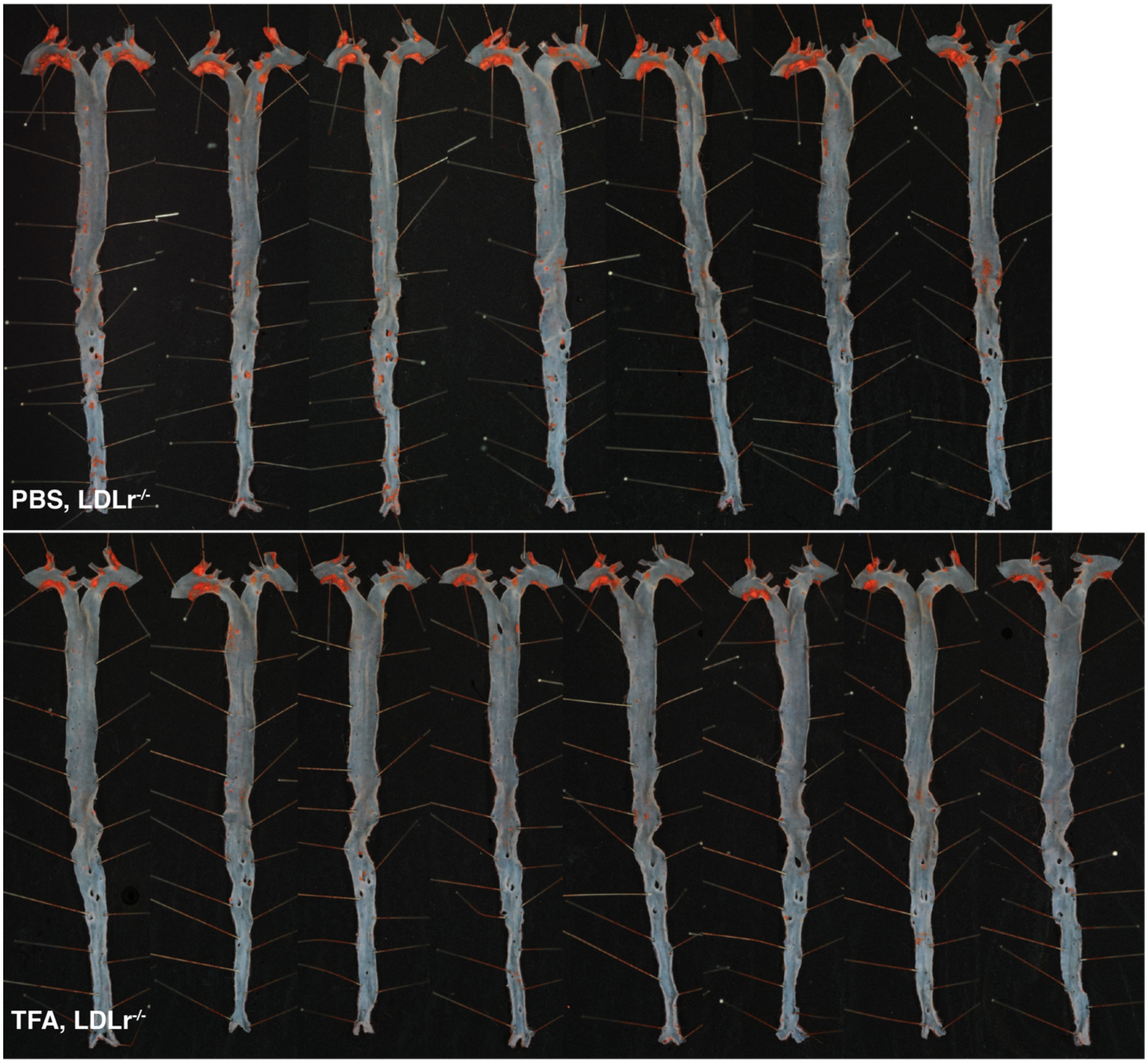

**Figure S7.** Aortas from TFA- and PBS vehicle-treated HFD-fed LDLr$^{-/-}$ mice (female) following ten-week treatment *ad libitum* in the drinking water (200 μmol/kg/day). The lipid is stained red.

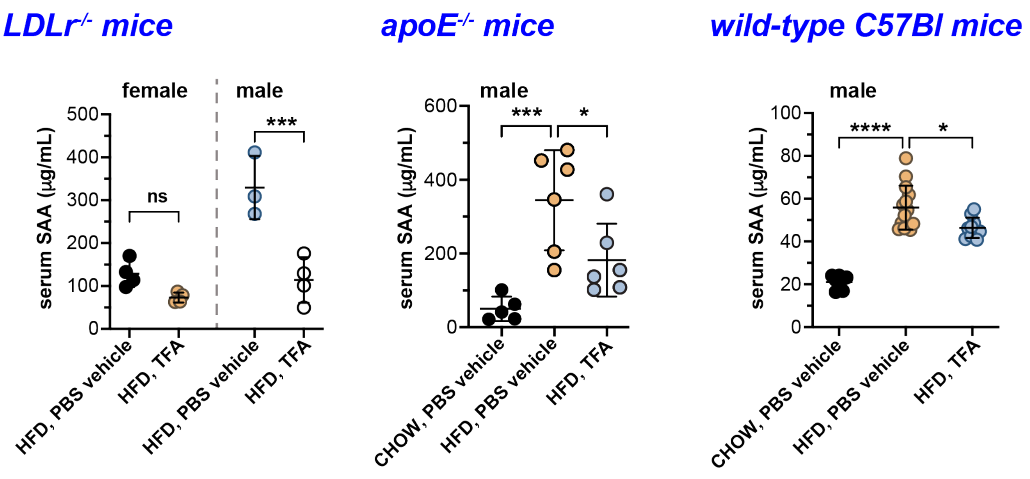

**Figure S8.** Serum SAA levels were lowered in LDLr$^{-/-}$, apoE$^{-/-}$, and wild-type C57Bl mice following TFA treatment. For LDLr$^{-/-}$ mice, plasma was taken following two weeks of a daily i.p. injection of TFA (200 µmol/kg). For wild-type C57Bl mice (male), plasma was taken following four weeks of a daily oral gavage of TFA (130 µmol/kg). The data are shown as mean ± SD. *p* values were determined by one-way ANOVA comparing the experimental group to PBS vehicle group; ns, not significantly different; **, $p \leq 0.01$; ***, $p \leq 0.001$; ****, $p \leq 0.0001$.

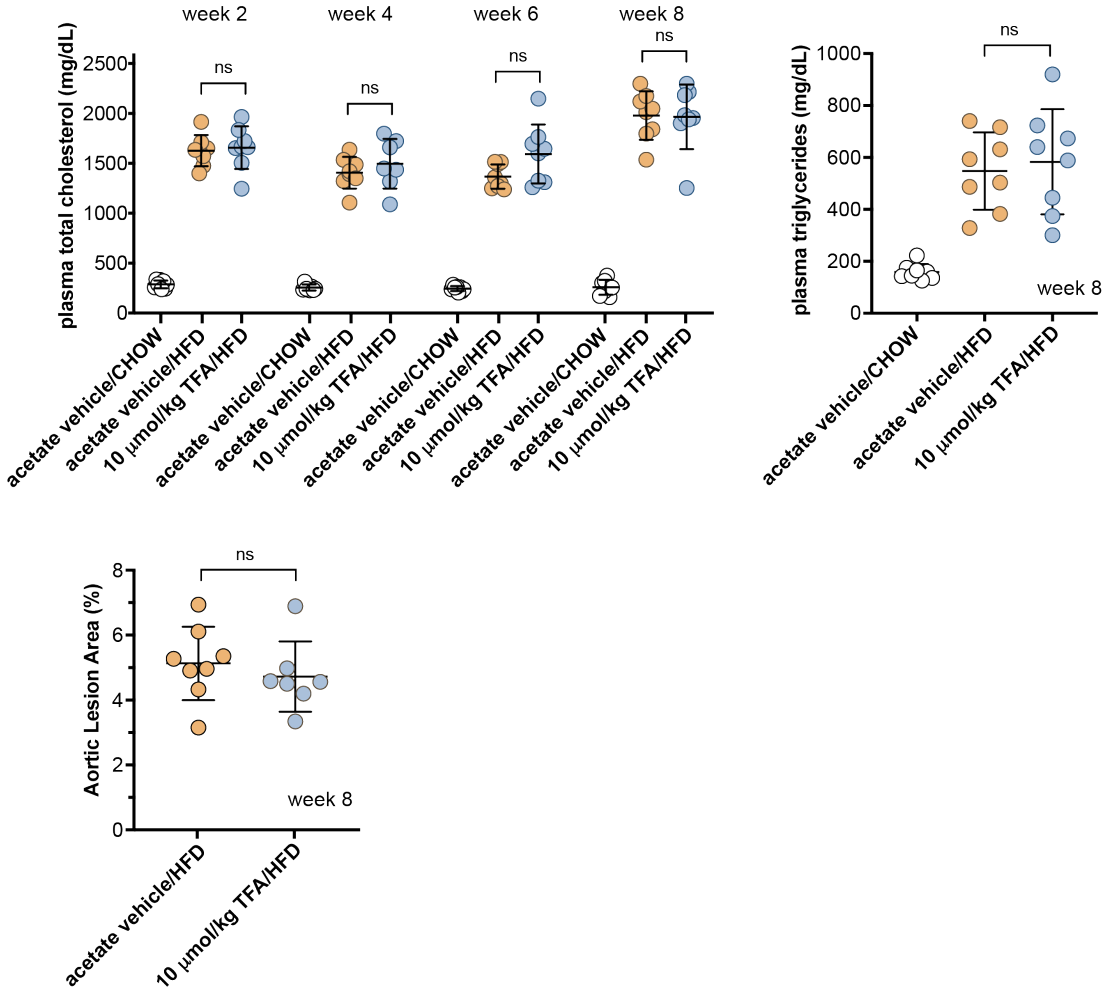

**Figure S9.** In an 8-wk atheroprotection study using a low dose of TFA (10 µmol/kg/day) with an i.p. route of administration in HFD-fed female LDLr$^{-/-}$ mice, TFA treatment did not cause changes in plasma cholesterol or triglyceride levels, nor did it reduce the development of aortic lesions. The vehicle in this study was 10 mM acetate buffer, pH 4. The data are shown as mean ± SD. *p* values were determined by one-way ANOVA comparing the experimental group to PBS vehicle group; ns, not significantly different.

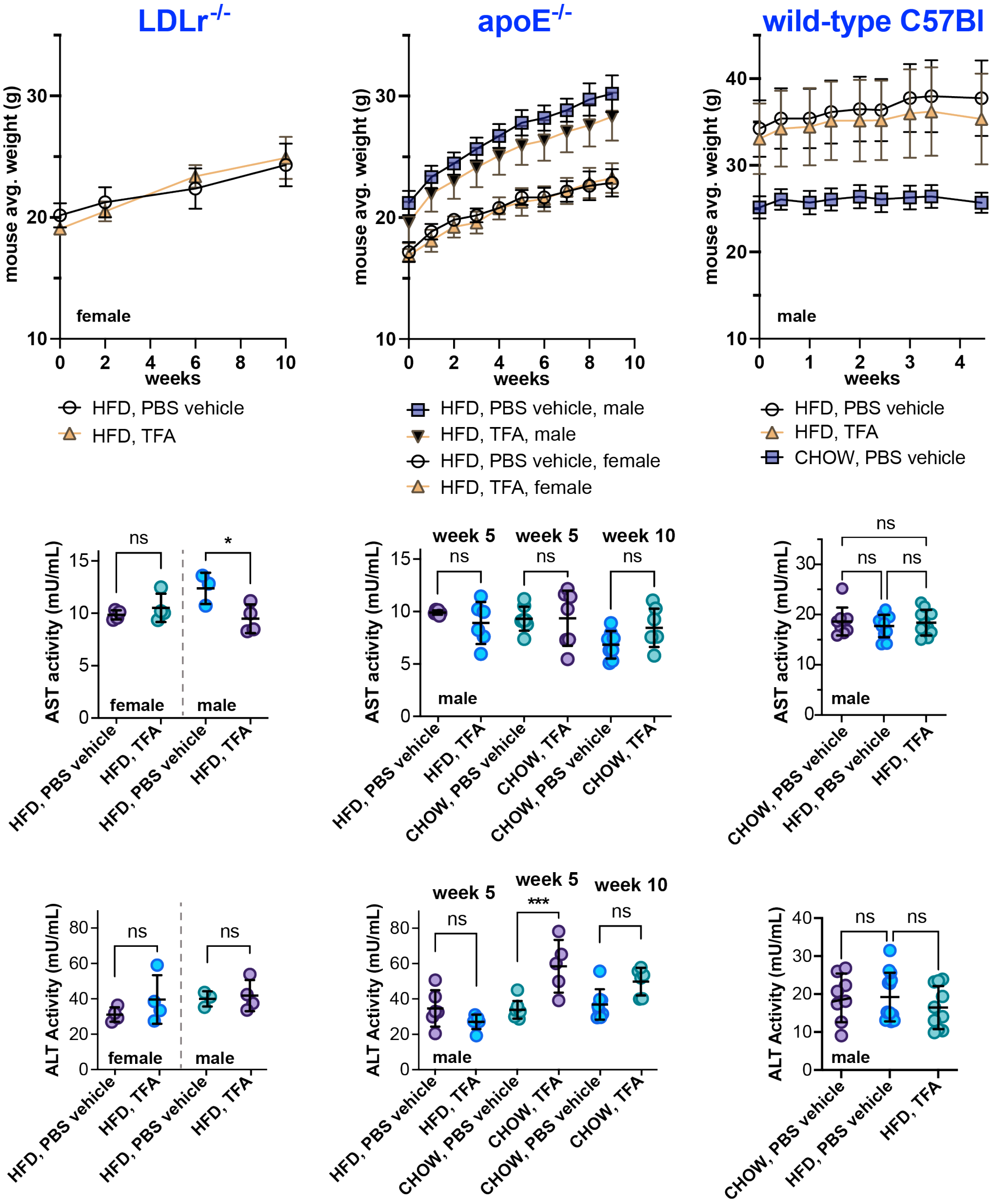

**Figure S10.** TFA treatment did not affect weight gain compared to vehicle-treated control animals over the course of multi-week studies, nor did TFA cause changes in the plasma levels of liver enzymes alanine transaminase (ALT) or aspartate aminotransferase (AST). The data are shown as mean ± SD. *p* values were determined by one-way ANOVA comparing the experimental group to PBS vehicle group; ns, not significantly different; **, $p \leq 0.01$; ***, $p \leq 0.001$; ****, $p \leq 0.0001$.

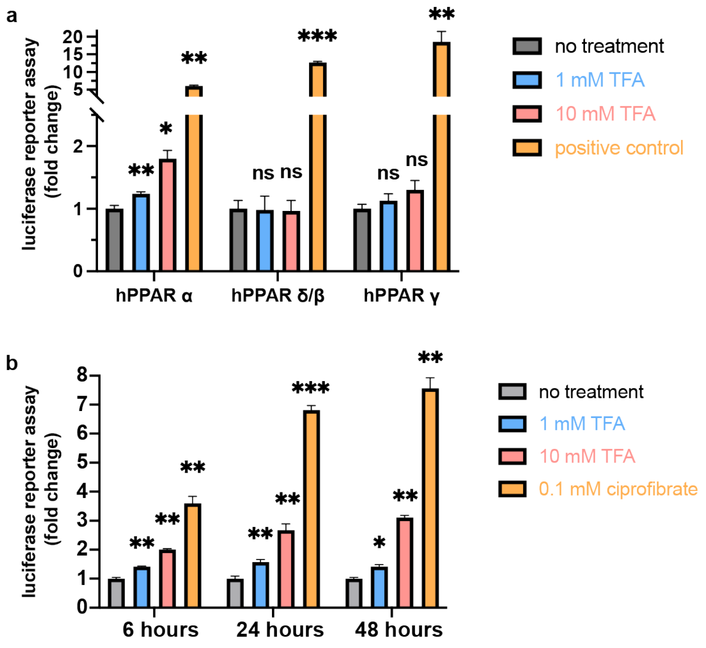

**Figure S11.** **a**) Reporter assay in human HepG2 liver cells transfected with different human PPAR isoforms, in which luciferase expression is driven by activation of the PPAR response element. A different positive control was used for each PPAR isoform: 0.1 mM ciprofibrate for PPAR-α, 1 μM GW501516 for PPAR-δ/β, and 10 μM rosiglitazone for PPAR-γ. **b**) Reporter assay in human HepG2 liver cells transfected with mouse PPAR-α, showing the effect of different TFA treatment times. The data are shown as mean ± SD. *p* values were determined by one-way ANOVA comparing the experimental group to PBS vehicle group; ns, not significantly different; **, $p \leq 0.01$; ***, $p \leq 0.001$; ****, $p \leq 0.0001$.

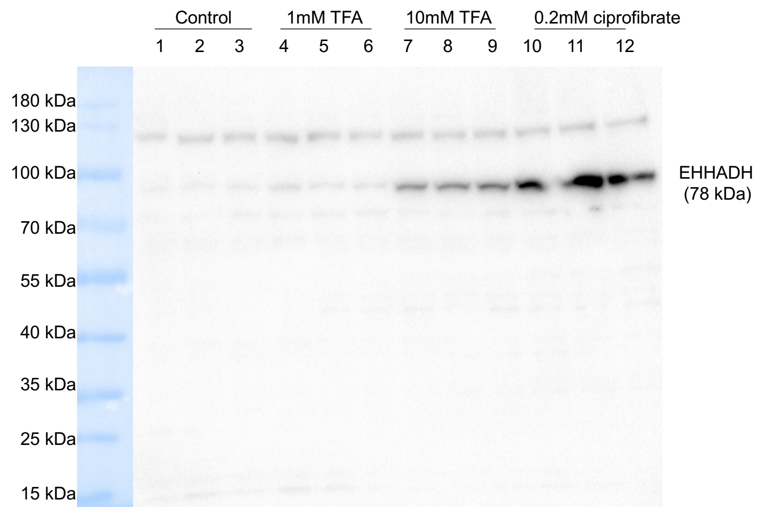

**Figure S12.** Western blot of rat FAO cell lysates showing that TFA treatment led to higher levels of EHHADH protein in the cells.

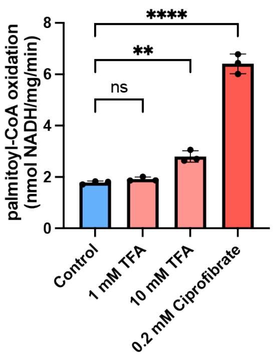

**Figure S13.** Palmitoyl-CoA oxidase activity in rat FAO liver cells following treatment with TFA or ciprofibrate. The data are shown as mean ± SD. *p* values were determined by one-way ANOVA comparing the experimental group to PBS vehicle group; ns, not significantly different; **, $p \leq 0.01$; ***, $p \leq 0.001$; ****, $p \leq 0.0001$.

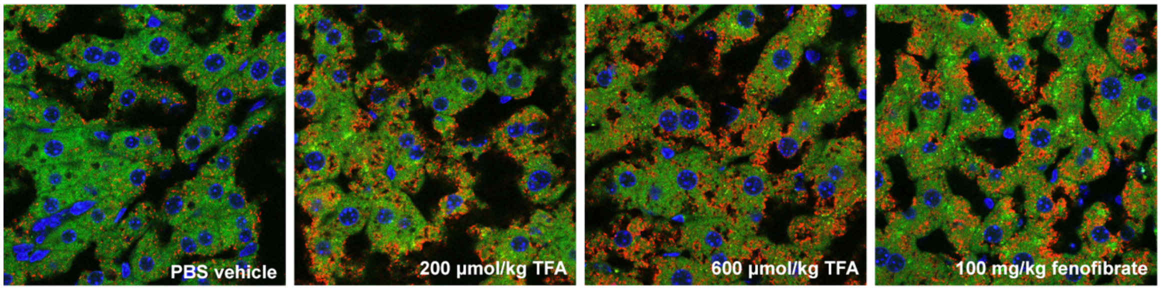

**Figure S14.** Confocal microscopy at 60x magnification of liver cells from the TFA- or vehicle-treated LDLr$^{-/-}$ animals, stained red for PMP70, a marker of peroxisome proliferation. Red, AF568 stain for PMP70; blue, DAPI stain for nuclei; green, autofluorescence.

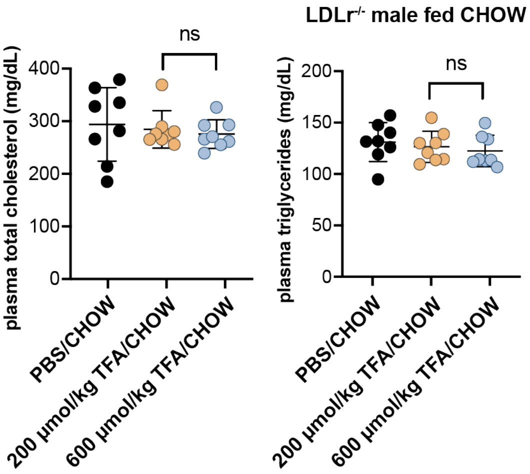

**Figure S15.** No changes were observed in triglyceride or cholesterol levels of male LDLr$^{-/-}$ mice fed a CHOW diet when treated with TFA for two weeks by daily oral gavage. The data are shown as mean ± SD. *p* values were determined by one-way ANOVA comparing the experimental group to PBS vehicle group; ns, not significantly different.

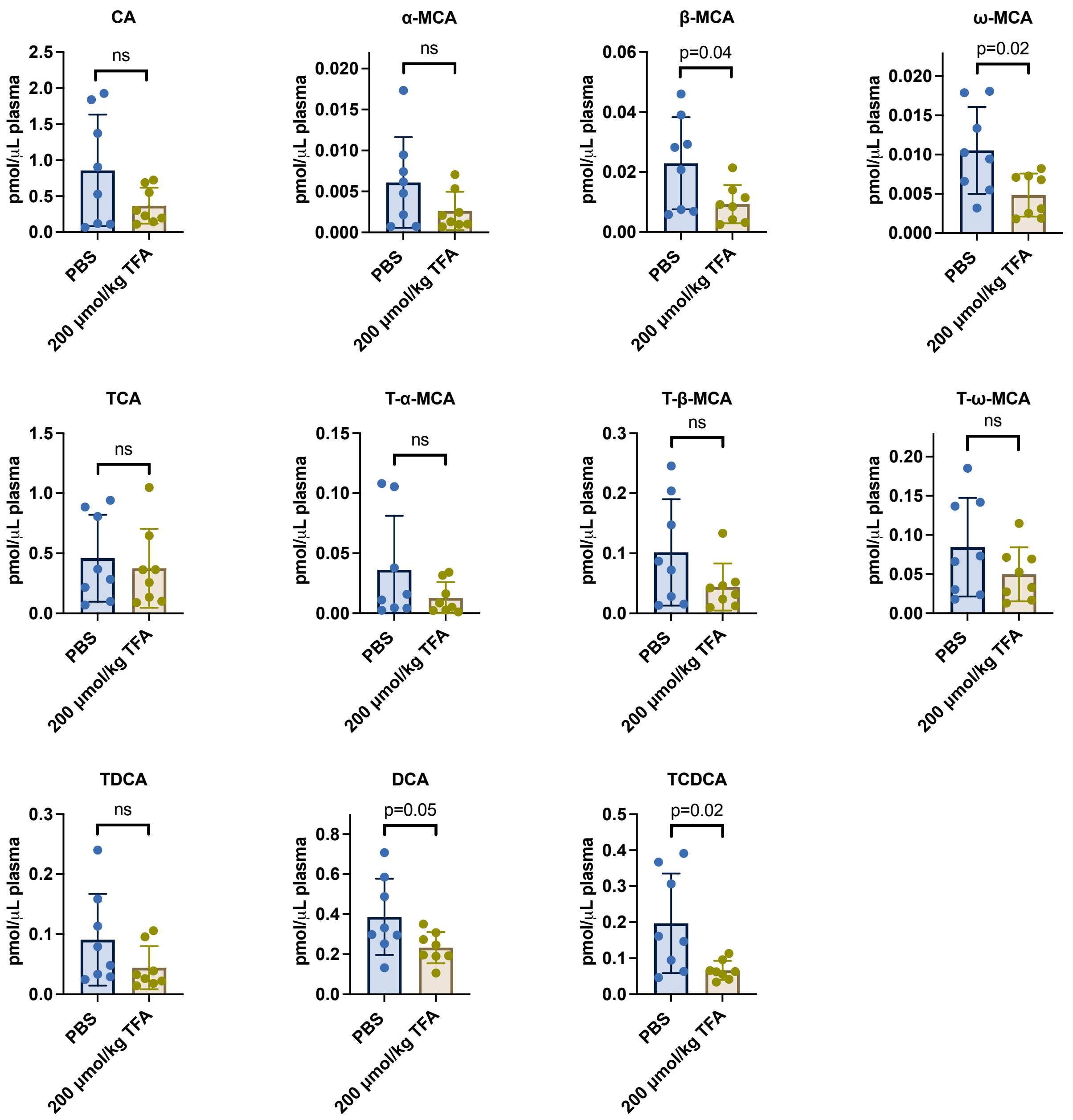

**Figure S16.** Observed bile acid levels in the plasma of male LDL-null mice (n=8 per group) following two-week administration of TFA (200 µmol/kg/day) or PBS vehicle in the drinking water. Bile acid levels were determined by targeted mass spectrometry. The data are shown as mean ± SD. *p* values were determined by one-way ANOVA comparing the experimental group to PBS vehicle group; ns, not significantly different.

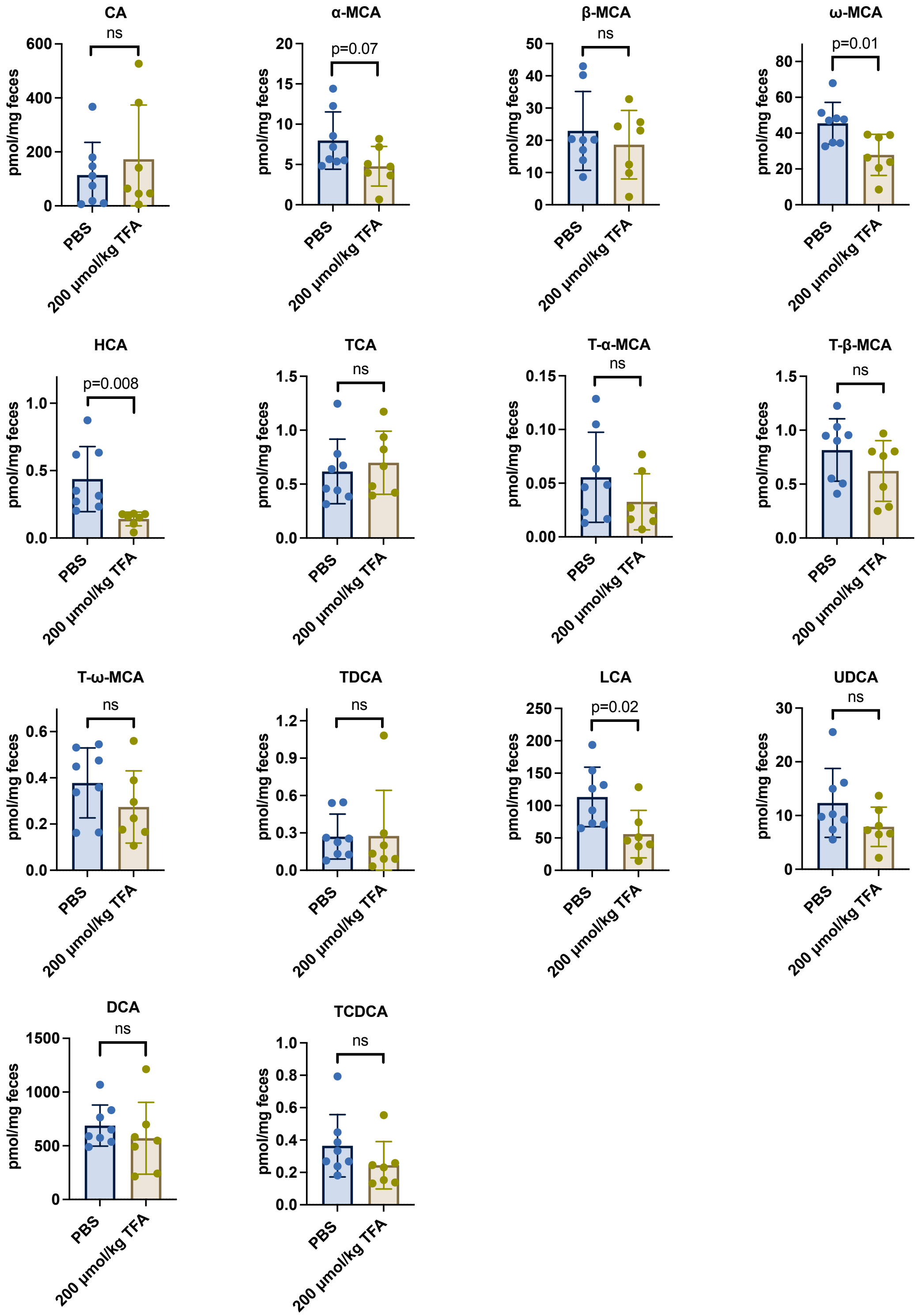

**Figure S17.** Observed bile acid levels in the feces of male LDL-null mice (n=7 or 8 per group) following two-week administration of TFA (200 µmol/kg/day) or PBS vehicle in the drinking water. Bile acid levels were determined by targeted mass spectrometry. The data are shown as mean ± SD. *p* values were determined by one-way ANOVA comparing the experimental group to PBS vehicle group; ns, not significantly different.

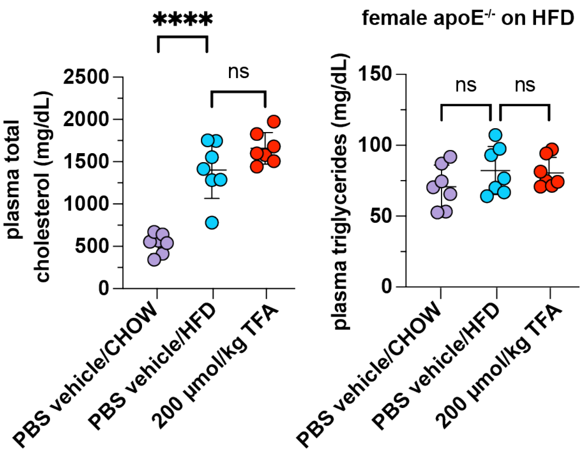

**Figure S18.** No changes were observed in triglyceride or cholesterol levels of apoE$^{-/-}$ mice following TFA treatment. The data are shown as mean ± SD. $p$ values were determined by one-way ANOVA comparing the experimental group to PBS vehicle group; ns, not significantly different; **, $p \leq 0.01$; ***, $p \leq 0.001$; ****, $p \leq 0.0001$.

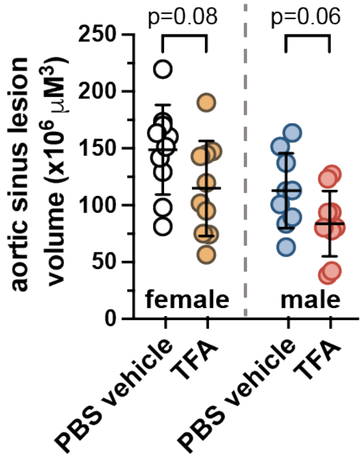

**Figure S19.** Aortic sinus atherosclerotic lesions were reduced, albeit not to a statistically significant degree, by daily administration of TFA (200 µmol/kg) in the drinking water for 10 weeks in HFD-fed apoE$^{-/-}$ mice. The data are shown as mean ± SD. $p$ values were determined by one-way ANOVA comparing the experimental group to PBS vehicle group.

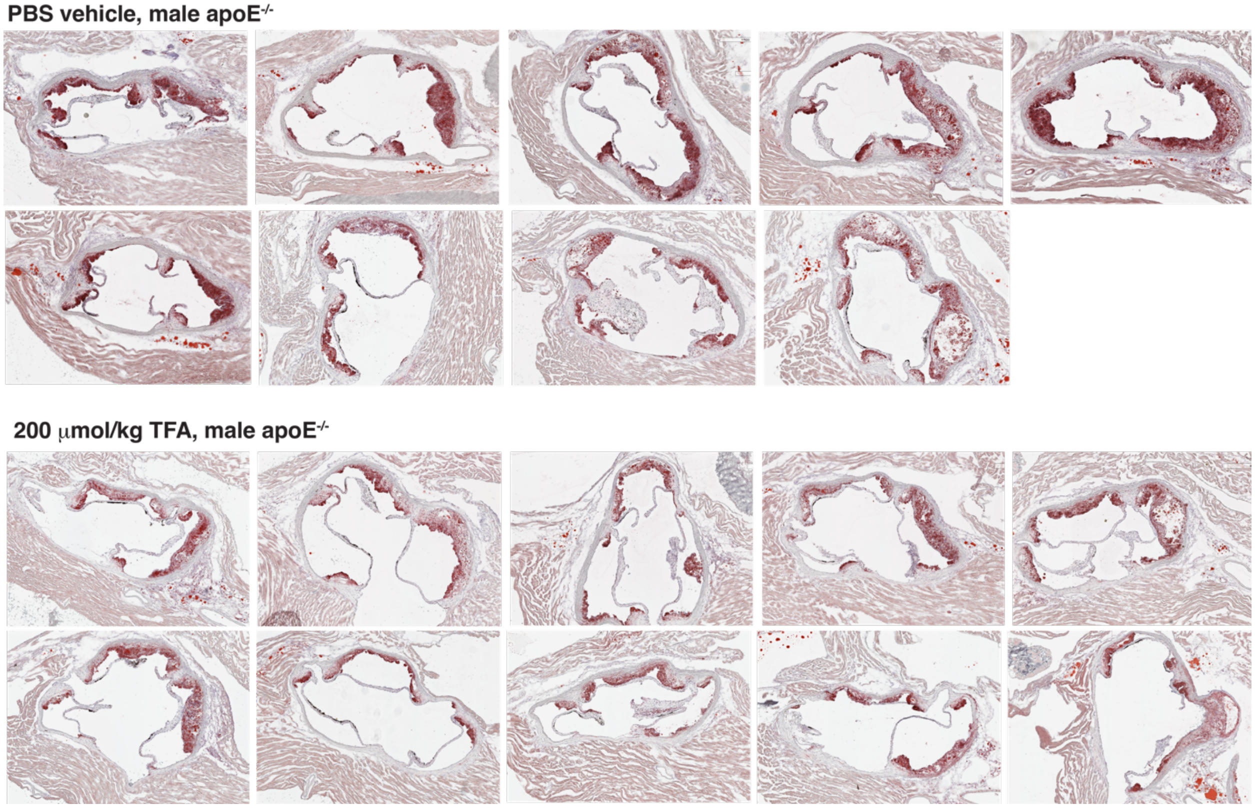

**Figure S20.** Aortic sinus cross-sections from male TFA- and PBS vehicle-treated HFD-fed apoE$^{-/-}$ mice following ten-week treatment *ad libitum* in the drinking water (200 μmol/kg/day). The lipid is stained red.

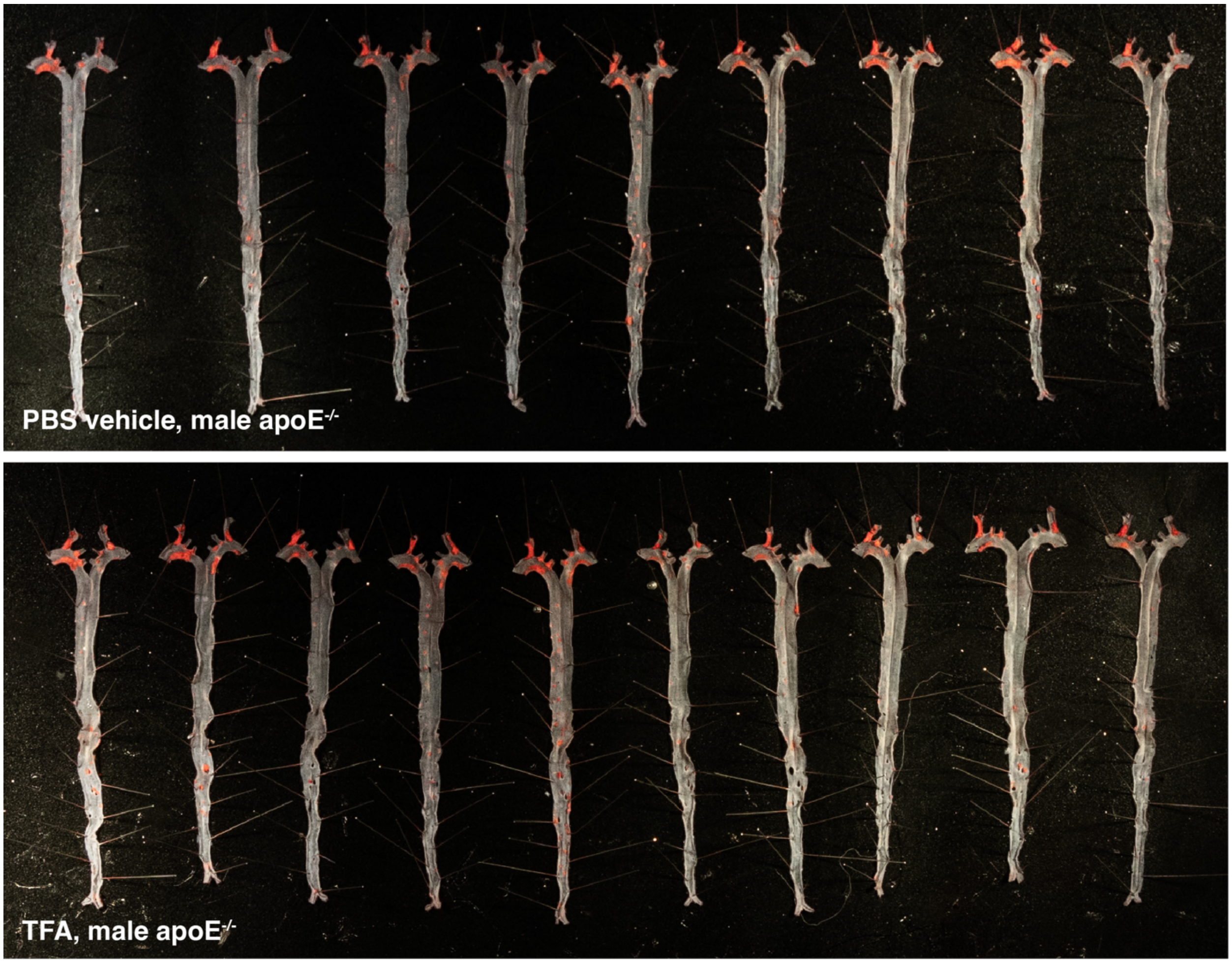

**Figure S21.** Aortas from male TFA- and PBS vehicle-treated HFD-fed apoE$^{-/-}$ mice following ten-week treatment *ad libitum* in the drinking water (200 μmol/kg/day). The lipid is stained red.

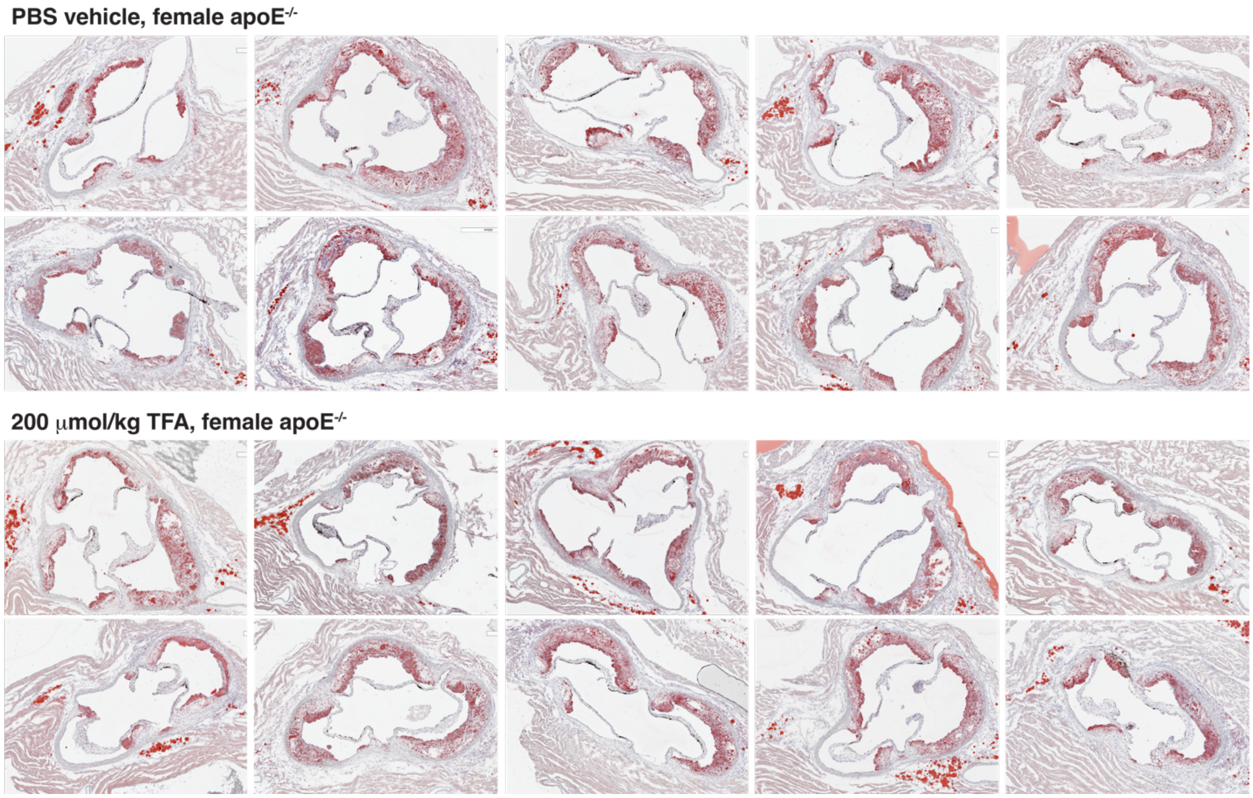

**Figure S22.** Aortic sinus cross-sections from female TFA- and PBS vehicle-treated HFD-fed apoE$^{-/-}$ mice following ten-week treatment *ad libitum* in the drinking water (200 µmol/kg/day). The lipid is stained red.

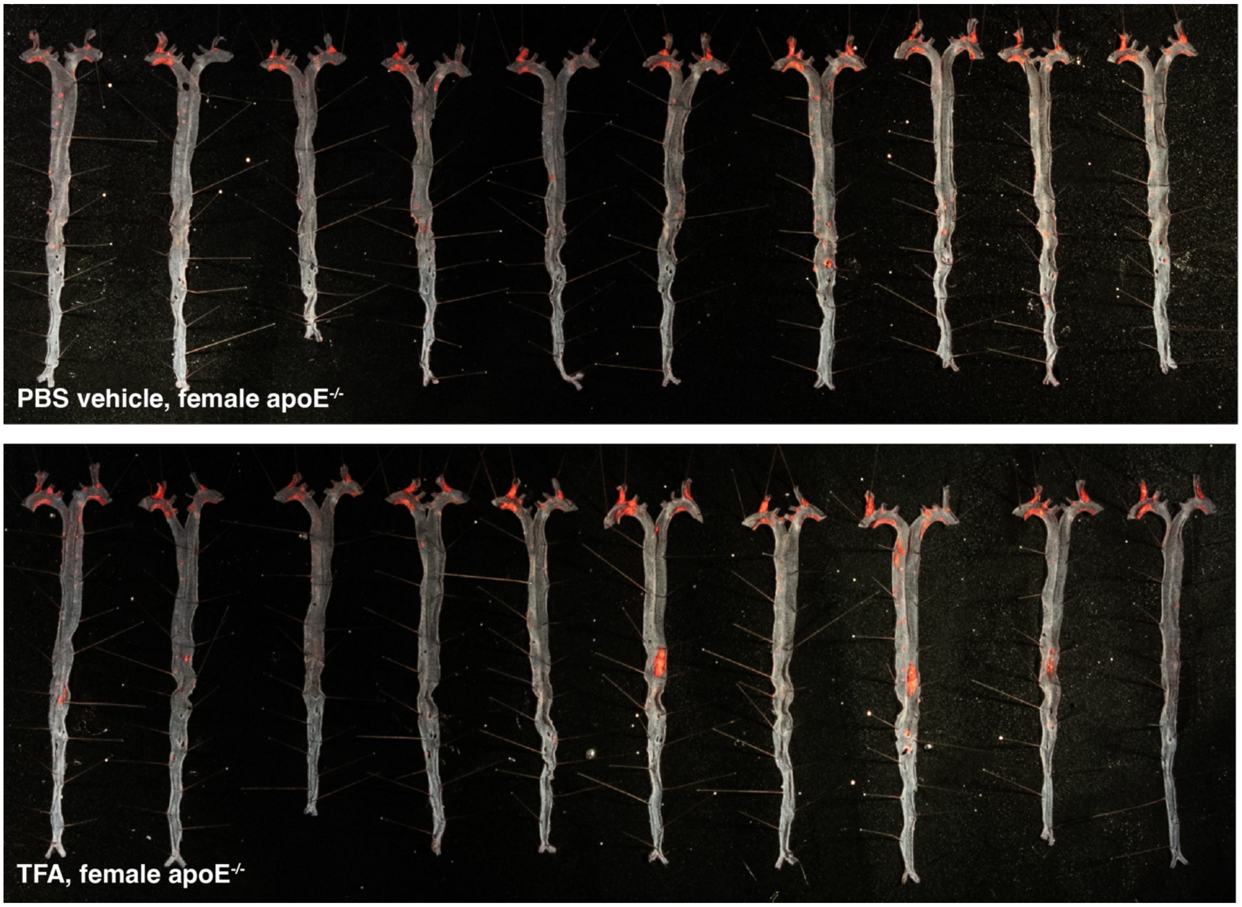

**Figure S23.** Aortas from female TFA- and PBS vehicle-treated HFD-fed LDLr$^{-/-}$ mice following ten-week treatment *ad libitum* in the drinking water (200 µmol/kg/day). The lipid is stained red.

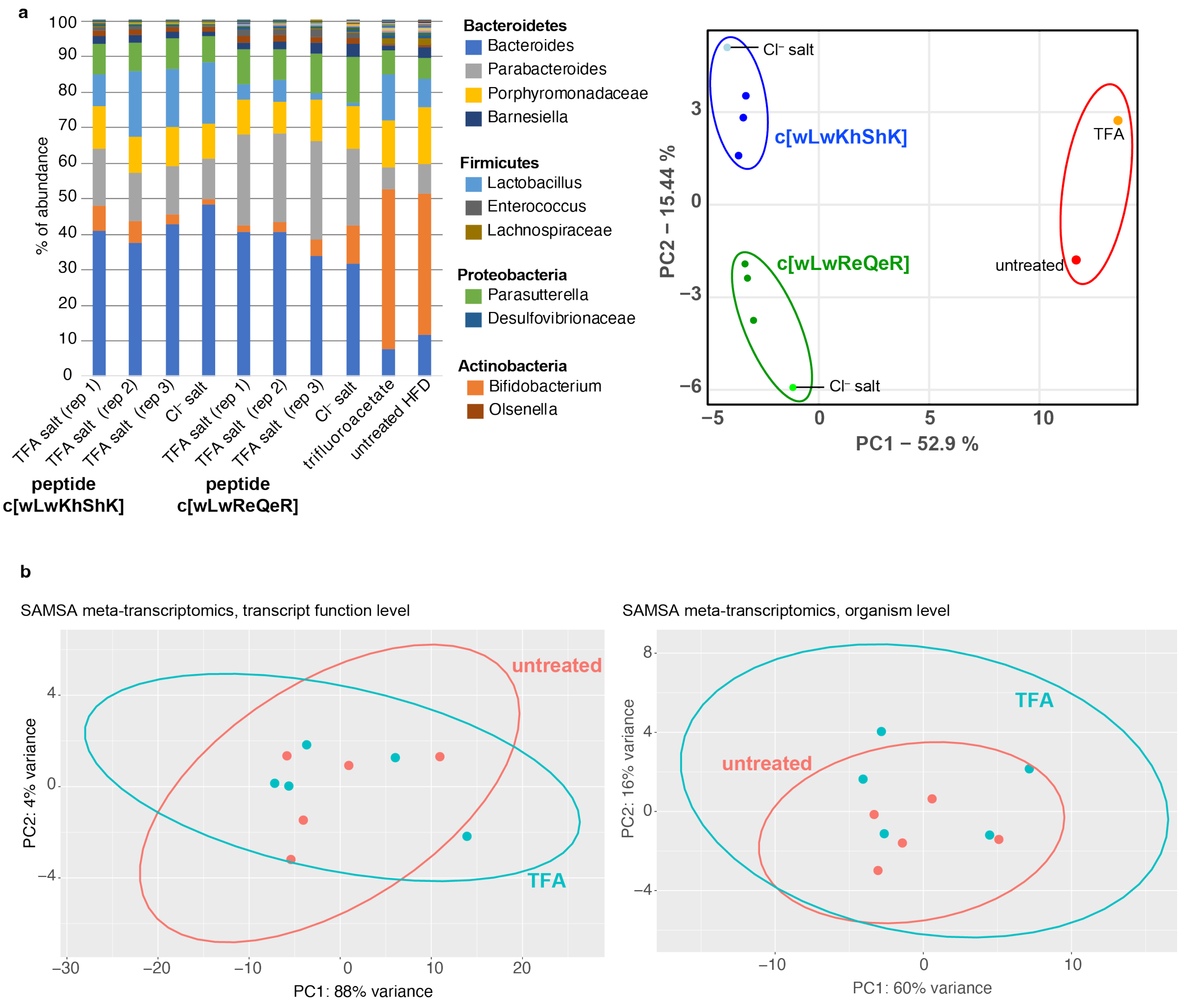

**Figure S24.** In vitro analyses of TFA effects on gut bacteria. **a)** Fresh gut microbiome content was isolated by surgically removing cecum under anaerobic conditions from *LDLr*$^{-/-}$ mice that had been fed HFD for two weeks. The cecal contents were then mixed with growth media, distributed in 96-well plates, and incubated overnight in the presence of TFA (1 mM) or positive control gut bacteria remodeling cyclic peptides c[wLwReQeR] or c[wLwKhShK] (64 µM). The peptides were tested either as TFA salt (three replicates) or Cl- salt. After the incubation, the composition of the gut microbiota community was assessed *en masse* by 16S rRNA sequencing of each well. As indicated by bacterial genera composition or Bray-Curtis beta diversity principal component analysis, the peptides had remodeled the gut bacteria community, whereas TFA did not appreciably remodel the community. **b)** To assess the possible effect of TFA on gut bacteria meta-transcriptomics, fresh microbiome content was isolated as described in part **a**. The cecal contents were mixed with growth media and incubated with TFA (1 mM) or mock buffer (n=5/group) for six hours. Total RNA was isolated. Library preparation and sequencing were carried out by Novogene. The data were analyzed using SAMSA2 on the transcript function level or organism level. No differences were observed between TFA treatment vs untreated samples.

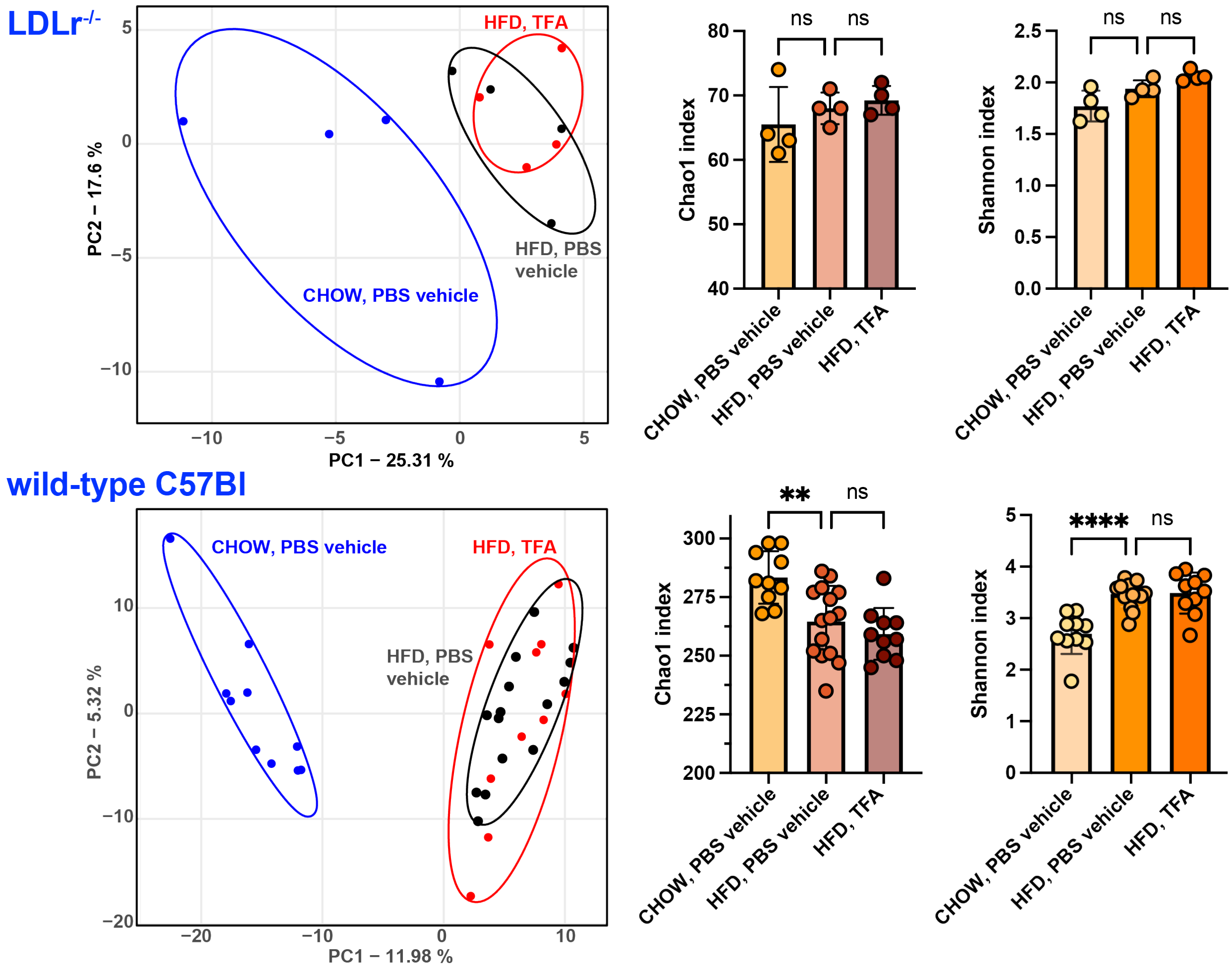

**Figure S25.** Analysis of 16S gut microbiome sequencing of fecal samples from PBS vehicle- or TFA-treated mice. Fecal samples were from LDLr$^{-/-}$ mice (two-week treatment, 200 μmol/kg/day, n=4/group) or wild-type C57Bl mice (four-week treatment, 200 μmol/kg/day, n=10-15/group). On the left is shown a Bray-Curtis principal component analysis of beta diversity between samples. On the right, plots of alpha diversity are shown. The data are shown as mean ± SD. *p* values were determined by one-way ANOVA comparing the experimental group to PBS vehicle group; ns, not significantly different; **, $p \leq 0.01$; ***, $p \leq 0.001$; ****, $p \leq 0.0001$.

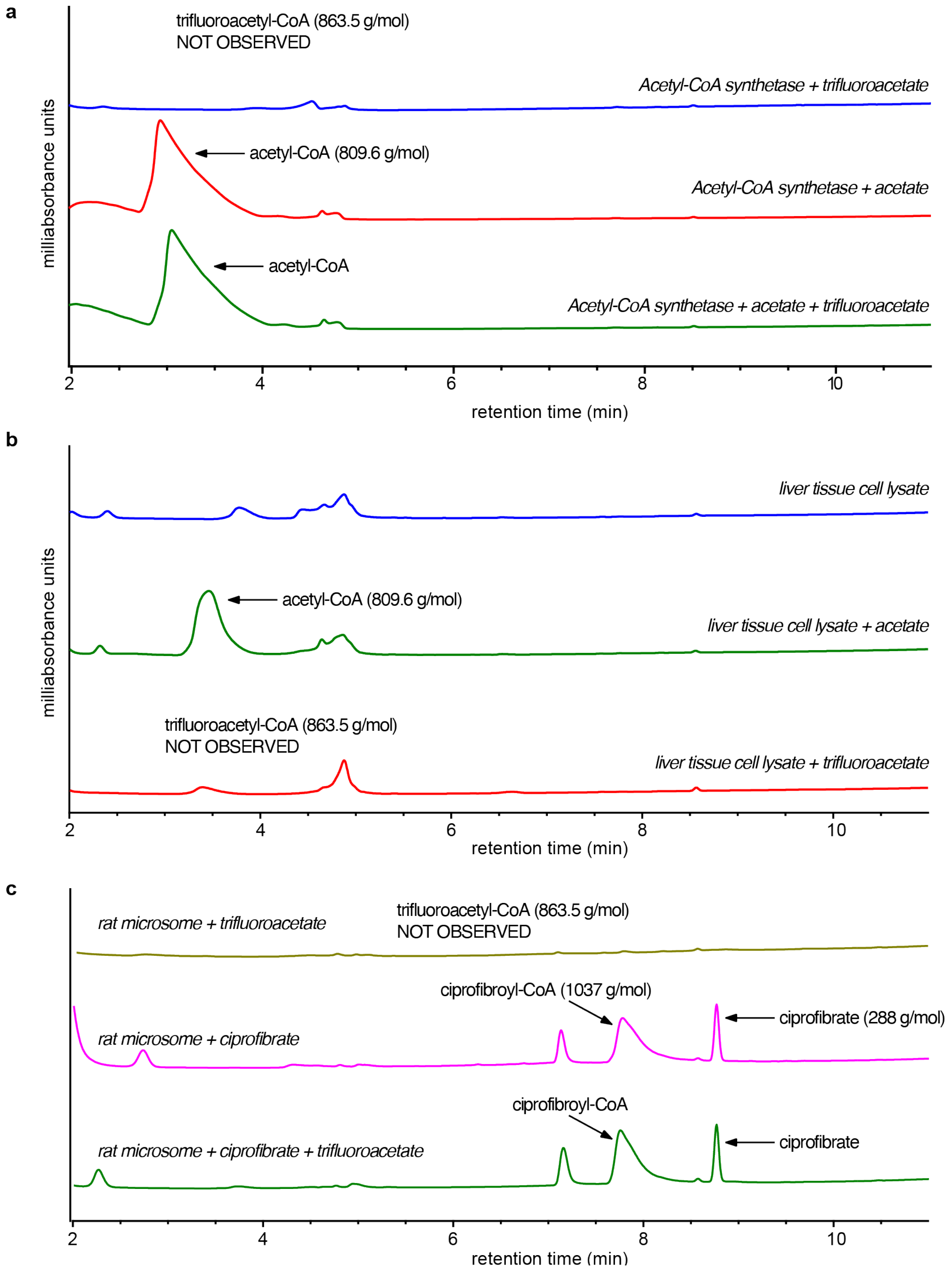

**Figure S26.** Biochemical experiments using purified acetyl-CoA synthetase (**a**), rat liver tissue lysate (**b**), or rat liver microsomes (**c**) suggest that trifluoroacetyl-CoA is not produced. Whereas acetyl-CoA and ciprofibroyl-CoA were observed under analogous conditions by LCMS analysis, trifluoroacetyl-CoA was not. Furthermore, the presence of TFA did not apparently affect the formation of acetyl-CoA or ciprofibroyl-CoA.

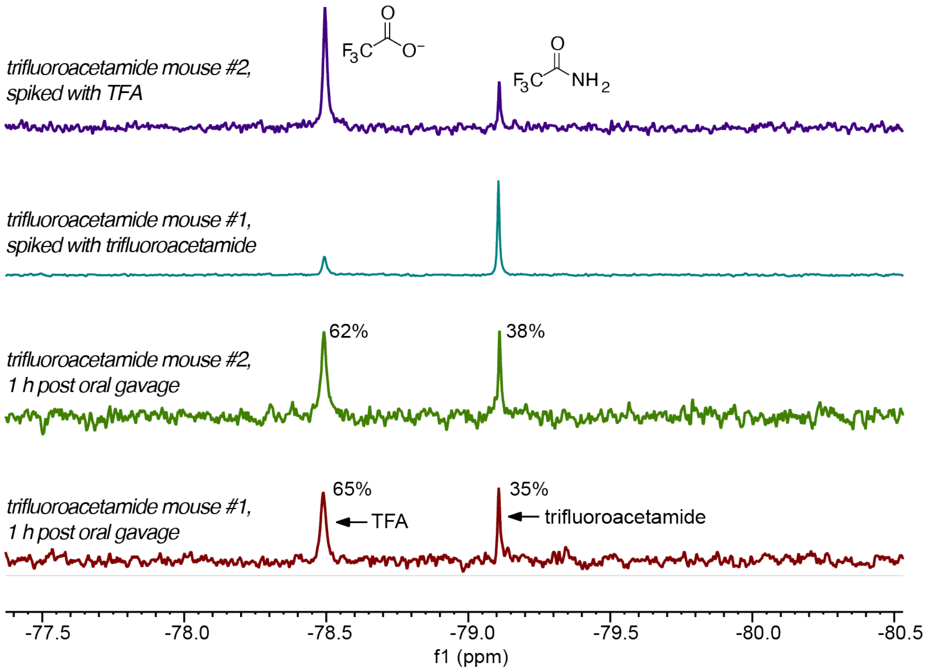

**Figure S27.** 2,2,2-Trifluoroacetamide is converted into TFA in vivo following oral administration. Mice were gavaged with a 200 µmol/kg dose of 2,2,2-trifluoroacetamide. After 1 h, blood was drawn and analyzed by $^{19}$F-NMR. Around 64% of the trifluoroacetylated species in plasma was TFA. Spiking of the plasma with TFA or trifluoroacetamide confirmed the identity of the observed peaks.

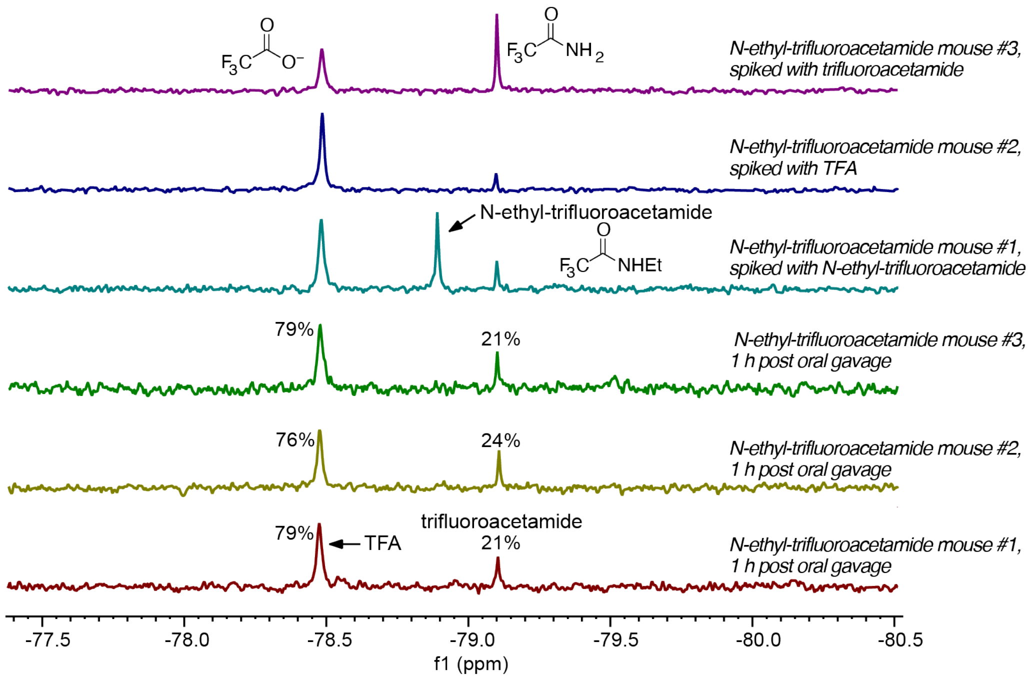

**Figure S28.** N-Ethyl-trifluoroacetamide is converted into TFA and trifluoroacetamide in vivo following oral administration. Mice were gavaged with a 200 µmol/kg dose of N-ethyl-trifluoroacetamide. After 1 h, blood was drawn and analyzed by $^{19}$F-NMR. Around 78% of the trifluoroacetylated species in plasma was TFA, with the remainder being 2,2,2-trifluoroacetamide. None of the administered N-ethyl-trifluoroacetamide was observed at 1 h post gavage. Spiking of the plasma with TFA, N-ethyl-trifluoroacetamide, or trifluoroacetamide confirmed the identity of the observed peaks.

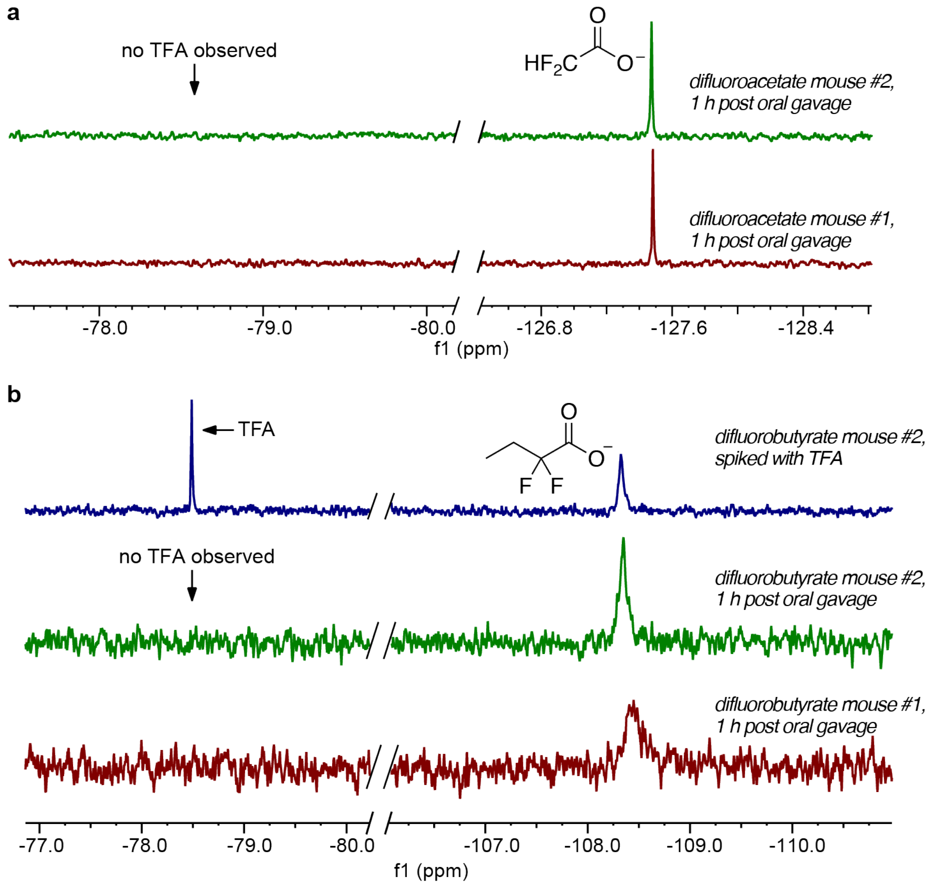

**Figure S29.** Difluoroacetate (**a**) and difluorobutyrate (**b**) are not appreciably converted into TFA over a period of 1 h in vivo following oral administration. Mice were gavaged with a 200 µmol/kg dose of the compound. After 1 h, blood was drawn and analyzed by $^{19}$F-NMR. No TFA was observed in either case at 1 h post gavage. Spiking of the plasma with TFA, difluoroacetate, or difluorobutyrate confirmed the identity of the observed peaks.

### Experimental details

**Abbreviations (not defined below):** ACOX1, acyl CoA oxidase-1; ACADS, acyl CoA dehydrogenase short chain; BSA, bovine serum albumin; DIC, 1,3-diisopropylcarbodiimide; DMEM, Dulbecco's Modified Eagle Medium; FBS, fetal bovine serum; HPLC-MS, high-pressure liquid chromatography-mass spectrometry; MBHA, 4-methylbenzhydrylamine; PBS, phosphate-buffered saline; TIS, triisopropylsilane. LDLr, low-density lipoprotein receptor; apoE, apolipoprotein E; NAD, nicotinamide adenine dinucleotide; NADH, nicotinamide adenine dinucleotide hydrogen; FAD, flavine adenine dinucleotide; SAA, serum amyloid A; CRP, C-reactive protein; CoA, coenzyme A.

**Materials.** Amino acids, Rink amide MBHA resin, and *N*-hydroxybenzotriazole (HOBt) were purchased from Advanced ChemTech, CombiBlocks, or NovaBiochem. Sodium trifluoroacetate was from Alfa-Aesar. N-ethyl-trifluoroacetamide, 2,2-difluorobutyric acid, and 2,2-difluoropropionic acid were from CombiBlocks. 2,2,2-trifluoroacetamide, 5-trifluoromethyltetrazole, and trifluoromethane sulfonamide were from Oakwood. Difluoroacetatic acid and sodium trichloroacetate were from Acros Organics. 3-(trifluoromethyl)-1,2,4-oxadiazol-5-ol was from 1ClickChemistry. Trifluoroacetic acid was obtained from Halocarbon Products (Hackensack, NJ). Antibodies were from Invitrogen or Life Technologies. Other chemicals were purchased from Sigma-Aldrich or Fisher Scientific or as noted below.

**Solid-phase peptide synthesis (SPPS).** Peptides were synthesized using standard Fmoc chemistry with an Advanced Chemtech Apex 396 peptide synthesizer. A typical synthesis was performed on 0.09 mmol scale using 0.6 mmol/g Rink amide MBHA resin (Novabiochem) to prepare a C-terminal amide. Standard side chain protecting groups included Cys(Trt), Gln(Trt), Lys(Boc), Thr(tBu), Trp(Boc), Glu(OtBu), Arg(Pbf), Ser(tBu). Chain elongations were achieved using DIC and HOBt in NMP (*N*-methylpyrrolidinone) with 90-minute couplings. Fmoc deprotection was achieved using 25% piperidine in NMP. Peptides were cleaved from the resin with concomitant side chain deprotection by agitation in a solution of 94:2.5:2.5:1 TFA/ethanedithiol/TIS/water for 3 h. The peptide was precipitated with ether, centrifuged, and washed three additional times with ether. The crude peptides were purified by preparative reverse-phase (RP)-HPLC on a Vydac or Phenomenex reversed phase C18 column. Purity was confirmed by analytical RP-HPLC. Purified peptides were characterized by analytical HPLC-MS and/or MALDI-TOF mass spectrometry. Analytical reverse-phase HPLC was performed using a Zorbax 300-SB C-18 column connected to a Hitachi D-7000 HPLC system. Binary gradients of solvent A (99% $H_2O$, 0.9% acetonitrile, 0.1% TFA) and solvent B (90% acetonitrile, 9.9% $H_2O$, 0.07% TFA) were employed for HPLC.

**Parent 23-residue peptide.** The sequence of parent peptide is H-Cys($CH_2CONH_2$)-Gly-Val-Leu-Glu-Ser-Phe-Lys-Ala-Ser-Phe-Leu-Ser-Ala-Leu-Glu-Glu-Trp-Thr-Lys-Lys-Leu-Gln-$CONH_2$. The N-terminal Cys residue was alkylated with iodoacetamide [22], resulting in a residue mass of 161.04 Da. MW = 2671.9 g/mol.

**FREL.** Peptide sequence H-Phe-Arg-Glu-Leu-COOH. MW = 563.6 g/mol.

**KERS.** Peptide sequence H-Lys-Glu-Arg-Ser-COOH. MW = 518.6 g/mol.

**CGVLESF.** Peptide sequence H-Cys($CH_2CONH_2$)-Gly-Val-Leu-Glu-Ser-Phe-COOH. The N-terminal Cys residue was alkylated with iodoacetamide [22], resulting in a residue mass of 161.04 Da. MW = 811.7 g/mol.

**KASF.** Peptide sequence H-Lys-Ala-Ser-Phe-COOH. MW = 451.5 g/mol.

**EEWTKKLQ.** Peptide sequence H-Glu-Glu-Trp-Thr-Lys-Lys-Leu-Gln-COOH. MW = 1061.2 g/mol.

**LSALEE.** Peptide sequence H-Leu-Ser-Ala-Leu-Glu-Glu-COOH. MW = 660.7 g/mol.

**In vitro pepsin digestions of parent 23-residue peptide.** To determine what peptide fragments would be produced by pepsin digestion, the parent peptide (0.5 mg/mL peptide, prepared in 10% acetic acid) was incubated with pepsin (P-6887 from Sigma) at 37 °C in 10% acetic acid (pH 2.1). Three different incubations were carried out to obtain less extensive and more extensive digests of the peptides: 0.5 U pepsin for 160 min, 1500 U pepsin for 3 min, and 1500 U pepsin for 160 min. Aliquots of the reactions were quenched at the desired times by addition of pepstatin (10 µg/mL) and dilution with acetonitrile. Samples were analyzed by HPLC-MS and product fragments identified manually based on UV absorbance and mass.

**Peptide counterion exchange.** The TFA counterion in LSALEE•TFA was exchanged with HCl by repeated lyophilization from aqueous HCl solutions [25]. Peptide LSALEE•TFA (340 mg) was dissolved in 30 mL of 100 mM HCl solution and lyophilized overnight. The process was repeated twice. Complete removal of TFA from the peptide was confirmed using $^{19}$F-NMR analysis.

**19F-NMR.** Spectra were recorded on a Bruker AV Neo 399 spectrometer equipped with a 5 mm BBFO smart probe. A coaxial tube insert containing 500 μM trifluoromethoxybenzene (CAS 456-55-3) dissolved in d6-DMSO was used to provide a lock signal and internal concentration standard. For each spectrum, the pulse signal was modified to be centered between the outermost $^{19}$F peaks (for example, for spectra containing TFA at -78 ppm and trifluoromethoxybenzene at -58 ppm, the pulse was set to -68 ppm). The d1 relaxation time was increased to 8 seconds. Spectra were typically recorded with 64–90 scans.

**Animals.** The TSRI Institutional Animal Care and Use Committee approved all experimental protocols involving live animals. LDL receptor-deficient (LDLr$^{-/-}$) mice on a C57BL/6J background were initially purchased from Jackson Laboratories (Bar Harbor, ME) and were bred in house. The mice were weaned at 4 weeks of age and were fed ad libitum a standard chow diet (LabDiet 5053). Mouse cohorts selected for study were between 10 and 12 weeks old when studies were initiated. The high fat diet (HFD) used for LDLr$^{-/-}$ mice contained 1.25% cholesterol, 37.3% kcal from fat, and no cholate (Inotiv TD.94059).

ApoE deficient (ApoE$^{-/-}$) mice on a C57BL/6J background were initially purchased from Jackson Laboratories (Bar Harbor, ME) and were bred in house. The mice were weaned at 4 weeks of age and were fed ad libitum a standard chow diet (LabDiet 5053). Mouse cohorts selected for study were between 6 and 7 weeks old when studies were initiated. The high fat diet (HFD) used for ApoE$^{-/-}$ mice contained 0.2% cholesterol, 42% kcal from fat, and no cholate (Inotiv TD.88137).

10-week old C57BL/6J DIO mice (Jackson stock # 380050) or C57BL/6J DIO control mice (stock # 380056) were purchased from Jackson Laboratories (Bar Harbor, ME). These mice are fed a HFD (Research Diets # D12492i, 60% kcal from fat) or matched control diet (Research Diets # D12450Ji, 10% kcal from fat) starting at 6 weeks of age at Jackson Laboratories. The mice were maintained on the same diets at Scripps Research. The mice were kept at Scripps Research for two weeks prior to start of treatment to allow the animals to acclimate.

**Solutions for administration to mice.** For the majority of studies involving LDLr$^{-/-}$ or apoE$^{-/-}$ mice, PBS was used as vehicle. The pH was adjusted to 7 using 5 N NaOH in cases where the test agent (TFA, TFA analog, or short peptide) caused a drop in pH of the vehicle. Solutions containing TFA were prepared using sodium trifluoroacetate, not trifluoroacetic acid. Acetate buffer (10 mM) at pH 4 was used as vehicle for the study involving wild-type C57Bl mice. Stock solutions of test agent in vehicle were prepared fresh every one or two weeks, stored at -80°C, thawed, and filtered through a 0.22 μm sterile syringe filter before administration. Test agents were administered by daily i.p. injection (0.2 mL) or by daily oral gavage (0.2-0.25 mL), or in the drinking water for two or ten weeks. The concentration of TFA for i.p. or gavage varied from 1-60 mM to achieve the corresponding 10-600 μmol/kg dose. In the drinking water, a 1 mM TFA solution was used to achieved a dose of ~200 μmol/kg, because mice drank ~4.5 mL per day. The prepared drinking water solution was added to 50 mL graduated polypropylene tubes fitted with standard rubber stoppers and stainless-steel sippers of a type used routinely in mouse water bottles. The solution was changed daily. There was no significant difference in water or food consumption between groups. Fenofibrate was prepared fresh daily as a suspension, 20 mg in 2 mL of 0.5% methylcellulose (MC) and administered by oral gavage (0.2 mL) for a dose of 100 mg/kg. Feeding and treatment schedules for the in vivo efficacy studies are shown in Figure S1.

**Analysis of mouse plasma lipids and biomarkers.** Blood (200 μL) was collected after a 6 hour fast by retro-orbital puncture into a heparinized capillary tube and transferred to $K_2EDTA$ collection tube. Plasma was separated immediately by centrifugation of the blood samples at 5000 rpm for 10 minutes at 4°C and stored at -80°C. Plasma total cholesterol (TC) was measured using an enzymatic colorimetric method kit (Amplex® red cholesterol assay kit, No. A12216, Life Technologies) according to the manufacturer's instructions. Plasma triglyceride, SAA and CRP levels were measured by using a Triglyceride Colorimetric Assay Kit (Cayman, No. 10010303, Mouse SAA Mouse ELISA Kit (Life Technologies, No. KMA0021) and Mouse CRP ELISA kit (Invitrogen, No. EM20RB), respectively, following the manufacturer's instructions. qPCR for ACOX1 and ACADS genes in C57Bl mice was carried out using TaqMan™ Gene Expression

Assay (Catalog # 4331182), assay ID Mm01246834_m1 for ACOX1 and assay ID Mm00431617_m1 for ACADS.

**Evaluation of atherosclerosis.** Atherosclerotic lesion severity was assessed in the aortae as previously described [68]. At euthanasia, animals were perfused with PBS, followed by 4% formaldehyde (10% UltraPure EM Grade from Polysciences diluted in PBS, pH 7.2). For en face analysis, the entire mouse aorta was dissected from the proximal ascending aorta to the bifurcation of the iliac artery by using a dissecting microscope. Adventitial fat was removed, and the aorta was opened longitudinally, pinned flat onto black dissecting wax, stained with Sudan IV, and photographed at a fixed magnification. The photographs were digitized, and total aortic areas and lesion areas were calculated by using Adobe Photoshop version CS4, Chromatica V, and NIH Scion Image software (http://rsb.info.nih.gov/nih-image/Default.html). The results were reported as a percentage of the total aortic area that contained lesions.

As a second assessment of atherosclerosis, lesions of the aortic root (aortic sinus) were analyzed by a modification of current methods of valve lesion analysis [68]. Utilizing stereological principles, lesion volume was estimated across a fixed distance of the aortic sinus. After 10 minutes fixation in 4% paraformaldehyde during perfusion, hearts were cut at an angle perpendicular to the atria of the heart and embedded in OCT (Tissue-Tek). Frozen hearts were sectioned on a Leica cryostat, with 10-μm sections collected from the beginning of the aortic sinus (defined as when a valve leaflet became visible) to 500 μm below the beginning of the sinus. For hearts cut at an angle that resulted in valve leaflets not appearing in the same section (due to poor section angle), the lagging leaflet was used to determine the 500-μm distance. Sections were collected in duplicate at 50-μm intervals. Sections were stained with oil red O, counterstained with Gill hematoxylin 1 (Fischer Scientific International), photographed, and digitized for lesion analysis. Scoring of valve lesion areas was done for each of the 3 valve cusps individually. Lesion areas found only within the valve cusp were measured. Lesion volume estimation was determined from a 1 in 10 sampling rate; hence, valve cusps spaced at 140 μm were used to determine the lesion volume for a total of 4 sections analyzed per valve cusp. Lesion volume was calculated from an integration of the measured cross-sectional areas. Prediction of the coefficient of error (CE) in approximating lesion volume was computed using the Cavalieri estimator derived from a covariogram analysis of an ordered set of estimates of cross-sectional areas. This yielded CE values of less than 10% that were acceptable for a stereological computation of lesion volume.

**Harvesting of liver samples.** Euthanized mice were perfused with sterile 10 mL PBS (left ventricle to right atrium) to remove blood from the liver. The liver was excised and weight recorded. Sections of liver were cut and processed according to the various downstream processing needs. A small amount, approximately 30 mg, was placed into RNAlater (Invitrogen/Thermo Fisher Scientific # AM7021, Lithuania) and frozen at -80°C for RNA extraction. One of the liver lobes was cut into ~ 5 mm sections and placed into 3% paraformaldehyde in PBS. Sections of ~30 μg size were placed into lysing buffer (male mice) or flash frozen in liquid nitrogen (female mice) for extraction of PMP70 for western blotting. Sections of ~100 μg size were flash frozen in liquid nitrogen for analysis of palmitoyl-CoA oxidase activity.

**Confocal immunofluorescence imaging of liver sections.** Liver tissue was fixed in 3.5 mL freshly prepared 3% paraformaldehyde in PBS overnight at 4°C. Following fixation, the tissue was transferred to 3.5 mL 15% sucrose in PBS. When the tissue sank to the bottom of the tube, it was transferred to 3.5 mL 30% sucrose in PBS. Tissue samples were stored overnight at 4°C, providing ample time for the samples to sink to the bottom of the tube. Tissue samples were blotted on a kimwipe and transferred to a mold containing OCT (Tissue-Tek # 4583, Sakura Finetek USA, Torrance, CA) and frozen on a block of dry ice. Frozen samples were transferred to a -80°C freezer for storage. Samples were sectioned onto slides at 10-micron thickness using a Leica 1800 cryostat set at -20 °C, 2 sections per slide. Sectioned slides were stored at -20°C. Slides were washed once in 0.3 M glycine in PBS for 5 minutes and three times in PBS for 5 minutes at room temperature. Slides were permeabilized in 0.1% Triton X-100 in PBS for 5 minutes on ice. Secondary antibody binding sites were blocked in 5% normal goat serum + 0.1% Triton X-100 for one hour at room temperature. Slides were incubated with primary antibody (polyclonal rabbit anti-PMP70 # PS1650, Life Technologies, Carlsbad, CA) at final concentration of 2 μg/mL in PBS containing 5% normal goat serum. Primary antibody was applied at 100 μL per section, overlaid with a coverslip, and placed in a humidity chamber for three hours at room temperature. Coverslips were floated off the sections in PBS and slides were washed three times in PBS for 5 minutes. Secondary antibody (goat anti-rabbit AF 568 # A11036, Life Technologies, Carlsbad, CA) was applied at final concentration of 4 µg/ml in PBS containing 5% normal goat serum. Secondary antibody was also applied at 100 µL per section, overlaid with a coverslip, and placed in a humidity chamber for one hour at room temperature. Coverslips were floated off the sections in PBS and slides were washed three times in PBS for 5 minutes. Slides were washed once in 0.1% Triton X-100 in PBS for 5 minutes and three more times in PBS for 5 minutes then stained for 2 minutes with DAPI (Biolegend # 422801, San Diego, CA) at final concentration of 0.5 µg/mL. Slides were washed a final three times in PBS, mounted with Vectashield (Vector Laboratories # H-1400, Burlingame, CA) at 30 µL per section, covered with a coverslip (22x50x1.5 mm) and sealed with clear nail varnish. Sealed slides were stored at 4°C, covered from light. Images were collected on a Zeiss LSM 710 confocal microscope, using a 60x Plan-Apochromat NA 1.4 objective (oil). Single 60x Z-planes were imaged sequentially with the following settings: pixel dwell time of 2.33 microseconds, and pixel dimensions of 0.098 microns (1380 x 1380 pixels); DAPI- 2.0% laser power, emission filter 410-585nm, master gain 696, digital offset 7.0, pinhole 58um; green autofluorescence- 77.3% laser power, emission filter 493-574nm, master gain 790, digital offset 0.00, pinhole 51 microns; red fluorescence- 7.9% laser power, emission filter 568-712nm, master gain 790, digital offset 8.0, pinhole 51 microns.

**TFA pharmacokinetics in vivo.** Male LDLr$^{-/-}$ mice (~25 g) were maintained on a chow diet. Sodium trifluoroacetate was formulated as a 25 mM solution in pH 7 PBS, and sterile filtered through a 0.2 µm filter. Groups of two mice received a 200 µmol/kg dose of TFA via intraperitoneal injection (0.2 mL) or oral gavage (0.2 mL). Three such groups were used so that no animal had more than two survival blood draws. 100 or 200 µL blood was drawn from the retro-orbital sinus into a heparinized capillary tube at different intervals beginning 15 min after dosing. Final blood collection (~500 µL) was done by cardiac puncture. Plasma was isolated immediately from the whole blood by centrifugation at 5000 rpm for 10 minutes at 4°C. Immediately after blood collection and plasma separation, plasma samples were flash frozen in liquid nitrogen and stored at -80 ºC until analysis. TFA concentrations in the plasma samples were determined by using $^{19}$F-NMR, with a lower limit of quantification of 10 µM. Plasma samples were thawed and diluted with deionized water to a final volume of 0.5 mL. Spectra were recorded using a coaxial tube insert containing 500 µM trifluoromethoxybenzene (CAS 456-55-3) dissolved in d6-DMSO to provide a lock signal and internal concentration standard. A standard curve for TFA in plasma was generated by spiking known concentrations of TFA into plasma from untreated animals. TFA concentrations were calculated by comparing the NMR integral for TFA with that of the internal concentration standard.

**Stability of TFA analogs in vivo.** Male LDLr$^{-/-}$ mice (~25 g) were maintained on a chow diet. The TFA analog (trifluoroacetamide, N-ethyl-trifluoroacetamide, 2,2-difluorobutyric acid, or difluoroacetic acid) was formulated as a 25 mM solution in PBS, with the pH adjusted to ~7 using 5 M NaOH if necessary, and sterile filtered through a 0.2 µm filter. Groups of four mice received a 200 µmol/kg dose of the analog via oral gavage (0.2 mL). One hour following administration, the mice were sacrificed and blood collected by cardiac puncture (~500 µL). Plasma was isolated immediately from the whole blood by centrifugation at 5000 rpm for 10 minutes at 4°C. Immediately after blood collection and plasma separation, plasma samples were flash frozen in liquid nitrogen and stored at -80 ºC until analysis by using $^{19}$F-NMR. Plasma samples were thawed and diluted with deionized water to a final volume of 0.5 mL. Spectra were recorded using a coaxial tube insert containing 500 µM trifluoromethoxybenzene (CAS 456-55-3) dissolved in d6-DMSO to provide a lock signal and internal concentration standard. Spiking of the plasma with TFA or the TFA analogs was used to confirm the identity of the observed peaks.

**RNA-seq sample preparation and sequencing.** Mouse liver tissue that had been stored in RNAlater at -80°C after harvest (around 20–30 mg) was used for RNA extraction with an RNeasy Mini Kit (Qiagen) following the manufacturer's instructions. RNA quality was assessed by Agilent's Bioanalyzer or Tapestation (Agilent Technologies). RNA quantity was measured by Qubit RNA BR/HS kit (Invitrogen). The rRNA was removed from total RNA with Ribo-Zero Plus rRNA Depletion Kit (Illumina). RNA-seq libraries were constructed using NEBNext Ultra II RNA Library Prep Kit for Illumina (NEB). Samples were purified with NEBNext® Sample Purification Beads (NEB) and quantified by Qubit dsDNA HS Assay Kit (Invitrogen). NEBNext Multiplex Oligos for Illumina (Dual Index Primer Set1) (NEB) was used to barcode libraries. Libraries with different barcodes were pooled, and paired-end sequencing (150 bp) was performed on an Illumina NextSeq platform at the Scripps Research Institute next-generation sequencing core.

**RNA-seq analysis.** RNA-Seq data were processed and aligned using the NF-core RNAseq v3.5 pipeline [69]. Reads were trimmed using Trim Galore (v0.6.7) and aligned using Star (v2.6.1d) against the GRCm38 reference genome and annotation. Transcript quantification was performed using Salmon (v1.5.2). Differential analysis was performed using DESeq2 (v1.28.0) and the R statistical environment (v4.0.2). Pathway analysis was performed using pathfindR (v1.6.3) [70] inside the R statistical environment (v4.0.2). The mouse KEGG gene set was used to determine the active subnetworks from the protein-protein interaction dataset. The false discovery rate (FDR) was used as the p-value correction methodology ($p<0.05$). Our RNA-seq data is deposited in Bioproject at accession #: PRJNA917696**.**

**Meta-transcriptomics sequencing and analysis.** In an anaerobic chamber, cecal material from HFD-fed LDLr$^{-/-}$ mice was resuspended in Chopped Meat Carbohydrate Broth (BD, 297307) at a ratio of 1.5 ml per 100 mg of cecal content. The mixture was vortexed for 5 minutes and then allowed to settle and strained through a 100 μm filter. The resulting supernatant was treated with 1 mM TFA or vehicle control and then incubated anaerobically at 37°C for 6 hours. The bacteria were collected by centrifugation and stored in PowerProtect DNA/RNA solution (Qiagen, Cat. 14800) at -80°C. Total RNA was extracted using the RNeasy PowerFecal Pro Kit (Qiagen, Cat. 78404) according to the manufacturer's instructions. The RNA quality was evaluated using the TapeStation (Agilent Technologies), and the quantity was determined with the Qubit RNA BR Kit (Invitrogen). Library preparation and sequencing were subsequently carried out by Novogene (Sacramento, CA). The SAMSA2 metatranscriptome analysis pipeline was used for organism and gene function identification.

**16S rRNA gene sequencing and analysis.** 16S rRNA sequencing was conducted following a previously established method [44] with minor modifications. In short, in an anaerobic chamber, cecal contents from mice fed HFD or CHOW diets were resuspended in PBS (1.5 ml per 100 mg of cecal content). The mixture was vortexed for 5 minutes, allowed to settle, and strained through a 100 μm filter. Supernatant was diluted 12.5-fold with Chopped Meat Carbohydrate Broth (Anaerobe Systems). TFA or a control gut microbiome remodeling cyclic peptide was added at the desired concentration, and the mixture was incubated anaerobically at 37°C overnight. Bacterial cells were then collected, and DNA was extracted using the DNeasy UltraClean Microbial Kit (Qiagen) according to the manufacturer's instructions. The V3-V4 region of the 16S rRNA gene was amplified using KAPA HiFi HotStart ReadyMix (Kapa Biosystems) with the following primers: forward primer, (5′-AATGATACGGCGACCACCGAGATCTACACNNNNNNNNACAC TCTTTCCCTACACGACGCTCTTCCGATCTACTCCTACGGGAGGCAGCAG-3′) and reverse primer, (5'-CAAGCAGAAGACGGCATACGAGATNNNNNNNNGTGACTGGAGTTCAGACGTGTGCTCTTCCGATCT nnnnnnnnnnGGACTACHVGGGTWTCTAAT-3'). Each sample was tagged with a unique 8-base barcode (N) for identification and a 10-base unique molecular identifier (n) to correct for PCR duplication artifacts during library preparation. Amplicons were quantified using the Qubit dsDNA Assay Kit (Invitrogen), and equimolar amounts of DNA from each sample were pooled and purified with Agencourt AMPure XP beads (Beckman Coulter). The final library was sequenced on an Illumina NextSeq platform. Sequencing data were demultiplexed and analyzed for taxonomic identification using QIIME2 pipeline.

**Cell culture.** Rat hepatoma FAO cells (obtained from Sigma-Aldrich) were cultured in Ham's F-12K (Kaighn's) medium (Gibco) containing 10% FBS (HyClone), 100 U/ml penicillin, and 100 μg/ml streptomycin. The HepG2 cell line (obtained from ATCC) was cultured in DMEM medium (Gibco) containing 10% FBS (HyClone), 100 U/ml penicillin, and 100 μg/ml streptomycin. Cells were maintained at 37 °C in a humidified atmosphere of 95% air and 5% $CO_2$.

**Luciferase reporter assay.** We followed a previously reported procedure [31] with some modifications. Plasmid PPRE X3-TK-luc, which encodes a firefly luciferase gene under the control of peroxisome proliferator response element (PPRE), was used to monitor the effect of TFA on different PPAR isoforms. Beta-galactosidase was used to normalize the luciferase signal by co-transfection with pSV-β-galactosidase control vector (Promega). All three PPAR isoform cDNAs were cloned into the HindIII site of pBApo-CMV plasmid (Takara) by NEBuilder® HiFi DNA Assembly kit (NEB). The primers used are listed in **Table S1**. All constructed plasmids were verified by Sanger sequencing (Eton bioscience). To carry out the luciferase reporter assay, HepG2 cells were seeded into a 24-well plate at ~$2.0 \times 10^5$ cells per well in 0.5 mL DMEM medium (Gibco) containing 10% FBS (HyClone), 100 U/ml penicillin, and 100 μg/ml streptomycin and grown overnight. Next, 400 ng PPRE X3-TK-luc plasmid, 100 ng pSV-β-galactosidase control vector, and 20 ng pBApo-CMV containing one of the PPAR isoforms were added to the culture media for transfection with lipofectamine 3000 (Invitrogen), according to manufacturer's protocol. After 24 h, the medium was replaced by fresh DMEM containing 10% FBS with different concentrations of TFA or positive control compound and incubated for the desired time. The cells were washed with PBS, and then were scratched and lysed into 150 µl reporter lysis buffer (Promega). Luciferase activity in the lysate was measured using the Bright-Glo™ Luciferase Assay System (Promega) and normalized by β-galactosidase activity using β-Galactosidase Enzyme Assay System (Promega).

**FAO cell treatment and RT-qPCR analysis.** Rat hepatoma FAO cells were seeded in a 24-well plate at ~2.0 × $10^5$ cells per well in Ham's F-12K (Kaighn's) medium (Gibco) containing 0.5% FBS (HyClone), 100U/ml penicillin, and 100 µg/ml streptomycin and grown overnight. The media was then replaced with fresh media containing 1 mM TFA, 10 mM TFA, or 0.1 mM ciprofibrate. The treated cells were cultured for 24 h before RNA isolation. For total RNA isolation, the supernatant was discarded, and the cells were washed with PBS. Total RNA was isolated using Trizol reagent (Invitrogen) following the manufacturer's protocol. Reverse transcription was carried out using QuantiTech Reverse Transcription Kit (Qiagen). The cDNA was diluted 10-fold and used for quantitative PCR with PowerUp™ SYBR™ Green Master Mix (Applied Biosystems) on a Bio-Rad CFX384 Touch Real-Time PCR Detection System, using primers specific for each gene (**Table S1**). The ΔΔCt method was used for quantification. Data were normalized to housekeeping gene glyceraldehyde-3-phosphate dehydrogenase (GAPDH).

**CRISPR-Cas9 gene knockout of PPAR-α.** To easily screen mutants, a fragment deletion strategy was used instead of indel mutation. To construct the FAO PPAR-$\alpha^{-/-}$ cell line, two gRNAs were designed using CRISPOR website [71] and inserted into the pSpCas9(BB)-2A-Puro (PX459) plasmid by Golden gate method. The constructed plasmids were verified by Sanger sequencing. The gene knockout protocol was modified from the previously described method [72]. Briefly, FAO cells were seeded into a 24-well plate at ~5.0 × $10^4$ cells per well and transfected with the constructed plasmid by lipofectamine 3000 (Invitrogen). After 6 hours, the transfection reagents were removed by replacing the media with fresh medium. The transfected cells were incubated for 48 hours to express the puromycin resistant protein. Cells were harvested and resuspend in fresh medium containing 20 µg/ml puromycin (Invitrogen) and seeded into a 60 mm petri dish. After 3 days, the medium was replaced with fresh medium containing no puromycin. A dilution-based method was used to isolate a single clonal cell line which had the designed mutation. The expected deletion mutant of the PPAR-α gene was confirmed by PCR.

**Western-blot analysis.** Protein fractions from fresh or flash-frozen tissue were extracted using T-PER tissue protein extraction reagent (Thermo Fisher Scientific) according to the manufacturer's protocol. Around 25 mg of liver tissue was homogenized in 500 µL extraction reagent containing 1 x Halt™ protease and phosphatase inhibitor cocktail (Thermo Fisher Scientific) using a pellet pestles cordless motor. After homogenization, samples were centrifuged at 10,000×g for 5 min at 4°C to pellet tissue debris. The supernatant was filtered using a 0.22 µm filter, and the total protein concentration was measured by using a BCA protein assay kit (Thermo Fisher Scientific). SDS-PAGE was carried out using a Bolt 4–12% gradient Tris-Bis gel following the manufacturer's protocol. 25-50 µg total protein was loaded on each well and run at 100 V for 100 min. PVDF membrane (Thermo Fisher Scientific) was used for transfer at 30 V for 2 h by wet transfer. After transfer, the membrane was blocked in 5% BSA for 1 h at room temperature. Immunoblotting was carried out using Pierce Fast Western Blot Kit and SuperSignal West Pico Substrate (Thermo Fisher Scientific), following the manufacturer's protocol. Briefly, the rabbit anti-PMP70 antibody (Invitrogen) and anti-GAPDH antibody (Invitrogen) were diluted 1:1000 in the diluent buffer and incubated with the PVDF blot at room temperature for 1 h. After washing, the PVDF blot was incubated with HRP-conjugated antibody at RT for 10 min. Excess secondary antibody was removed using three washed with washing buffer. SuperSignal West Pico Substrate was used to detect the HRP-conjugated secondary antibody. Imaging was conducted via Bio-Rad ChemiDoc XRS+ Imaging system.

**Palmitoyl-CoA oxidation assay.** Around 150 mg liver tissue that had been perfused with PBS was homogenized in 750 µL lysis buffer containing 50 mM Tris-HCl (pH 8.0), 152 mM KCl, 0.1% triton X-100 and 1 x Halt™ protease and phosphatase inhibitor cocktail (Thermo Fisher Scientific). Homogenization was performed using a pellet pestles cordless motor on ice. Homogenates were centrifuged at 2500g for 15 min at 4 °C. The supernatant was collected, and protein concentration was determined using a BCA protein assay kit (Thermo Fisher Scientific). Supernatant was stored at -80 °C until analysis. Palmitoyl-CoA oxidation activity was determined based on a previous method [31] with slight modification. Briefly, around 100 µg protein was preincubated in 1 ml reaction mixture containing 50 mM Tris-HCl (pH 8.0), 0.2 mM $NAD^+$, 0.01 mM FAD, 0.1 mM coenzyme A, 6.25 mM dithiothreitol, 0.15 mg/ml bovine serum albumin, 1.5 mM KCN and 0.01 % (w/v) triton X-100 for 5 min in a cuvette at 37°C in a Cary 100 UV-Vis spectrophotometer. Palmitoyl-CoA oxidation was initiated by adding 0.05 mM palmitoyl-CoA substrate (Sigma Aldrich, cat # P9716). The absorbance at 340 nm was recorded for 15 min at 37 °C. The rate of NADH formation was calculated using the initial linear region of the curve (first 5 min) and converted to NADH concentration units using an extinction coefficient of 6.22 $mM^{-1}$ $cm^{-1}$.

**Biochemical assays for formation of acetyl-CoA and trifluoroacetyl-CoA.** Reactions were carried out in 50 µL buffer containing 100 mM Tris-HCl (pH 8.0), 5 mM ATP, 5 mM $MgCl_2$, 2 mM CoA, 1 mM DTT, and 0.15% Triton X-100. Sodium acetate and sodium trifluoroacetate were both added at a final concentration of 10 mM. Acetyl-CoA synthetase (Sigma Aldrich, A1765) was used at 5 µM. For liver tissue lysate reactions, 70 µg of total protein was added to the 50 µL mixture. To generate ciprofibroyl-CoA, 2 mM ciprofibrate (TCI America) and 50 µg of rat microsomes (Gibco) were included. After incubating at 37°C for 1 hour, the reactions were quenched by adding 200 µL of chloroform/methanol (2:1, v/v) followed by vigorous mixing. After centrifugation, the upper phase was directly used for LC-MS analysis. Each sample (10 µL) was injected onto an Agilent Poreshell 120 EC-C8 (4.6 x 50 mm, 2.7 µm) analytical column using the Agilent InfinityLab LC/MSD system (Agilent Technologies). Buffer A (0.1% formic acid in water) and buffer B (0.1% formic acid in MeCN) were used in a gradient (flow rate 0.5 mL/min) of 5 % buffer B for 1 min, followed by a linear increase to 100 % buffer B over 9 min, 100 % buffer B for 2 min, followed by descent to 5% buffer B.

**Targeted bile acid metabolomics.** Bile acids were extracted from plasma or feces samples according to published procedures [73]. Briefly, fecal samples were lyophilized and homogenized. Five milligrams of fecal powder were extracted with 250 µL of methanol containing heavy internal standards. Fifty microliters of plasma were extracted with 150 µL of methanol containing heavy internal standards. After vortexing for 10 minutes and centrifuging (16,000g, 4°C, 10 min), supernatants were transferred to glass vials for injection and analysis by LCMS. Bile acids were analyzed on a Dionex Ultimate 3000 LC system (Thermo) coupled to a TSQ Quantiva mass spectrometer (Thermo) fitted with a Kinetex C18 reversed phase column (2.6 µm, 150 x 2.1 mm i.d., Phenomenex). The following LC solvents were used: solution A, 0.1% formic acid and 20 mM ammonium acetate in water, solution B, acetonitrile/methanol (3/1, v/v) containing 0.1% formic acid and 20 mM ammonium acetate. The following reversed phase gradient was utilized: at a flow rate of 0.2 mL/min with a gradient consisting of 25-29% B in 1 min, 29-33% B in 14 min, 33-70% B in 15 min, up to 100% B in 1 min, 100% B for 9 min and re-equilibrated to 25% B for 10 min, for a total run time of 50 min. The injection volume for all samples was 10 µL, the column oven temperature was set to 50°C and the autosampler kept at 4°C. MS analyses were performed using electrospray ionization in positive and negative ion modes, with spray voltages of 3.5 and -3 kV, respectively, ion transfer tube temperature of 325°C, and vaporizer temperature of 275°C. Multiple reaction monitoring (MRM) was performed by using mass transitions between specific parent ions into corresponding fragment ions for each analyte.

**Untargeted metabolomics.** The untargeted metabolomics study was carried out by the Scripps Research Center for Metabolomics and Mass Spectrometry core. Plasma samples (100 µL) from female $LDLr^{-/-}$ mice treated by daily oral gavage for two weeks with a 200 µmol/kg TFA daily dose (n=10) or PBS vehicle (n=10) were extracted with cold MeOH (400 µL) and analyzed individually by reversed phase LCMS, in both positive and negative mode. MS data was analyzed (peak alignment, integration, annotation, statistics) using XCMSOnline (www.xcmsonline.scripps.edu) to run statistical comparison between the sample groups. The following cutoff thresholds were used for analysis: a fold-change >1.5, p-value < 0.01, and maximum peak intensity >5000. The XCMSOnline output for positive and negative mode analyses is provided in Spreadsheet S2. The mass fragmentation spectra were manually searched for the presence of potential trifluoroacetyl fragment ions (97.0 Da for $F_3CCO$ and 69.0 Da for $F_3C$ fragments). The lists of metabolite M/Z from XCMSOnline were also searched manually for pairs of parent ion masses that differed by the mass of a trifluoroacetyl modification (96.0 Da for $+F_3CCO-H$). The raw metabolomics data for this study is available for download from the National Metabolomics Data Repository with the Study ID ST003595 at http://dx.doi.org/10.21228/M88R7M.

**Preparation of samples for proteomics.** Samples of mouse liver homogenate (100 µl, 2 mg protein / ml) were combined with urea (48 mg) in Protein LoBind Tubes (2.0 ml, Eppendorf 022431102). To each sample was added a solution of tris(2-carboxyethyl)phosphine hydrochloride and potassium carbonate (50 µl, 100 mM and 300 mM respectively, in DPBS), and the samples were incubated at 37°C for 30 min with shaking. A solution of iodoacetamide (70 µl, 400 mM in DPBS) was then added and the samples were incubated at ambient temperature for a further 30 min in the absence of light. Cold chloroform (220 µl), methanol (880 µl), and water (440 µl) were then added to each sample, and the mixtures were briefly vortexed followed by centrifugation (10,000 g, 10 min). The upper liquid phases were carefully removed, and cold methanol (600 µl) was added to each sample. The samples were homogenized in a water bath sonicator, centrifuged (10,000 g, 10 min), and the supernatants were removed. To each sample was added triethylammonium bicarbonate solution (TEAB, 160 µl, 100 mM, pH 8.5) and the protein residue was resuspended by probe sonication (1 s, 7 pulses, 15 % amplitude, 15 ms on, 40 ms off). A solution of LysC (20 µl, 50 µg / ml in TEAB, New England BioLabs P8109S) was added to each sample. Samples were then incubated for 2 h at 37°C with shaking. Solutions of trypsin (20 µl, 100 µg / ml in TEAB, Promega V5111), Protease Max (2 µl, 1 % w/v in TEAB, Promega V2072), and calcium chloride (2 µl, 100 mM) were added, and the mixtures were incubated overnight at 37°C with shaking. The samples were centrifuged (12,000 g, 10 min) and peptide concentrations were estimated using a Pierce Quantitative Fluorometric Peptide Assay according to the manufacturer's instructions (Thermo 23290). Volumes corresponding to 25 µg digested protein were transferred to new Protein LoBind Tubes (1.5 ml, Eppendorf 022431081), and each sample was made up to 35 µl with TEAB followed by the addition of acetonitrile (12.5 µl). The peptides were labeled with a TMT10plex™ Isobaric Label Reagent Set (Thermo 90406) according to the manufacturer's instructions. The reactions were quenched through the addition of hydroxylamine solution (3 µl, 5 % *w/w* in water) and incubated for 15 min before the addition of formic acid (5 µl). Solvents were then removed from the samples by vacuum centrifugation. Each residue was dissolved in trifluoroacetic acid solution (40 µl, 0.1 % *v/v*), and the solutions were combined together. An additional volume of trifluoroacetic acid solution (200 µl, 0.1 % *v/v*) was used to wash any residual peptide solutions into the combined solution (final volume 600 µl). The combined peptide solution was eluted into eighteen fractions using a Pierce High pH Reversed-Phase Peptide Fractionation Kit (Thermo 84868) according to the manufacturer's instructions. The first fraction was combined with the tenth, the second with the eleventh, etc. to yield nine solutions which were then dried by vacuum centrifugation. The residues were redissolved in formic acid solution (65 µl, 0.1 % in water), centrifuged (10,000 g, 10 min), and the supernatants transferred to LC-MS vials for analysis.

**Mass spectrometric proteomic analysis.** Proteomic samples were analyzed as described previously [74]. Briefly, from each sample 10 µl was loaded on to a precolumn (Acclaim PepMap 100, 75 µm × 2 mm) and separated using an Acclaim PepMap RSLC analytical column (75 µm × 15 cm) on an UltiMate 3000 RSLCnano system (Thermo Fisher Scientific). The following gradient was applied at a flow rate of 300 µl/min and column temperature of 35°C: 2% buffer B for 10 min, rising to 30% buffer B over 192 min, followed by 60 % buffer B for 5 min, 60 % rising to 95 % buffer B over 1 min, 95% buffer B for 5 min, falling to 2% buffer B over 1 min, and held at 2 % buffer B for 6 min (total 220 min, buffer A is 0.1 % *v/v* formic acid in water, buffer B is 0.1 % *v/v* formic acid in acetonitrile). The eluted peptides were analyzed with a Thermo Fisher Scientific Orbitrap Fusion Lumos mass spectrometer. The spectrometer cycle time was 3 s and a nano-LC electrospray ionization source was used with an applied voltage of 2,000 V. The data were collected using Xcalibur (v.4.1.50). The MS1 scan range was set to 375 to 1,500 *m/z*, with a maximum injection time of 50 ms (dynamic exclusion enabled, repeat count 1, duration 20 s). The resolution was set to 120,000 with an automatic gain control value of $1 \times 10^6$ ions. The peptides selected for MS2 analysis were isolated with the quadrupole (isolation window 1.6 *m/z*) and fragmented using collision-induced dissociation (30 % collision energy), and the resultant fragments were detected using the ion trap (automatic gain control $1.8 \times 10^4$, maximum injection time 120 ms). The MS3 spectra were generated through high-energy collision-induced dissociation (65 % collision energy). Synchronous precursor selection was used to isolate up to 10 MS2 ions for MS3-based quantitation.

**Proteomic data analysis.** Proteomic analysis was performed with using the SEQUEST HT algorithm in the Proteome Discoverer 3.0 software package (Thermo Fisher Scientific). Fragment tolerances were set to 0.6 Da, and precursor mass tolerances set to 10 ppm with one missed cleavage site allowed. Carbamidomethyl (C, +57.02146) and TMT-tag (K and N-terminal, +229.1629) were specified as static modifications while oxidation (M, +15.994915) and optionally trifluoroacetate (K, +95.9823) were specified as variable modifications. Spectra were searched against the Mus musculus protein database (UniProt, 17,158 sequences) using a false discovery rate of 1 % (Percolator). MS3 peptide quantitation was performed with a mass tolerance of 20 ppm. Protein abundances were normalized to the median abundance of each channel. Identified proteins were required to have at least two unique peptides. TMT ratios obtained by Proteome Discoverer were $\log_2$-transformed, and p-values were calculated via Student's two-tailed t-tests with three biological replicates. The mass spectrometry proteomics data have been deposited to the ProteomeXchange Consortium via the PRIDE partner repository [75] with the dataset identifier PXD057035. Open search was performed using the MSFragger (4.0) and Philosopher (5.1.0) software packages running under FragPipe (20.0), using the 'Open' workflow with mass range of -150 to 500 Da [76-79].

**Table S1. Primers used in this study.**

| Primers | Sequence (5'to 3') |
|---|---|
| *For RT-qPCR* | |
| fEHHADH-F | TCAGTTGGCGTTCTTGGCTT |
| fEHHADH-R | GAAGCTTGGCCGTTCTGATG |
| fGAPDH-2ndF | GCCTGGAGAAACCTGCCAA |
| fGAPDH-2ndR | CACAGGAGACAACCTGGTCC |
| fACOX1-2ndF | CGTGCAGCCAGATTGGTAGA |
| fACOX1-2ndR | AACGACCACGTAGTGGCAAT |
| fACAA1-F | TCTACCACGGCTGGAAACTC |
| fACAA1-R | ATAGGACCTCAGGACGCCAA |
| *For PPAR isoforms construction* | |
| CMVmPPAR-α-F | GTCGACCTGCAGGCATGCAAGCTTGCCACCATGGTGGACACAGAGA |
| CMVmPPAR-α-R | TTTCAGTTAGCCTCCCCCAAGCTTCCATCTCAGGAAAGATCA |
| CMVmPPAR-δ/β-F | GTCGACCTGCAGGCATGCAAGCTTGCCACCATGGAACAGCCACAGGAG |
| CMVmPPAR-δ/β-R | TTTCAGTTAGCCTCCCCCAAGCTTGGCTGCGGCCTTAGTACA |
| CMVmPPAR-γ-F | GTCGACCTGCAGGCATGCAAGCTTGCCACCATGGTTGACACAGAGATG |
| CMVmPPAR-γ-R | TTTCAGTTAGCCTCCCCCAAGCTTGGGTGGGACTTTCCTGCTAA |
| *For CRISPR-Cas9 knock-out* | |
| fPPAR-α-gRNA1-F | CACCGACATCGAGTGTCGAATATG |
| fPPAR-α-gRNA1-R | AAACCATATTCGACACTCGATGTC |
| fPPAR-α-gRNA2-F | CACCGCTGTGGAGGTCCCATAATA |
| fPPAR-α-gRNA2-R | AAACTATTATGGGACCTCCACAGC |
| fPPAR-αVeriF | GCATGTATCACGACCCCTGT |
| fPPAR-αVeriR | TACCTCACCCTTCTCCCGAG |
